# Investigating the landscape of plant-pollinator interactions in a hybrid zone

**DOI:** 10.64898/2026.02.25.708067

**Authors:** N.J. Engle-Wrye, O.M. Carril, C.G. Mohottige, T.E. Mlsna, R.A. Folk

## Abstract

Little is known about environmental drivers of opportunities for hybridization, but its phylogenetic distribution across species and areas is heterogeneous, suggesting that ecological traits may play an important role in concert with postzygotic isolation. Because plant-pollinator interactions are responsible for gene flow in most plant species, differences in the mosaic landscape of plant-pollinator interactions could explain why some plants are particularly prone to hybridization. Prezygotic isolation is mediated by sometimes complex pollen presentation; conversely, conserved pollination strategies would lead to evolutionary constraints on pollinator assemblage divergence in the speciation process and therefore predict higher opportunities for gene flow, although this hypothesis has yet to be tested. The plant taxonomic tribe Heuchereae (Saxifragaceae) is a well-characterized system for pollinator interactions and particularly for floral scent, the primary pollinator attractant in the group. Floral volatile organic compounds (VOCs) in this clade are hypervariable at the population level and are thought to be responsible for pollination selectivity, leading to divergent pollinator assemblages. Observing a contrast of hybridizing and non-hybridizing species, the levels of attractant divergence may therefore predict levels of hybridization.

We investigated pollination biology in the plant genus *Heuchera*, notable for frequent interspecific gene flow compared to tribal relatives, asking whether high rates of hybridization may be associated with low interspecific divergence of VOCs and the pollinator assemblages they shape, using as our system the hybrid zone between *H. americana* var. *americana* and *H. richardsonii* in the midwestern USA. We optimized a closed-space collection and GC-MS (gas chromatography-mass spectrometry) protocol to characterize VOCs in *Heuchera* flowers. To identify floral visitation and effective pollinators we conducted pollination observations at 40 *Heuchera* populations over the span of two field seasons. GC-MS data from 89 *Heuchera* specimens representing 69 populations suggests that classes of VOCs, and to a large extent individual compounds, are shared within the hybrid complex while other *Heuchera* that are not thought to hybridize with these species have distinct species-specific compounds. Pollination observations and metabarcoding of pollinator pollen loads confirm shared effective pollinators in the hybrid zone and between adjacent parental populations. Attractant and visitation data considered together suggest that conservatism of pollinator interactions may be a typical feature associated with frequent hybridizers, perhaps arising from developmental or biochemical constraints on prezygotic isolation, and more broadly that the macroevolution of isolation mechanisms may be predictive of natural hybridization rate.

## Introduction

Hybridization, or the flow of genetic information between varieties or species, has a major influence on evolution by creating novel genotypes, changing allele frequencies in populations, and possibly even contributing to the diversification process (Rieseberg, 2006; Mallet, 2007; Soltis and Soltis, 2009; Arnold et al., 2012; Abbott et al., 2013; Mitchell et al., 2019; Turchetto et al., 2022) by generating new species or by causing extirpation through homogenization (Rhymer and Simberloff, 1996; Wolf et al., 2001; Buggs and Pannell, 2006; Todesco et al., 2016; Campbell et al., 2019; Turchetto et al., 2022). The rate at which hybridization occurs is heterogenous across the tree of life (Dowling and Secor, 1997; Mallet, 2005; Baack and Rieseberg, 2007; Whitney et al., 2010), being apparently absent in many sexual organisms, and very common in others, like many plants. This basic observation about the natural history of hybridization has been discussed for two centuries (Kölreuter, 1766; Mendel, 1866; Goulet et al., 2017), but the driving factors behind the differing prevalence and evolutionary significance of hybridization in different taxa remains poorly understood. A statement as commonplace as “plants hybridize frequently” cannot be easily explained with a general mechanism, although it is obvious that this is not happenstance and that aspect of the plant body or lifestyle generally enable it, as could be said for other taxa.

Evolutionary constraints on hybridization (i.e., isolation mechanisms, which we shall call “constraints” to signal our interest in their absence) can be conceptualized in two categories: intrinsic and extrinsic. Intrinsic constraints, those inherent to the organism in isolation, such as intersterility, are intensively studied (Sobel et al., 2010; Christie and Strauss, 2018, 2019) and would form a straightforward explanation of hybridization frequency insofar as these mechanisms differ among taxa. More perplexing is the situation of taxa that generally lack strong genetic isolation mechanisms yet seem to maintain “good” species, yet this is what is commonly observed in plant genera. Orchids are illustrative of the importance of extrinsic drivers, those rooted in organismal context: orchid intergeneric and interspecific hybridization is often easily induced in the greenhouse and the basis of thousands of commercial cultivars, but is less commonly observed in the wild than prolific horticultural hybrids would suggest (Johnson, 2018); many similar examples can be produced from familiar landscaping plants (Soltis and Soltis, 2009; Whitney et al., 2010). Wagner (1970) responded in part to the distorted depiction of evolution yielded by greenhouse crossing experiments, famously declaring hybridization and polyploidy largely an intellectual distraction from the main line of evolution. The prominence of this “biosystematic noise” in the phylogenetic picture has been generally enriched in the following half century, but differences in the picture of hybridization as painted by field and greenhouse studies still suggests that the simple occurrence and prevalence of strong genetic isolation mechanisms alone poorly predicts which taxa are liable to hybridize.

Extrinsic drivers can generally be further divided into abiotic and biotic components. Some of the most prevalently evoked ideas about drivers of hybridization are focused on abiotic variables such as climate change, disturbances, or geological factors, and how these impact geographic distribution; this idea has a long history of work (Rosendahl et al., 1936; Anderson and Stebbins, 1954; Stebbins, 1959; Abbott and Rieseberg, 2012; Chunco, 2014; Folk et al., 2016; Vallejo-Marín and Hiscock, 2016; Folk et al., 2018b, 2023). Clearly, overlapping or adjacent environmental niche clearly is important in creating geographic opportunities to exchange genes, normally a dispersal-limited process; reinforcement acts against this (Servedio and Noor, 2003) but frequent change in sympatry over time would create instability in the landscape of interactions that would then undermine reinforcement (Butlin, 1987; Servedio, 2000; Coyne and Orr, 2004). One classical hypothesis about an abiotic driver of hybridization is that of Pleistocene glaciation (Anderson and Stebbins, 1954) whereby historical climate change pushed previously discontinuous populations into contact, thereby leading to novel gene flow opportunities. This Pleistocene glaciation hypothesis is one of the most frequently cited in plants (Folk et al., 2018b; Turchetto et al., 2022; Folk et al., 2023). Subsequently, post-Pleistocene warming and aridification can be evoked as a mechanism that can lead to the cessation of range overlaps, thereby ceasing opportunities for gene flow and rendering gene flow as a historical process. While climatic factors are most frequently cited as abiotic drivers of hybridization opportunities, other abiotic factors disposed to lability over time are similarly suitable candidates, such as disturbance regimes like fire or volatile landscape connectivities like land bridges.

Biotic interactions are less commonly assessed in hybridizing groups (Anderson and Stebbins, 1954; Whitney et al., 2010; Goulet et al., 2017), but remain as an important class of factors for plants, which usually rely on other species to mediate gene flow. Conservatism of biotic niche (Wiens et al., 2010), that is, the tendency of closely related species to share ecological interactions with other organisms, could explain why some plants appear to be inherently prone to hybridization. Most flowering plants depend on pollinators to mediate gene flow (Tepedino, 1979; Kearns and Inouye, 1997), and many taxa have elaborate mechanisms to filter available pollinators to those that are most effective in moving pollen and to target conspecifics (Johnson and Steiner, 2000; Fenster et al., 2004). Bilaterally symmetric, highly derived flowers like orchids use elaborate shape, chemistries, and even mechanical contraptions within the flower to exquisitely manage pollen deposition against outside genotypes (Darwin, 1890; Johnson and Steiner, 2000; Tremblay et al., 2004). On the other hand, many plant species possess simple radially symmetrical and stereotyped flowers, seemingly lacking the traits that would yield control over visitors and their behaviors (Gong and Huang, 2009; Yoder et al., 2020; Keasar and Wajnberg, 2025). Conserved pollination strategies (a form of biotic niche conservatism), arising through differences in evolutionary rate or developmental constraints such as radial symmetry (Neal et al., 1998; Fenster et al., 2004; Muchhala and Potts, 2007), would lead to a lack of divergence between pollinator assemblages that visit closely related species and would therefore lead to opportunities for gene flow. While some taxa have elaborate pollinator exclusion mechanisms of a mechanical nature (Schiestl and Schlüter, 2009; Van Der Kooi et al., 2021; Zhao et al., 2025), floral displays, as mediated by visual or chemical cues, are the most typical form of plant control over plant-pollinator interactions (Schiestl and Johnson, 2013; Schiestl, 2015) and therefore remain as an important potential driver of hybridization opportunities.

*Heuchera* (Saxifragaceae) is an ideal model to investigate drivers of hybridization in plants because it is one of the most prolific examples of interspecific hybridization, in terms of species number and evolutionary distances (Folk et al., 2016); cytonuclear discord attributable to hybridization involves eight of the fifteen genera in the tribe Heuchereae and a slight majority (59.5%) of species within *Heuchera* (Folk et al., 2017, 2018b, 2023). Conveniently, there are hybridizing and non-hybridizing species in several groups, yielding an evolutionary contrast. The hybridizing complex between *Heuchera americana*, *Heuchera richardsonii*, yielding the hybrid *H. americana* var. *hirsuticaulis*, makes for an excellent study system because: 1) hybridization in this group has seen nearly a century of study and has been confirmed via evidence from both the nucleus and chloroplast genomes as well as morphological and experimental crossing evidence (Rosendahl et al., 1933, 1936; Wells, 1979, 1984; Soltis et al., 1990, 1991; Soltis and Kuzoff, 1995; Folk et al., 2016), 2) members of this group are thought to have intrinsic factors that enable hybridization, e.g. poor pre- and postzygotic isolation mechanisms (Rosendahl et al., 1936; Wells, 1979, 1984), and, 3) *Heuchera* species differ quantitatively in the strength of genetic isolation but all investigated species lack completely efficient prezygotic barriers ((Wells, 1979); see also (Okuyama and Kato, 2009)). Finally, 4) hybridization in *Heuchera* is almost exclusively at the diploid level (Folk and Freudenstein, 2014), freeing its investigation from the equally interesting but confounding effect of (allo-)polyploidy (Folk et al., 2018a).

Although possessing the traits of a compelling hybridization model, investigation of pollinator interactions in *Heuchera* remains relatively poor to date (reviewed in Folk et al. 2021, but see (Segraves and Thompson, 1999; Okuyama et al., 2008; Gaier et al., 2023a; Fogel et al., 2024)). Previous investigators have described *Heuchera* species as generalists with different floral morphologies targeting differing pollinator guilds in some species (Folk et al., 2021). However, *Heuchera*’s close relatives have been well-characterized for plant-pollinator interactions (Robertson, 1925; BROCHMANN and HÅPNES, 2001; Okuyama et al., 2004, 2008; Friberg et al., 2013, 2019; Okamoto et al., 2015; Robson et al., 2019; Gaier et al., 2023b) and found to be specialists; adding additional context from *Heuchera* would assist in better understanding the relationship with gene flow. Of particular interest to this study are the investigations into *Asimitellaria* (Okuyama et al., 2004, 2008, 2018; Okamoto et al., 2015) and *Lithophragma* (Goldblatt et al., 2004; Friberg et al., 2013, 2019), because they demonstrate that these genera, also belonging to taxonomic tribe Heuchereae (Saxifragaceae), exhibit: 1) floral volatiles as the primary mediator of plant-pollinator interactions (similarly *H. americana*, *H. richardsonii*, and their hybrid lack a showy visual display), 2) floral scent profiles that are hypervariable with interspecific and intraspecific variation, and 3) diverse specialized pollinator assemblages that correlate with floral scent profile variation within and among species. Additional study is needed to yield further that 4) low interspecific gene flow is associated with divergent floral scent profiles (limited observed hybrids are reported in *Asimitellaria*, itself an allotetraploid, and *Lithophragma*, but more than half of the genus *Heuchera* is thought to be of hybrid origin with many events (Folk et al., 2016)). Although there have been anecdotal observations of potential pollinators of *Heuchera* and limited studies of isolated species (Segraves and Thompson, 1999; Okuyama et al., 2008), there has never been a systematic effort to record a comprehensive list of effective pollinators for *Heuchera* (reviewed in Folk et al., 2021). Likewise, neither have floral volatile attractants been studied, although adding these data would utilize the strong context of similar data in related genera.

In general, rapid biotic niche divergence would erect barriers to inter-/intraspecific gene flow in closely related species; by extension the aim of the present study is to test what we may term the biotic niche conservatism hypothesis, that is, that the rate of evolution of plant-pollinator interactions and other aspects of mating strategy mediate rates of interspecific gene flow. Although biotic niche conservatism has been evoked as a driver of speciation (Kozak and Wiens, 2006) and the link between hybridization and scent attractants has been explored (Marques et al., 2016), this work lacks a hybridizing-non-hybridizing contrast and to our knowledge ours is the first study to directly test the potential role of attractant evolution as a mechanism for rate variation in hybridization. Testing this hypothesis would require us to show that conserved attractants are associated with observed hybridization; see Figure 1. Thus we built a comprehensive population genetic framework in the *Heuchera americana* var. *americana × H. richardsonii* hybrid zone and paired this with geographically comprehensive floral volatile organic compounds (VOCs) and pollinator visitation data. Our overall aim was to test the predictions that (1) floral VOC overlap predicts pollinator visitation and that (2) conservatism of pollinator assemblages can specifically predict gene flow in a hybridizing-non-hybridizing contrast.

**Fig. 1.**
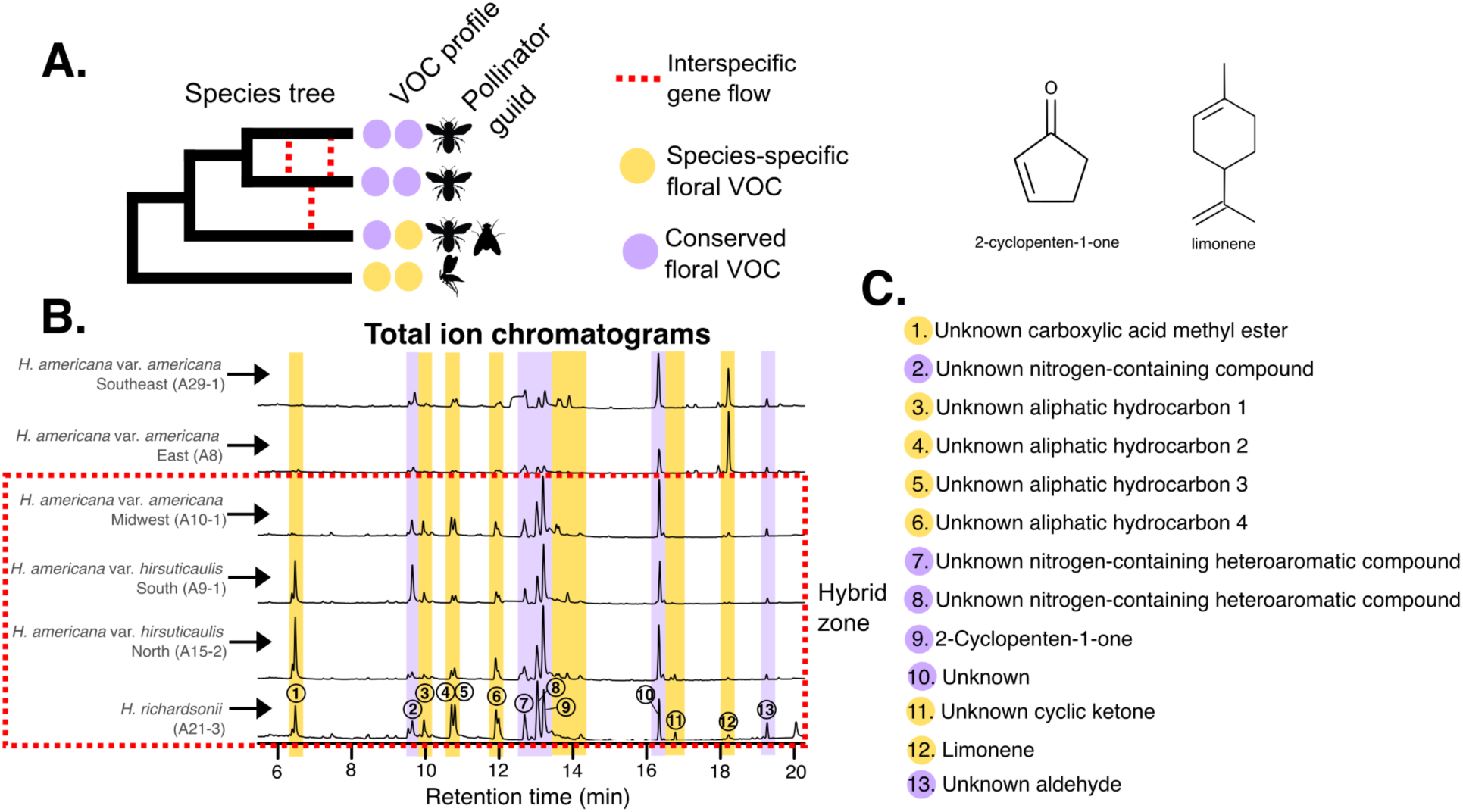
A) A conceptual model showing how conserved elements of a floral scent profile (purple circles) can enable interspecific gene flow (red dotted lies) via shared effective pollinators (shared insect symbols) while differences in elements of floral scent profiles (yellow circles) can attract different pollinators (unique insect symbols) leading to less gene flow. B) Geographically representative examples of total ion chromatograms (TIC) where shared volatiles are highlighted in purple and trace compound differences are highlighted in yellow. (B) legend indicating species for which the TICs (on the left) and pollinator assemblage (on the right) belong. (C) Compound identification (only to compound classes with quality scores >50). Two compounds were confidently identified, 2-cyclopenten-1-one and limonene (structures shown), which were also the most prevalent compounds varying between the parents.

## Materials and methods

### Taxon sampling

We collected living plants, field-collected genetic material, and dried herbarium specimens spanning a majority of the parental taxa and the entirety of the hybrid’s ranges (see Fig. 5 in Rosendahl et al. 1936; Figs. 55, 56, and 64 in Wells 1984), extending from Louisiana into Michigan. We collected one to ten individuals from field sampling localities for the purpose of DNA sequencing, and collected one living individual, when possible, for cultivation at Mississippi State University for greenhouse-controlled floral scent profiling. Living individuals were chosen to maximize geographic and taxon representation, as well as overlap with field studies (below). Herbarium specimens were chosen to cover additional areas of the range not covered by field studies, as well as outgroup comprising *H. alba, H. bracteata, H. caroliniana, H. cylindrica, H. eastwoodiae, H. elegans, H. glabra, H. glomerulata, H. inconstans, H. longiflora, H. longipetala, H. maxima, H. mexicana, H. micrantha, H. missouriensis, H. novomexicana, H. parviflora, H. parvifolia, H. pubescens, H. puberula, H. sanguinea, H. villosa, H. woodsiaphila,* and *H. wootonii*. Sampling for individual aims is reported below.

Field identifications of hybrids and parental species were accomplished using hypothesized ranges and morphologies as outlined in (Rosendahl et al., 1936) and (Wells, 1984). Since there is a gradient in morphology among the taxa, a three-prong approach was necessary for identifying some of the *Heuchera* with intermediate morphology. *Heuchera americana* var. *americana* was the identification of those specimens with smaller radially symmetrical flowers, exserted stamens, and glabrous petioles; *H. richardsonii* circumscribed those specimens with hairy petioles, larger bilaterally symmetric flowers, and a majority of stamens included or level with the sepal margins; the hybrid comprised those specimens with hairy petioles and exserted stamens but intermediate between the two parental taxa in flower size and degree of floral zygomorphy; see Fig. 2A. When taxonomic diagnostics based solely on morphology were questionable, species ranges were considered and coupled with placements recovered in genetic data to determine identities for the purpose of downstream analysis.

**Fig. 2.**
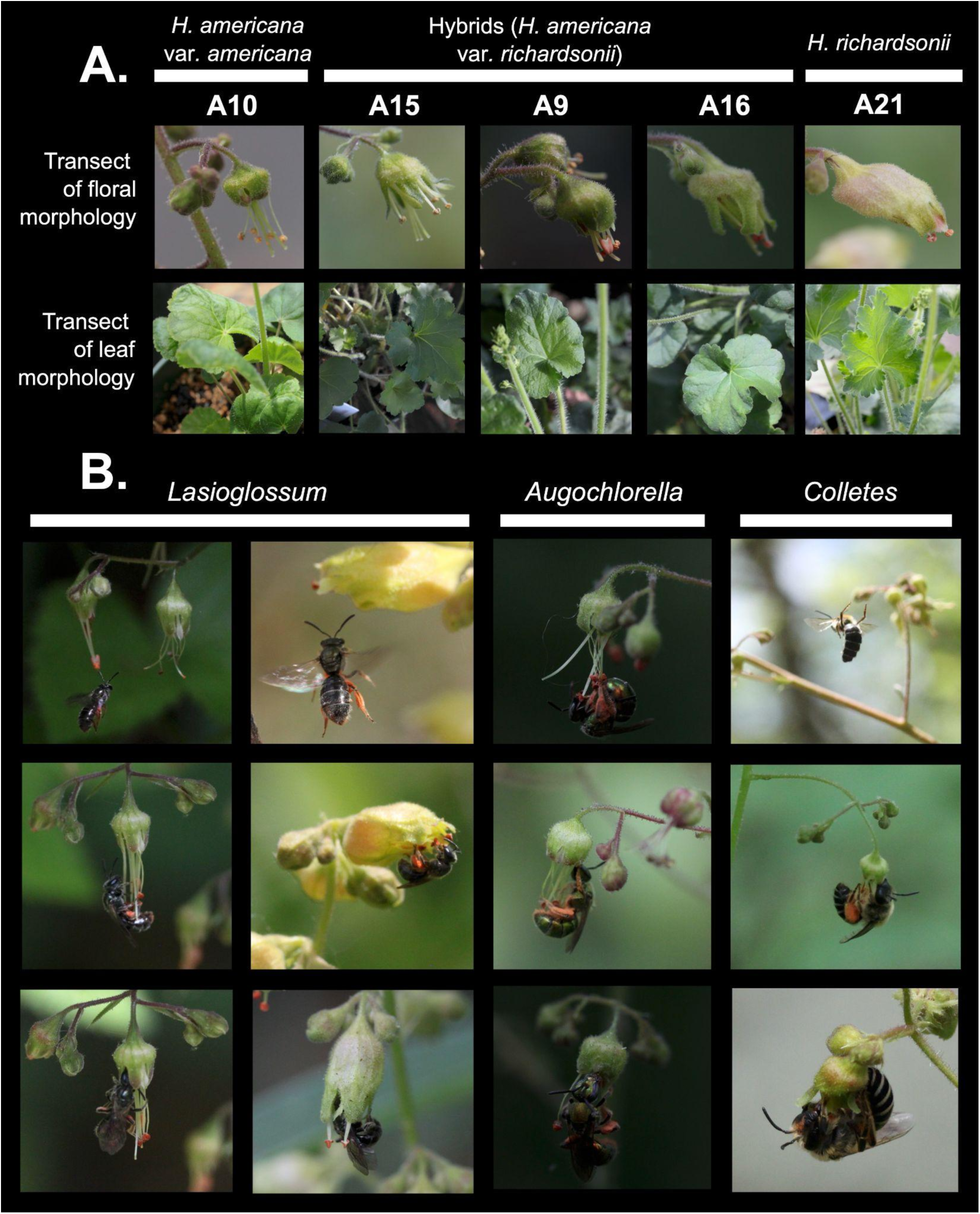
(A) Morphological characters of parents and hybrids, from plants grown in a common garden, sampled from populations in an approximate transect across the *Heuchera americana* × *H. richardsonii* hybrid zone from Indiana (left) northwest to Minnesota (right). First row: flower morphology, with the extreme left representing the *H. americana* var. *americana* parent sampled near the hybrid zone, and the extreme right representing the *H. richardsonii* parent, also sampled near the border of the hybrid zone. The middle three forms show a morphological gradient from more radially symmetrical flowers with strongly exerted stamens and styles characteristic of the *H. americana* parent (A15) to more zygomorphic flowers with included stamens characteristic of the *H. richardsonii* parent (A16). Second row: Leaf morphology, representing the same populations in the same order as flowers. Hybrid forms are characterized by hairy petioles; some also have stronger leaf lobation. Both of these traits are characteristic of the *H. richardsonii* parent. (B) Action shots in the wild of the primary effective visitors, *Lasioglossum*, *Augochlorella*, and *Colletes*. Columns from top to bottom show sequential behavior from scenting to tube entry. In *Lasioglossum* two series are shown for *H. americana* (left) and *H. richardsonii* (right), similar despite very different flowers.

### Phylogenetic construction

We first sought to perform further population-level DNA analysis to confirm hybrid status across the hybrid zone, which previously relied on only limited genetic sampling (Soltis et al., 1991; Folk et al., 2016) and primarily morphological evidence (Rosendahl et al., 1936; Wells, 1984). Subsequently, we aimed to derive phylogenetic distances among plants for comparison to floral volatile and pollinator observation data as outlined below. Thus, a suite of related phylogenetic methods was used to investigate hybrid placement relative to parental taxa. DNA extraction used a modified version of CTAB DNA extraction approach (Doyle and Doyle, 1987) as described in (Folk and Freudenstein, 2014), first building Illumina libraries using NEBNext Ultra II prep kits (New England Biolabs, Ipswich, Massachusetts), and then performing sequence capture with biotinylated RNA baits targeting a customized panel of 277 low-copy nuclear loci specific to *Heuchera* (as reported in (Folk et al., 2015)), followed by sequencing on an Illumina Hiseq 3000 instrument (Novogene, San Diego, California).

We assembled nuclear phylogenomic data using aTRAM (Allen et al., 2018), a reference-based assembler; the assembly module used was SPAdes (Bankevich et al., 2012) with a maximum of 5 iterations. Ortholog choice for tree reconstruction was based on maximum bitscore against the reference. Nuclear assemblies were trimmed for missing data by removing nucleotide sites with greater than 50% missing data and taxa with greater than 20% missing data. Additionally, once the bulk of missing data were removed from the alignments, low-quality or non-homologous sequence portions of individual samples (which could arise from assembly errors, large indels, or other sources) were further detected and removed using spruceup (Borowiec, 2019). All default settings were used, no guide tree was specified, and trimming cutoffs were set to 0.95, 0.97 and 0.99 to assess the impact of different cutoffs, with 0.95 chosen for downstream analysis. Then we inferred phylogenetic trees using a concatenated analysis in RAxML-NG (Kozlov et al., 2019) with a GTR+Γ model with genes treated as distinct partitions. Additionally, we ran a coalescent ASTRAL analysis (Mirarab and Warnow, 2015) with gene trees inferred using RAxML. Each gene alignment was analyzed without partitions within genes and branches in the result with less than 10% bootstrap support were collapsed per developer instructions but otherwise following the concatenated analysis. The phylogeny was rooted with the clade containing *Heuchera maxima* and *Heuchera villosa*. After cleaning the data, we yielded a phylogeny with 532 ingroup and 123 outgroup specimens, totaling 655 specimens. Of the ingroup, we had 297 *H. americana* var. *americana*, 135 *H. americana* var. *hirsuticaulis*, and 100 *H. richardsonii* accessions (Supplementary Table S1).

### Network construction

To corroborate previous work that detected hybridization in this species complex via comparison of nuclear and plastid DNA (Soltis et al., 1991; Folk et al., 2016), we sought to detect hybridization in the nuclear genome, involving the focal members of this study, by inferring networks using PhyloNetworks (Solís-Lemus et al., 2017). PhyloNetworks estimates the best-fit network given gene trees in a model comparison framework, and estimates the relative parental contribution parameter *γ*. These networks were built using the RAxML gene trees inferred above, parameterized with a minimum of zero and a maximum of five hybridizations (i.e., *h_max_* = 0-5) allowed and 50 runs for each *h_ma_*. The output networks files were then examined for the networks with the best pseudolikelihood value, and examined for potential inference errors (i.e., hybridization edges that disallow networks to be rooted), for each *h_max_*=0-5. The overall best pseudolikelihood value, and a plot made to visualize pseudolikelihood for each *h_max_*=0-5, were used to determine the best network.

### Genetic ordination

Species networks do not directly provide population information such as differences in ancestry, so an additional analysis was conducted ordinating the principle axes of genetic variation. The concatenated nuclear genetic matrix was taken as a SNP dataset and a distance matrix was calculated in MegaX (maximum composite likelihood with the following options: uniform rates, transitions and transversions, pairwise missing data deletion). This dataset was then subjected to multidimensional scaling (MDS) in R and colored by taxonomic determination. In this type of analysis, we expect to recover parental taxa as discrete genetic clusters and hybrids as a smear or cluster between them (Wu et al., 2018), the relative position determined by parental contribution and isolation of the hybrid descendant.

### Floral scent profiling

Living specimens comprising 46 outgroup and 43 ingroup accessions (totaling 89 specimens) from 69 populations were collected in the field and brought to Mississippi State University and held for greenhouse-controlled floral scent profiling of as many of the populations visited as possible. Once flowering commenced, we processed flowers following an optimized version of the GC-MS protocol of (Okuyama, 2015), outlined as follows. At the beginning of each data collection day, a SPME fiber (85μM CAR/PDMS StableFlex fiber, Supelco/Sigma-Aldrich, Bellefonte, PA, USA) was put in the GC-MS inlet for a 5 minute “burn out” to desorb any ambient volatiles on the fiber and output data were used to identify background noise. The purged SPME fiber was then used to adsorb floral volatiles in a closed silicone capped glass vial headspace for 30 minutes at 30°C, being as careful as possible to control for ambient or introduced VOC contamination. Two glass vial sizes (2 mL or 4 mL) were used to accommodate the differences in flower sizes between the smaller *H. americana*, intermediate *H. americana* var. *hirsuticaulis*, and the larger sized *H. richardsonii*, but otherwise the headspace was standardized. Each vial was loaded with 5-10 flowers to leave ∼3/4 headspace so a SPME fiber could be inserted without touching flowers for the adsorption of floral volatiles. Each vial was counted as one replicate and, when possible, we sampled three replicates for each individual plant specimen, totaling 232 replicates or an average of 2.6 replicates per specimen. These were then subjected to GC-MS (gas chromatography mass spectrometry) using an Agilent 5975C series machine equipped with a Mass Selective Detector (MSD) and Triple-Axis High Energy Dynode (HED) Electron Multiplier (EM) detector. The column was a J&W DB-1 GC Column part number:123-1063 UNSPSC Code 41115710, with 60 m length, 0.32 mm inner diameter, 1.00 µm film thickness, and 7-inch cage. UHP helium was used as the carrier gas and the inlet settings were 6.3392 psi, a septum purge flow of 3 mL/min, splitless mode, purge flow to split vent 50 mL/min at 0.75 min and gas saver on 20 mL/min after 2 min, and an injector temperature of 250°C and thermal auxiliary #2 heated zone of 280°C. The column settings were a flow rate of 1.2661 ml/min at a pressure of 6.3392 psi, with average velocity of 28.778 cm/sec, holdup time of 3.4749 min, constant pressure setting, and post run set to 4.2522 psi. Temperature programming was as follows: Initial oven temperature was 35°C, hold time ending at runtime 3 minutes, followed by an initial ramp at a rate of 15°C/min until temperature reached 150°C where it was held for 5 minutes ending at runtime 15.667. A second ramp happened at a rate of 50°C/min until temperature reached 250°C where it was held for 5 minutes ending at runtime 22.667. The final ramp was at a rate of 30°C/min until 280°C was reached and this was then held for 5 minutes ending at the final runtime of 28.667; see Supplemental Table S2. In addition to each day’s initial SPME purge run, additional randomized purged runs were performed to control for changes in ambient VOCs throughout any given data collection day. Likewise, dissected floral parts and leaf samples were tested for volatile profiles to control for background floral damage and other plant volatiles not associated with the floral attractants.

Total Ion Chromatograms (TIC) were used to characterize the floral scent profiles of each specimen and individual peak areas (the sum of mass to charge ratios (m/z) at a specific retention time) were used to quantify individual signals. Estimates of peak identification (compound classes) were made using NIST 08 Mass Spectral Search Program (National Institute of Standards and Technology) library. Floral scent profiles were established by first averaging TIC replicates of individual specimens; then the consensus TIC was trimmed of background noise by establishing a baseline peak height cutoff and removing peaks that were present in all samples and negative control runs; and finally, presence or absence of peaks were recorded as well as their relative abundance, calculated as area under the curve using the Agilent 5975C acquisition software. From our initial sampling efforts of 89 specimens, the data cleaning steps left us with 41 ingroup and 43 outgroup (84 total) specimens for floral scent profile characterization.

### Pollination observations

From the populations we visited for the collection of genetic materials, we returned to 40 populations, selected to achieve an even distribution across the range of the hybrid zone and parents, particularly the wide-ranging *H. americana* var. *americana*; we also prioritized high population density because some sites had very low visitation (see Results). While on site, special attention was made to mitigate our contributions to floral attractants, i.e. no floral prints, perfumes, food, etc. Each population was observed at different times throughout the course of the day; further night observations were made in one population to verify the lack of nocturnal visitors. Observations were recorded in fifteen-minute intervals, with observation bouts ranging from fifteen minutes to a few hours and duplicate observations at either a different time frame within a day or on a different day, when possible. Observation data points included GPS coordinates, locality description, weather conditions, inflorescence and flower number counts, estimates of fruit set, nectar reward measurements, pollinator morphotypes, insect visitor behavior, and color of pollen on insect visitors. Weather conditions included temperature, cloud cover, relative wind, and precipitation. All inflorescences were counted in each population and flower estimates were made by counting flowers on three average-sized inflorescences, averaging, and multiplying by inflorescences present. Fruit set (proportion) was measured as the ratio of mature/ing fruits divided by the sum of mature/ing fruits and aborted fruits; this ratio was extrapolated in the same manner as flower count. Nectar measurements were taken under two experimental conditions, plants growing in the wild and plants grown under greenhouse conditions, and were made by dabbing a glass capillary tube (0.4 mm inner diameter, 75 mm length; Drummond Scientific Company, Broomall, Pennsylvania) on the nectar disk at the base of the hypanthium. Measurements were then taken in millimeters using a ruler and converted to microliters.

Floral visitors were defined as any animal that made contact with any of the reproductive organs of a flower. This distinction was made because we often saw predatory or parasitic organisms present on inflorescences or the outside of flowers. Floral visitor behavior recordings included scenting behavior, approach manner, and notes of nectar collection and/or pollen collection. When pollen was observed on potential pollinators, the color was recorded. While *Heuchera* pollen is not microscopically distinctive, the color serves as an indicator; in the study species it presents a bright saffron orange (Wells 1984) and no similar pollen in nearby species was noted. Floral visitors were photographed, collected, and identified by Dr. Olivia Messinger Carril; all floral visitors were insects and predominantly bees. Collection of insects involved capturing them in a one-gallon Ziplock bag and euthanizing them with acetone. Floral visitors were deemed potentially effective pollinators if they were observed with the distinct orange *Heuchera* pollen and this identification was corroborated by pollen metabarcoding (see below), or observed interaction with the floral reproductive organs and likewise corroborated by pollen metabarcoding results. For a host plant species, the effective pollinator assemblage (EPA) was taken as the census of all effective pollinators recorded.

### Metabarcoding

Floral visitors were prepared for molecular analysis in an AirClean 600 PCR Workstation decontaminated on each session with a 50% bleach solution followed by fifteen minutes of UV light. DNA extraction of pollen was performed by removing one, ideally pollen-laden, leg and performing a modified silica column DNA isolation method based on (Williamson et al., 2014) and optimized here for insect materials. Legs were loaded in a round bottom 2 mL Eppendorf tube with six sterilized zinc plated #6 steel beads and 500 *μ*L lysis buffer solution (see Williamson et al. 2014). This sample was then ground in a Fisherbrand bead mill 24 at a speed of 3.10 m/s for durations of thirty seconds, for four cycles with a five second pause delay between each cycle, which was rerun until all samples were homogenized. Ground samples were incubated in a water bath at 65°C for twenty minutes, after which they were rehomogenized in the bead mill and rerun with the above programming and then incubated again for ten minutes at 65°C. Samples were then centrifuged at 10,000 g for two minutes and the supernatant was transferred into another 2 mL Eppendorf tube with 200 *μ*L of potassium acetate solution (see Williamson et al. 2014) and chilled at -20°C between one and twenty-four hours. Following the -20°C incubation period, centrifugation was performed for thirty minutes at 10,000 g and the supernatant was transferred into another tube containing 1.2 mL of guanidine solution. The guanidine-supernatant mixture was then added to a spin filter column in 700 *μ*L batches and centrifuged for two minutes at 10,000 g, until the entire volume was transferred through the filter. The flow-through was discarded, then the filter was washed with 500 *μ*L of wash solution (see Williamson et al. 2014) followed by 80% ethanol, both times being centrifuged at 10,000 g for two minutes and again discarding the flow-through. The spin column was then dried by spinning for five minutes at 10,000 g, and then transferred to a new two mL Eppendorf tube. Finally, 200 *μ*L of elution solution (see Williamson et al. 2014) was added to the spin filter column where it was incubated at room temperature for ten minutes and then spun for ten minutes at 10,000 g.

For optimization of metabarcoding we aimed to characterize pollen taxa using both the nuclear and chloroplast genome. For the nucleus, we used the internal transcribed spacer region (primer pair ITS1 and ITS2 (White et al., 1990; Ankenbrand et al., 2015; Coghlan et al., 2021)). For the chloroplast, there was no clear choice in the literature so we tested primers for *trn*L (UAA) intron primer pairs tabg and tabh (Taberlet et al., 2007), chloroplast *mat*K primers matK2-F and matK2-R (Coghlan et al., 2021), and chloroplast *rbc*L primers orbcL2-F and orbcL2-R (Coghlan et al., 2021). Of the potential metabarcodes tested, only ITS and rbcL were able to distinguish *Heuchera* from other Viridiplantae.

PCR validation reactions for both ITS and *rbc*L followed the same ratios for one reaction with a total volume of 20 *μ*L. Each reaction contained 6.6 *μ*L H_2_O, 10 *μ*L DreamTaq master mix, 0.2 *μ*L F primer (10 *μ*M), 0.2 *μ*L R primer (10 *μ*M), and 3 *μ*L DNA template. The F and R primers for ITS and rbcL, respectively, were ITS1/ITS2 and orbcL2-F/orbcL2-R; each primer had a 5’ sequence tag added for compatibility with Illumina library chemistry following the sequencing core guidelines. Thermocycler programs for both ITS and *rbc*L began with an initial hold at 95°C for three minutes, followed by 35 cycles beginning with a hold of thirty seconds at 95°C then one minute at 50°C and then one minute at 72°C, next there was a hold at 72°C for ten minutes, and finally a thermocycler hold at 4°C. Once PCR was complete, DNAs were quantified using an Invitrogen Qubit HS dsDNA (Invitrogen, Waltham, Massachusetts, U.S.A.) assay kit in a Qubit 4 fluorometer. Samples were also analyzed via electrophoresis using 2% agarose gels and SYBR Safe (Invitrogen) stain to verify presence of a product. Samples were submitted for sequencing on an Illumina MiSeq instrument (300 bp paired end reads) by the Michigan State University RTSF facility.

QIIME 2 (Bolyen et al., 2018) was used for both *rbc*L and ITS metabarcoding reads to assess the taxa present in pollen samples. The taxonomic database used for ITS was UNITE (Nilsson et al., 2019) and the *rbc*L database was taken from (Bell et al., 2017b; Bell, 2021). ITS reads were used primarily for taxon identification; *rbc*L metabarcoding reads were additionally used to indicate which chloroplast clade (sensu Folk et al. 2017) the *Heuchera* pollen could be assigned as the parents differ in this respect. To perform chloroplast clade assignments, the original *Heuchera* records from Bell et al. (2017; 2021) were removed and replaced with vetted *rbc*L records extracted from recent chloroplast genome reports (clade A: *Heuchera alba* MN496063; clade B: *Heuchera longipetala* var. *longipetala* MN496067; clade C: *Heuchera abramsii* MN496062; all from (Folk et al., 2020)). Fungal and metazoan reads were excluded in both cases after classification for the verification of pollen identities. Following this, samples with zero reads were removed, including negative controls and samples with no detectable pollen. Percent pollen by pollinator was used to create stacked taxon bar plots and evaluate (1) whether *Heuchera* pollen was moved in the legs, which in combination with behavior data would suggest an effective pollinator; and (2) whether the pollen load was primarily *Heuchera*, which may suggest specialist behavior when coupled with pollinator observation that indicated no visitation to surrounding plants. By contrast, percent pollen by host plant species was used to see if the candidate effective pollinators associated with each species differentially moved *Heuchera* pollen and what other non-*Heuchera* hosts were detected.

### Statistical methods

*GC-MS—*Statistical analyses of floral volatile profiles were run in R. A Mantel test using Bray–Curtis dissimilarities of the cleaned GC-MS output and patristic distances was run to determine if phylogenetic structure is observed in the floral scent profiles. This was conducted both with outgroup taxa (i.e., assessing phylogenetic conservatism of floral volatiles at the level of genus) and without outgroup taxa (i.e., assessing phylogenetic conservatism of floral volatiles within the hybrid zone). We then performed parallel Mantel tests using the Jaccard distance, thus considering only the presence or absence of compounds in the floral scent profiles for the conservatism test. Next, a linear discriminant analysis (LDA) was run to evaluate if the sum of floral VOCs (i.e., the floral scent profiles) are able to distinguish taxa, followed by a jackknife to assess whether the discriminant model can predict unknown samples.

#### Pollinator assemblages

The same statistical tests performed on the floral scent profiles were repeated for the effective pollinator assemblages (EPAs) so pairwise comparisons could be made. First, a Mantel test was run in order to investigate whether phylogenetic structure of EPAs could be observed across *Heuchera*. Similarly to floral volatile data, phylogeny was represented with patristic distances and Bray–Curtis dissimilarities were used to compare EPAs. The test was conducted both with outgroups (assessing phylogenetic conservatism of EPAs at the genus level) and with only ingroup *Heuchera* species (assessing phylogenetic conservatism of EPAs within the hybrid zone). Finally, an LDA and jackknife were performed to assess whether EPA composition is predictive of host taxa.

#### Nectar

Because the effect of two independent categorical variables (species and greenhouse vs. wild) were being evaluated for their effect on nectar volumes we considered a two-way ANOVA. However, a Levene’s test for homogeneity of variance indicated that our data did not meet the required assumption of equal variance across groups (*p* = 0.03135). For this reason, we chose to use a Generalized Linear Model (GLM) capable of accommodating unequal variance to determine differences among taxa in nectar reward; see Supplementary Figure 1 for diagnostic plot.

#### Fruit set

A Levene’s test indicated the fruit set data met the assumption of homogeneity of variance (*p* = 0.3313, also see Supplementary Figures 2 and 3 for diagnostic plots) so we performed an ANOVA to determine if there were significant differences in the fruit set ratios between taxa.

**Fig. 3.**
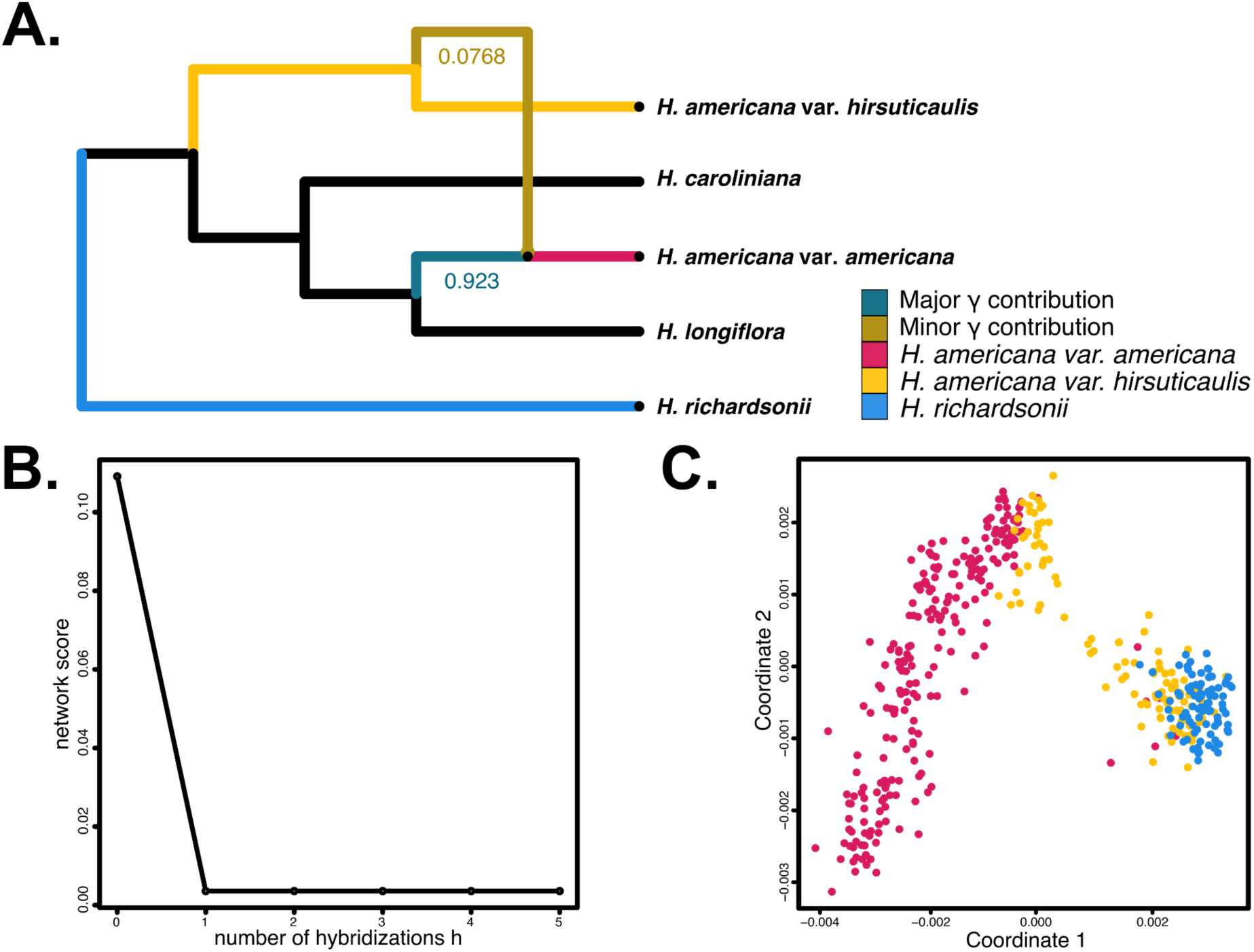
(A) PhyloNetworks best result indicating one hybridization with major genetic contributions to *H. americana* var. *americana* (γ = 0.923) from the lineage giving rise to *H. americana* var. *americana* and minor genetic contributions from an unsampled lineage sister to *H. americana* var. *hirsuticaulis* (γ = 0.0768). (B) Line graph of network pseudo-deviances. On the x-axis are *h_max_* intervals, maximum number of hybridizations (h) allowed for each tested network, and on the y-axis are the -log-likelihood with lower results being a better fit. The lowest -log-likelihood (0.0035959446656174806 for *h_max_* = 1) and broader slope heuristic imply a network with one hybridization node is the best fit. (C) Molecular MDS of nuclear DNA for *H. americana* in red, *H. richardsonii* in blue, and the hybrid in yellow. The MDS shows typical intermediate and overlapping placement of the hybrid with the parental taxa, and clear separation of *H. richardsonii* and *H. americana*, with a couple of notable exceptions of *H. americana* populations overlapping with *H. richardsonii*. The rare cases of *H. americana* that overlapped with *H. richardsonii* are from populations at the extreme western limits of *H. americana*’s range in Arkansas and Oklahoma where you would expect to find either the hybrid or *H. richardsonii*, respectively.

## Results

### Phylogenetics

ASTRAL (Fig. 4), treating samples as independent taxa, rendered a monophyletic *H. richardsonii*, if assigned hybrids are excluded, and a paraphyletic *H. americana* var. *americana*; this is a long-noted topology we are addressing in a future work. Field-identified *H. americana* var. *hirsuticaulis* hybrids exhibited two distinct topological placements, with different populations placing nearest whichever parental taxa to which they are geographically proximal (see similar results in MDS, below). Thus, more northerly populations placed within *H. richardsonii*, and more southerly populations placed within *H. americana* var. *americana*. The backbone had primarily low support values (LPP < 0.5) reflecting high conflict in the data, while numerous subclades were recovered with low to high support (LPP > 0.5; these are noted in Fig. 4). In particular, a strongly supported clade containing primarily *H. richardsonii* and hybrid accessions was recovered with strong support (LPP = 0.95). A corresponding clade comprising most *H. americana* accessions received weak support (LPP = 0.53) but contained numerous subclades receiving support (Fig. 4).

**Fig. 4.**
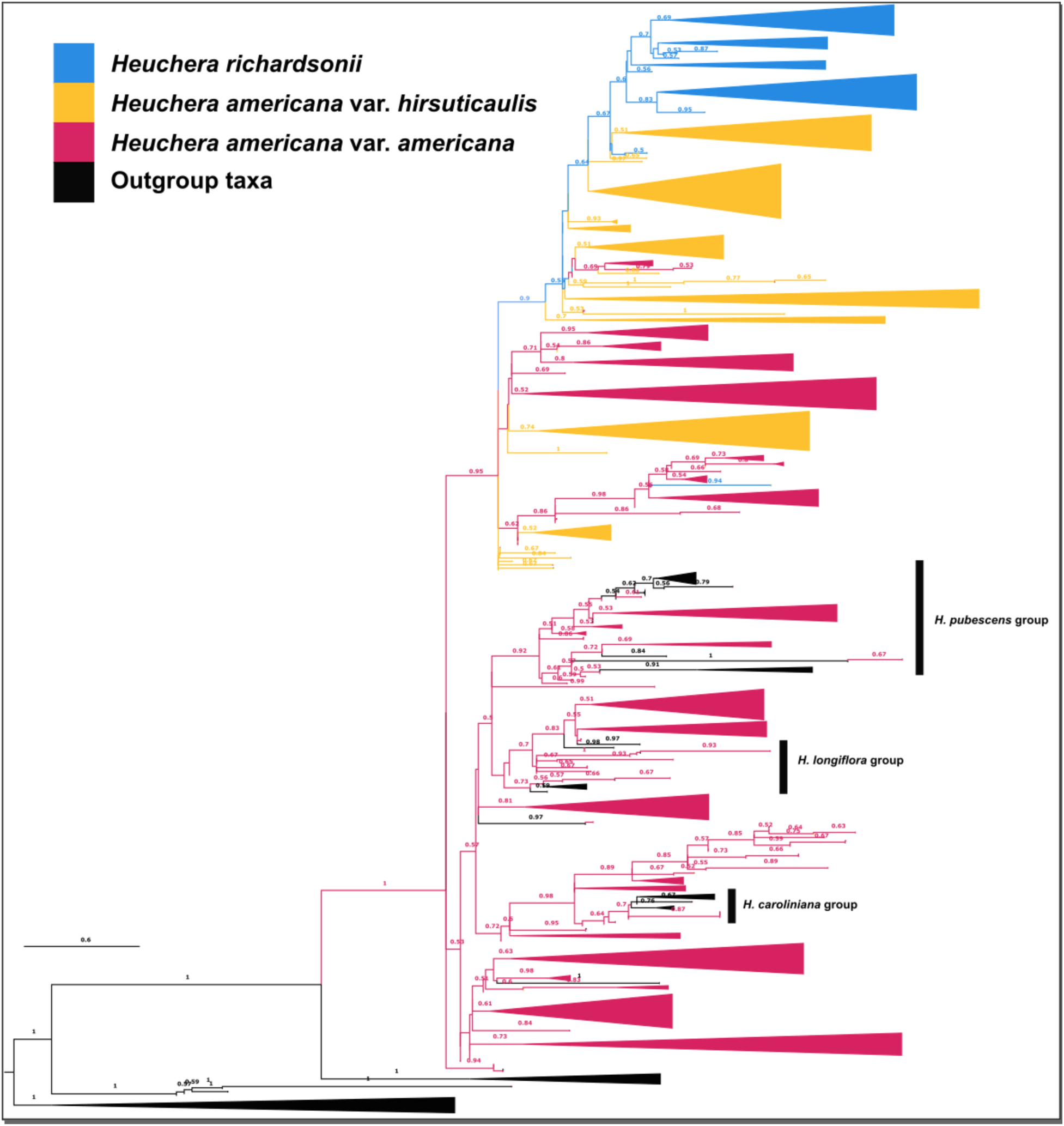
ASTRAL phylogeny of nuclear DNA, with parental taxa marked following the legend; branch labels represent LPP (local posterior probability). Some outgroup taxa marked in black are nested in *Heuchera americana*; these are members of *Heuchera* subsect. *Heuchera* that are outside the hybridizing complex; we found all of them embedded in *H. americana* as currently recognized, indicating *H. americana* is paraphyletic.

RAxML (Fig. 5) results largely mirrored ASTRAL results in topology and primarily differed in support. RAxML, likewise, rendered a monophyletic *H. richardsonii* and paraphyletic *H. americana* when treating samples as independent taxa and excluding hybrids in consideration of monophyly. Again, the field identified *H. americana* var. *hirsuticaulis* hybrids followed the same geographic proximity-based placement pattern in topology. The backbone mostly had low support values (BS < 50) and numerous subclades were recovered with low support, with a few having high support (BS > 50; these are noted in Fig. 5). The clade containing primarily *H. richardsonii* was recovered with moderate support (BS = 77) and only had three well supported subclades (BS > 50) while the corresponding clade comprising most *H. americana* var. *americana* accessions received better support (BS = 86) and had many subclades with low to high support (BS > 50). Overall, RAxML recovered fewer subclades with support (BS > 50) than ASTRAL, but yielded clearer *H. richardsonii* and *H. americana* clades with better support (i.e. the *H. richardsonii* clade contained fewer identified *H. americana* var. *americana* accessions and the *H. americana* var. *americana* clade had higher support).

**Fig. 5.**
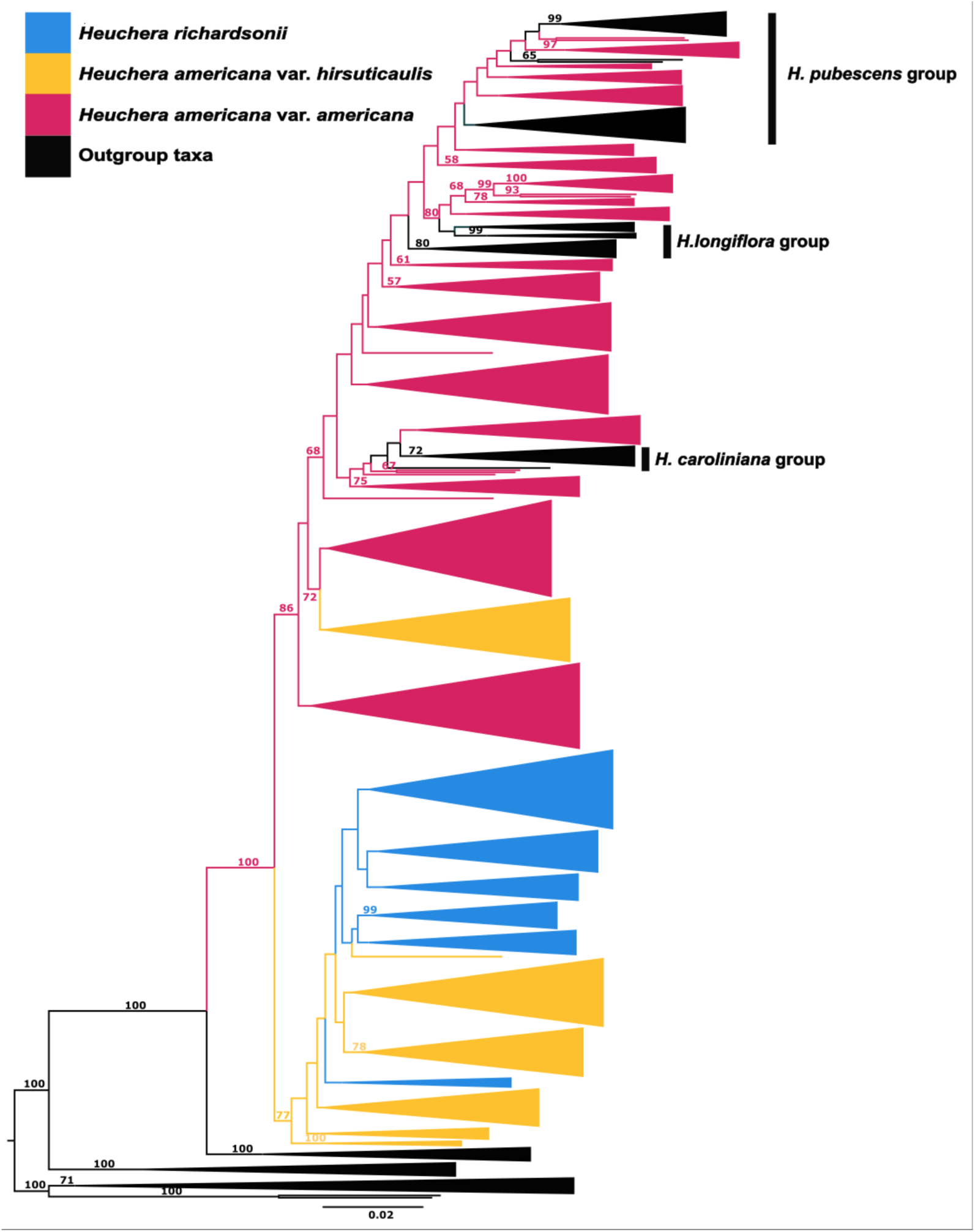
RAxML phylogeny of nuclear DNA, with parental taxa marked following the legend; branch labels represent LPP (local posterior probability). Some outgroup taxa marked in black are nested in *Heuchera americana*; these are members of *Heuchera* subsect. *Heuchera* that are outside the hybridizing complex; we found all of them embedded in *H. americana* as currently recognized, indicating *H. americana* is paraphyletic.

### PhyloNetworks

Our results indicated that the best network had one hybridization event in the lineage leading to contemporary *H. americana*. The genetic contributions to *H. americana var. americana* (*γ* = 0.9232) came from an unsampled lineage sister to *H. americana var. hirsuticaulis* (*γ* = 0.0768); see Figure 3. Polarity of networks can be difficult to interpret (compare Figs. 4, 5); depending on the rooting this analysis can also be interpreted to mean that the ancestor of *H. americana* var. *hirsuticaulis* contains contributions from *H. americana* var. *americana* and an extinct relative of *H. richardsonii*.

### Floral scent profiles

Total Ion Chromatograms (TICs) demonstrated shared peaks among all of the ingroup taxa, showing complex patterns of especially minor volatile compounds and their abundance. Notably, TICs were very similar within the hybrid zone with an apparent geographic pattern of accumulating differences in trace compounds as geographic distance from the hybrid zone increased (geographically representative examples in Fig. 1B; see also Supplemental Table S1). *Heuchera americana* var. *americana* samples outside the hybrid zone had notably different and less uniform TICs compared to those within the hybrid zone. Many compound identifications were low in confidence; therefore, the data were analyzed downstream in the form of TICs grouped by retention time and area under the curve, making for the purpose of comparison the assumption of identity among identical TIC peaks. That said, based on examination of TICs and m/z plots of individual peaks indicated an overall profile comprising mainly aliphatic hydrocarbons, terpenes, and ketone-related compounds. Most prevalent were various unknown nitrogenous compounds, and two confidently identified molecules: limonene (a monoterpene; up to 21%), and 2-cyclopenten-1-one (a cyclic ketone; up to 20% of fragrance). Limonene is a typical floral fragrance compound; 2-cyclopenten-1-one is rarer but resembles more complex substituted molecules previously found in bee-pollinated flowers (Dekebo et al., 2022). Notably, 2-cyclopenten-1-one and limonene were not only prevalent compounds but differentiate the parental taxa better than other compounds (Fig. 1) with the former being in low abundance in *H. americana* outside the hybrid zone and the latter being absent or low-abundance in the hybrid zone and in *H. richardsonii*. Thus, the floral scent in the *H. americana* group varies from terpene-dominated (*H. americana* background) to ketone-dominated (*H. richardsonii* background) fragrances. Dissection and separate analysis of an *H. richardsonii* individual showed that 2-cyclopenten-1-one is not emitted from the floral disc or gynoecium, which emit few volatiles, but is emitted from the androecium in a small amount and mainly from the hypanthium and free sepals. Thus floral fragrance appears to mainly emit from the hypanthial area in this genus.

### Pollinator assemblages

Pollination observations were conducted at 40 unique *Heuchera* populations (47 observation bouts), comprising 32 ingroup and eight outgroup populations (four of the latter were naturally occurring and the remaining four in an urban setting in the native range), and totaled 6,780 minutes or 113 person-hours (see Supplemental Table S4). Insect visitation rates were found to peak during the height of the day between 1100 and 1400. Observations were done during temperatures ranging from 15.6°C to 28.3°C and the rates of pollinator visitations did not change noticeably at any point in the range of observation temperatures. Likewise, pollinator visitation rates did not noticeably change from sunny to cloudy skies, but did drop off during rain and returned to normal frequencies within 40 minutes of rain cessation. Visitation rates tended to be low where the plants were not prevalent and particularly at the edge of parental ranges. Five pollinator guilds (bee, wasp, syrphid, moth, and beetle) were observed visiting *H. americana* var. *americana*, *H. americana* var. *hirsuticaulis*, and *H. richardsonii* populations (Table 1). The average number of pollinators observed visiting each population across our observations was 15.82, 15.2, and 31.33 per host plant taxon respectively. They visited these host taxa at a rate of 119.28, 87.2, and 169.44 visitations per population per day respectively (a day extrapolated as 8 hours of observation, approximating the length of day with active visitors); with pollination rates per hour at each population being 14.91, 10.9, and 21.18 respectively; see Supplemental Table S3. Among all of the *Heuchera* included, the observed pollinator assemblage was 59.17% *Lasioglossum* (Hymenoptera: Halictidae), 21.81% *Colletes* (Hymenoptera: Andrenideae), 12.78% Augochlorini (Hymenoptera: Halictidae; only *Augochlorella* in ingroup taxa), 3.19% *Bombus* (Hymenoptera: Apidae; observed only in outgroup taxa), 1.53% Syrphidae (Diptera), and 1.53% other pollinators; see Supplemental Table S3 for a complete host and pollinator list. Among ingroup populations, pollinator assemblages were predominantly (96.51%) made up of the three bee genera *Lasioglossum* 63.12%, *Colletes* 25.42%, and *Augochlorella* 7.97%; at least two of these are observed at most sites. Among these three key visitors, relative percentages, at individual host populations, of pollinator prevalences appear to display a geographic gradient with *Augochlorella* most important in the southern range and *Lasioglossum* in the north; see Fig. 6. The remaining 3.49% of the ingroup pollinator assemblages consisted of Syrphidae 1.66%, Ichneumonidae 1.33%, Lepidoptera 0.33%, and Coleoptera 0.17%.

**Fig. 6.**
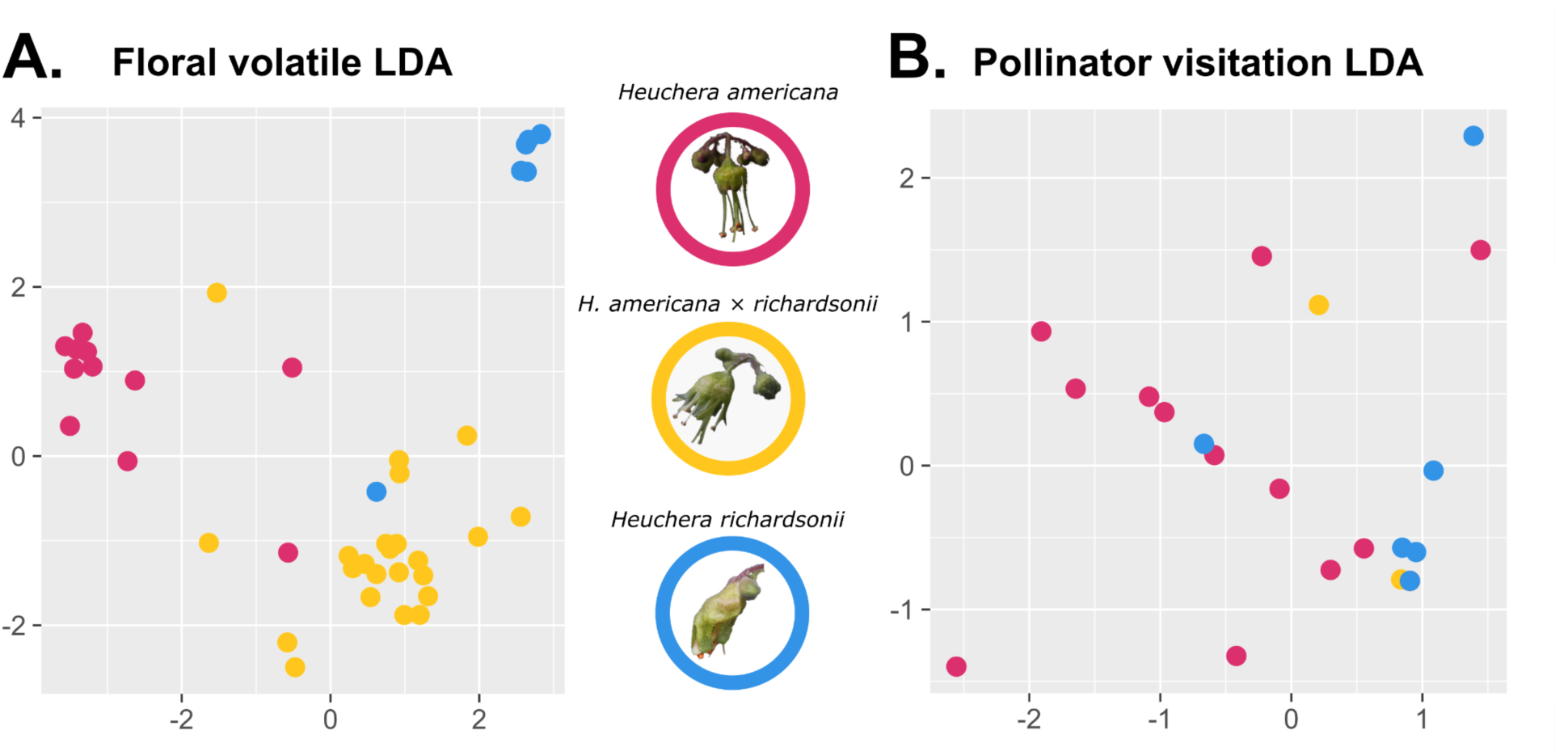
**(A)** LDA of VOCs for ingroup *Heuchera*. (B) LDA of pollinator assemblages for ingroup *Heuchera*.

**Table 1.** Averaged pollinator visitation data by ingroup species. Pollinator averages and observed pollinator visitations are the averages of actual pollinator counts during observation periods, while the rates were extrapolated from the observed pollinator visitations per unit time. Original data from all populations included in this study can be found in Supplemental Table S3.

| Host species | Observed<br>pollinator<br>visitations | Pollinator<br>rate/<br>hour | Pollinator<br>rate/ day<br>(8 hours) | Pollinator/<br>flower/<br>day (8<br>hours) | Pollination/<br>inflorescence/<br>day<br>(8 hours) | *Lasioglossum* | *Augochlorella* | *Colletes* | Syrphid | Wasp | Moth | Coleoptera |
| --- | --- | --- | --- | --- | --- | --- | --- | --- | --- | --- | --- | --- |
| *Heuchera americana* var.<br>*americana* | 15.82 | 14.91 | 119.28 | 0.24 | 11.8 | 8.33 | 1.17 | 4.33 | 0.33 | 0.21 | 0.08 | 0.04 |
| *Heuchera americana* var.<br>*hirsuticaulis* | 15.2 | 10.9 | 87.2 | 0.24 | 6.6 | 7.75 | 0.63 | 0.25 | 0.25 | 0.13 | 0 | 0 |
| *Heuchera richardsonii* | 31.33 | 21.18 | 169.44 | 0.34 | 16.94 | 19.67 | 2.5 | 7.83 | 0 | 0.33 | 0 | 0 |

### Metabarcoding

ITS results (Fig. 8) indicate that the floral visitors found on all of the host taxa were predominantly carrying pollen assignable to *Heuchera* or Saxifragaceae (Fig. 8A). All visitors, except for Syrphidae and Ichneumonidae, were primarily and often overwhelmingly carrying *Heuchera* and Saxifragaceae pollen. This result was consistent with field observations, where the bee floral visitors carrying an obvious pollen load had the saffron-orange pollen of *Heuchera* and were not obviously carrying other colors. Both Ichneumonidae and Syrphidae had negligible visible pollen loads and correspondingly few reads recovered. Ichneumonidae was found to be carrying *Heuchera* and Saxifragaceae pollen in small amounts, but mostly had legume pollen present. Syrphidae reads were solely identified as *Chlamydomonas* (Fig. 8B), likely environmental contamination or a misidentification of the very few reads recovered from syrphid legs.

**Fig. 7.**
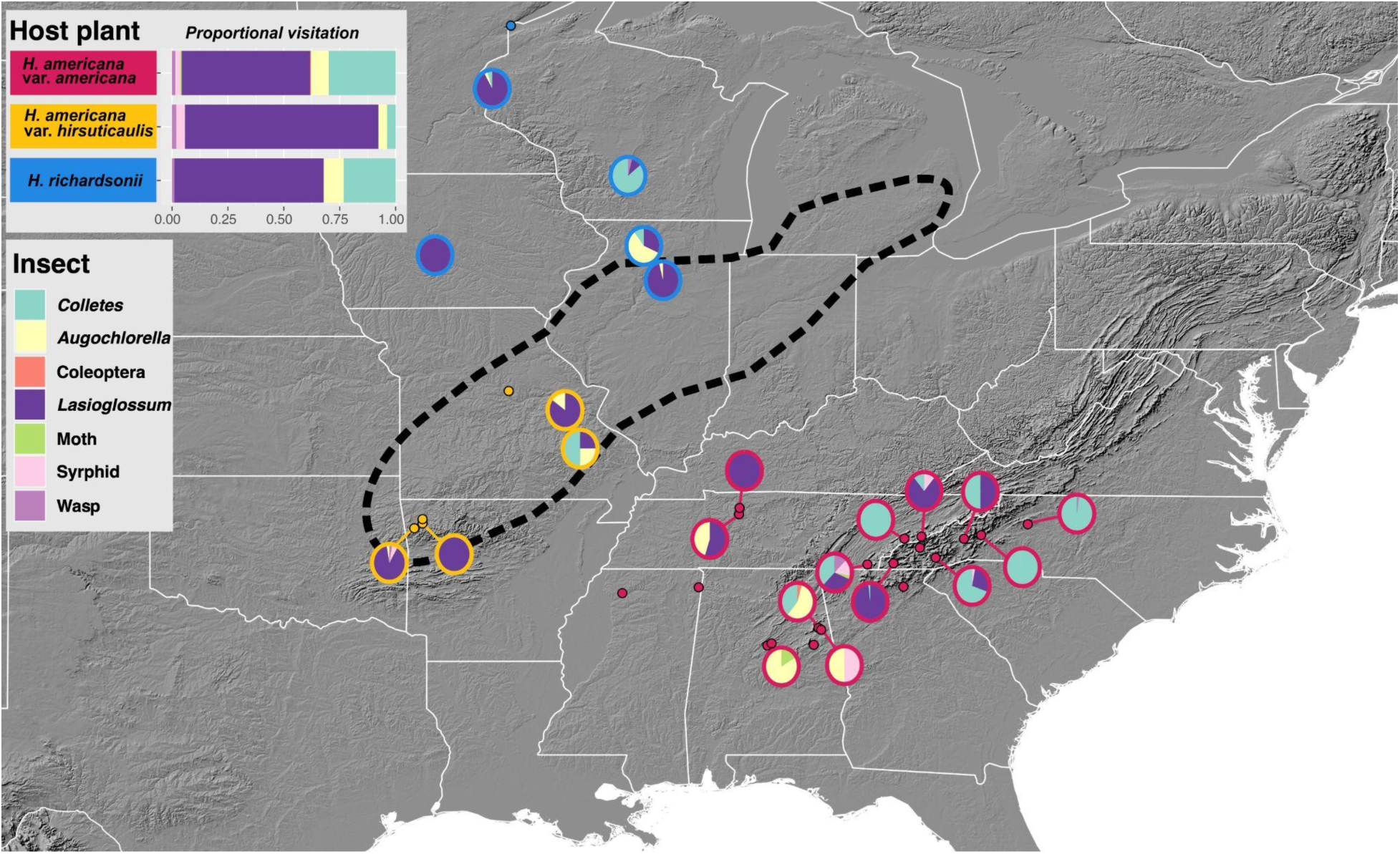
Map showing the geographic distribution of the 32 unique populations of *Heuchera* where pollinator observations were done; the hybrid zone as hypothesized by Wells et al. (1984), is outlined in a black dotted line. Stacked bar plots in the left-hand inset indicate the overall pollinator assemblage of host plant taxa and match the insect legend below. Pie charts in the map are marked over, or connected to, population locality points where pollinator observations were done and indicate the overall observed pollinator assemblage at each population. Outline colors of pie charts and fill colors of locality points match the color of the host plant in the top legend and follow Figs. 3-6. Insects are indicated according to the legend in the left-hand inset. Locality points without a connecting pie chart indicate a population where observations were conducted, but no pollinators were present.

**Fig. 8.**
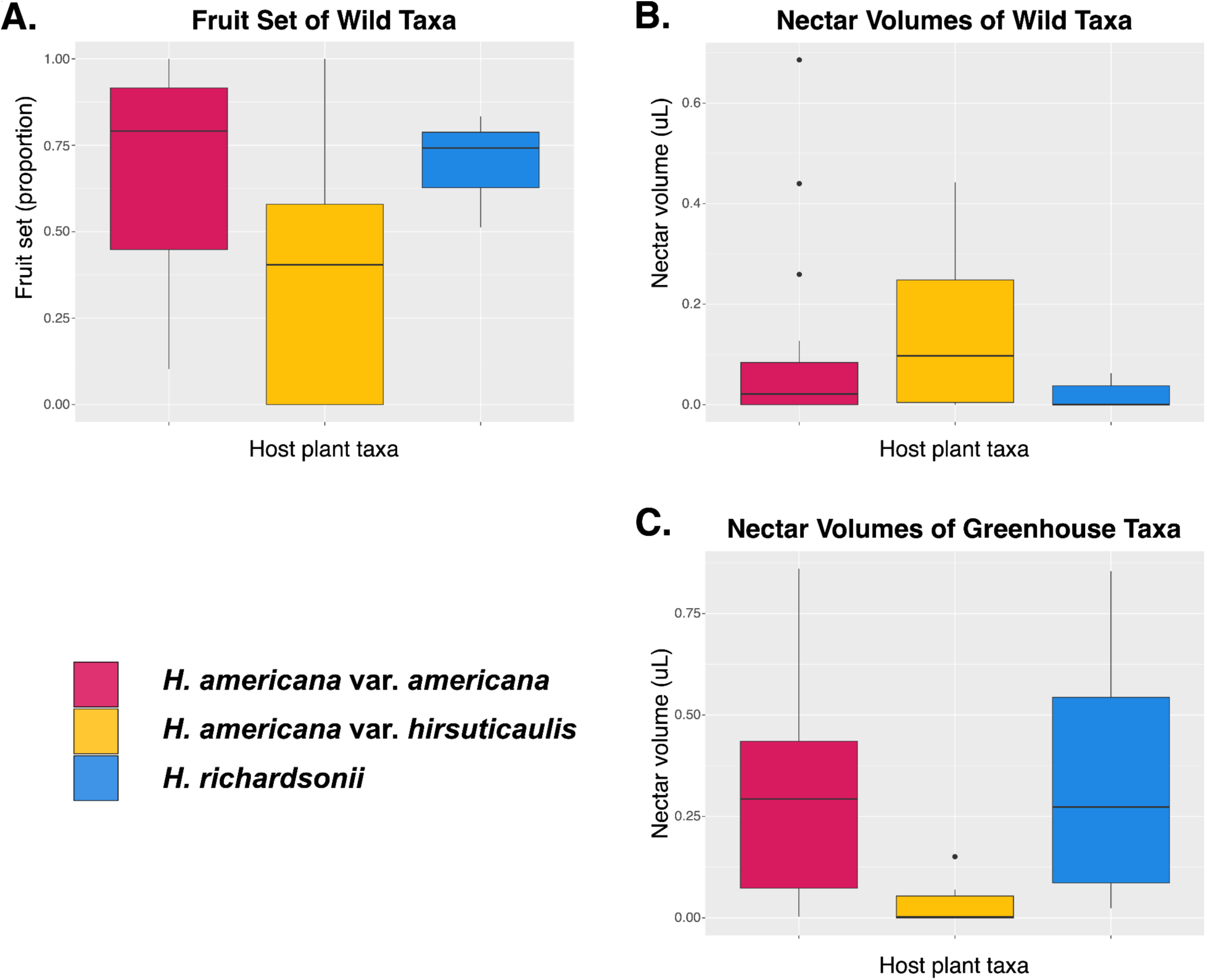
The legend in the bottom left identifies the host plant taxa (following Figs 3-7), or x-axes, for the (A) fruit set box plot representing proportion of fruit set for wild taxa, (B) box plot representing nectar volumes (μL) for the wild host plant taxa, and (C) box plot representing the nectar volumes (μL) of host plant taxa grown under greenhouse conditions.

Chloroplast *rbc*L results (Fig. 9) were approximately similar to ITS, although they estimated a lower percent of pollen from *Heuchera* or Saxifragaceae. As well as identifying pollen taxa, we attempted to assign these to recognized chloroplast clades, since more than one distinct clade occurs in the host plant species (Folk et al., 2017). Our results indicate that the pollinators visiting *H*. *americana*, *H. longiflora*, & *H*. *macrorhiza* were predominantly carrying pollen from Saxifragaceae plastid clade A (Folk et al., 2016). Surprisingly, pollinators visiting *H*. *richardsonii* only had small amounts of pollen on them assignable to plastid clade A, and predominantly non-*Heuchera* pollen (misidentified as Zingiberales), while pollinators found on *H*. *americana* var. *hirsuticalulis* had no detectable pollen present from plastid clade A and were mostly carrying pollen from Fagaceae; see Fig. 9A. This result was inconsistent with ITS and may indicate less taxonomic resolution and more uncertainty in the relatively conserved *rbc*L locus. *rbc*L is much more conserved than ITS and has a less representative reference database, leading to less robust pollen identifications (see also (Bell et al., 2017b, a; Martin et al., 2024)). Among the pollinators, *Colletes* and *Bombus* were almost exclusively carrying pollen of plastid clade A while *Lasioglossum*, *Augochlorella*, and *Nasonovia* were the only other floral visitors carrying plastid clade A, in decreasing quantities. Ichneumonidae and Syrphidae had no detectable pollen from *Heuchera* CP clade A, consistent with the ITS results; see Fig. 9B. Some insect samples were carrying out of season pollen, which may be due to incorrect identifications, ambient pollen presence, or to the nature of the insects’ living conditions, e.g., den hibernation.

**Fig. 9.**
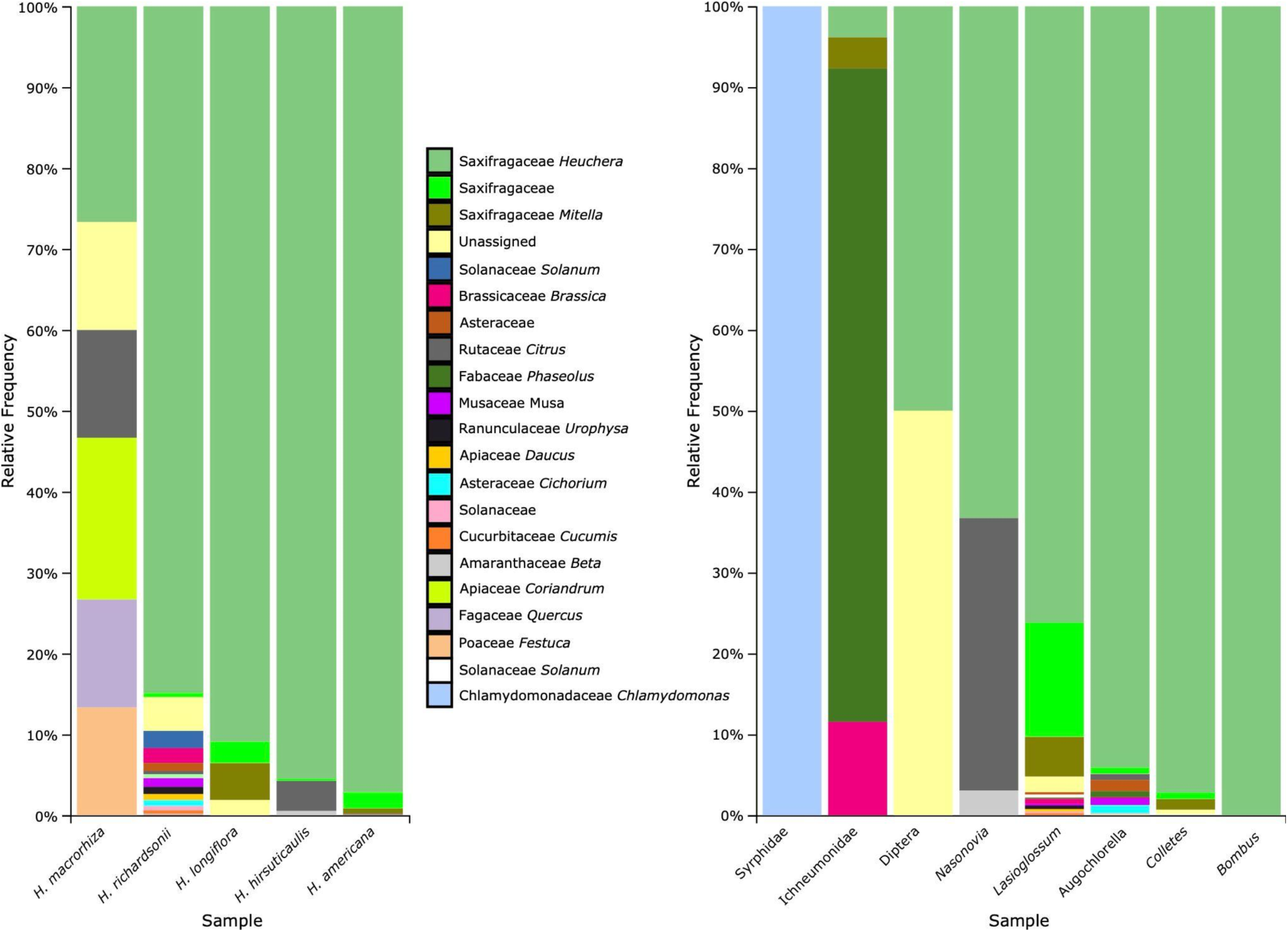
ITS metabarcoding results. X-axes reflect (A) host taxa and (B) pollinators, while Y-axes reflect (A) percent pollen type found on pollinators visiting each host taxa or (B) percentage of pollen type on each pollinator.

**Fig. 10.**
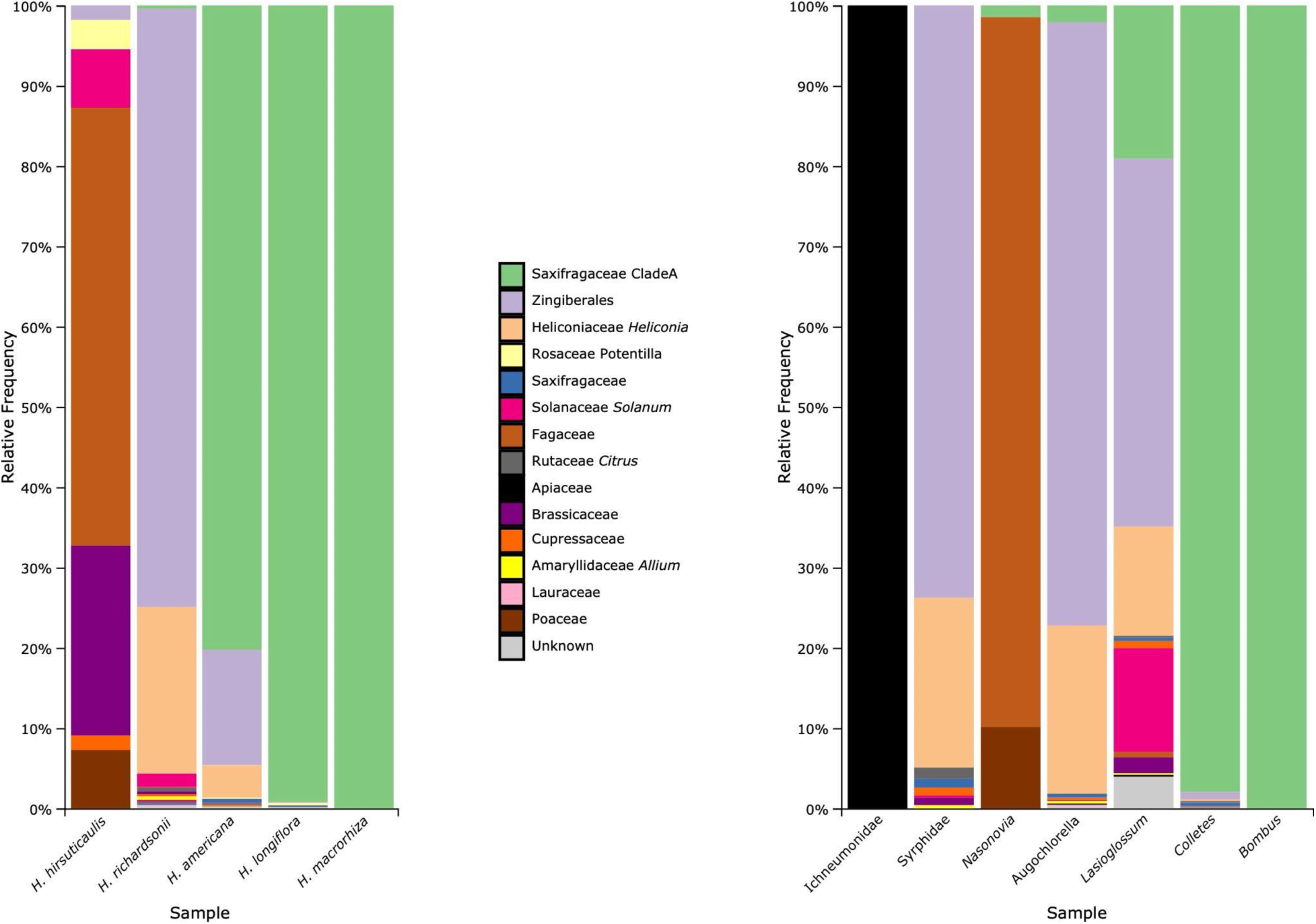
Chloroplast rbcL Metabarcoding results. X-axes reflect (A) host taxa and (B) pollinators, while Y-axes reflect (A) percent pollen found on pollinators visiting each host taxa or (B) pollen on each pollinator.

### Tests for phylogenetic conservatism

#### Floral scent profiles

A Mantel test including both the 41 ingroup and 43 outgroup taxa we characterized floral volatiles for indicated that floral scent profiles were conserved across *Heuchera* (*p* = 0.024). By contrast, Mantel tests of only the ingroup *Heuchera* samples displayed no phylogenetic structure of floral attractants (*p* = 0.14). We then used an LDA to investigate the related question of whether taxa were significantly different (Fig. 5A); this indicated broad overlap of VOCs across ingroup taxa. Although LDA recovers some taxonomic grouping of floral scent profiles (Fig. 5A), a jackknife analysis indicated that there was no better than a random chance of predicting the three taxa using floral volatiles (success rate 37%).

A Mantel test between the Bray-Curtis dissimilarity of ingroup floral scent profiles and geographic distances of ingroup populations indicated that there was no significant geographic structure to ingroup floral scent profiles (*p* = 0.97). Similarly, when considering only the presence and absence of compounds comprising the floral scent profiles of ingroup taxa using Jaccard distances, there was no significant geographic structure to the floral attractants of the ingroup taxa (*p* = 0.90).

### Pollinator assemblages

A Mantel test between the patristic distance of both in- and outgroup *Heuchera* and their observed pollinator assemblages indicated that pollinator assemblages were conserved across *Heuchera* species included in the analysis (*p* = 0.04). By contrast, Mantel tests of the ingroup populations show a lack of phylogenetic structure of pollinator assemblages within the hybridizing complex (*p* = 0.52). Similar to VOC results, an LDA indicated broad overlap in pollinator assemblages of ingroup taxa (Fig. 5B). Again, despite visitation trends differing among the taxa, a jackknife indicated that there was a no better than random chance of predicting the three taxa using pollinator assemblages, with a success rate of 41%.

### Floral, nectar, and fruit set data

Floral and fruit data summaries can be found in Table 2. *Heuchera americana* var. *americana,* the hybrid *H. americana* var. *hirsuticaulis,* and *H. richardsonii* averaged 1,374, 921, and 679 flowers per population and averaged 32, 28, and 30 inflorescences per population, respectively. Nectar volumes of plants grown in the wild were lowest in parental taxa and highest in the hybrid (Fig. 7A). By contrast, in greenhouse conditions, nectar rewards were lowest in the hybrid taxon and higher in both parents (Fig. 7C), suggesting wild nectar volumes are the result of foraging. We found a significant difference in nectar volumes between *H. americana* var. *americana* and the hybrid (*p* = 0.03), but a lack of difference between the parental taxa (*p =* 0.50), when analyzing only the wild grown *Heuchera* (Table 3). Likewise, under greenhouse conditions, despite the sign change, the parental nectar volumes do not differ among the parents (*p* = 0.79), but a significant difference between *H. americana* var. *americana* and the hybrid is recovered (*p* = 0.02).

**Table 2.** Averaged floral, fruit, and nectar metrics, and fruit set percentages for ingroup host plants. Numbers of populations the averages were calculated from are listed, trailing the column header names, in order of *Heuchera americana* var. *americana, Heuchera americana* var. *hirsuticaulis,* and *Heuchera richardsonii*. Original data from all populations included in this study can be found in Supplemental Tables S3, S4, and S5.

| Host species | Average floral counts | Average inflorescence count | Fruit set |  | Nectar |  |
| --- | --- | --- | --- | --- | --- | --- |
| *Heuchera americana* var. *americana* | 1347.46 | 31.95 | *Fruits* | 28.08 | Wild | 0.78 mm |
|  |  |  | *Aborted* | 14.00 |  | 0.10 uL |
|  |  |  | *Percentage* | 68.94% | Greenhouse | 2.56 mm |
|  |  |  |  |  |  | 0.32 uL |
| *Heuchera americana* var. *hirsuticaulis* | 920.63 | 27.67 | *Fruits* | 26.33 | Wild | 1.11 mm |
|  |  |  | *Aborted* | 36.00 |  | 0.14 uL |
|  |  |  | *Percentage* | 42.72% | Greenhouse | 0.30 mm |
|  |  |  |  |  |  | 0.04 uL |
| *Heuchera richardsonii* | 678.5 | 29.83 | *Fruits* | 16.00 | Wild | 0.16 mm |
|  |  |  | *Aborted* | 9.33 |  | 0.02 uL |
|  |  |  | *Percentage* | 69.60% | Greenhouse | 2.84 mm |
|  |  |  |  |  |  | 0.36 uL |

**Table 3.**
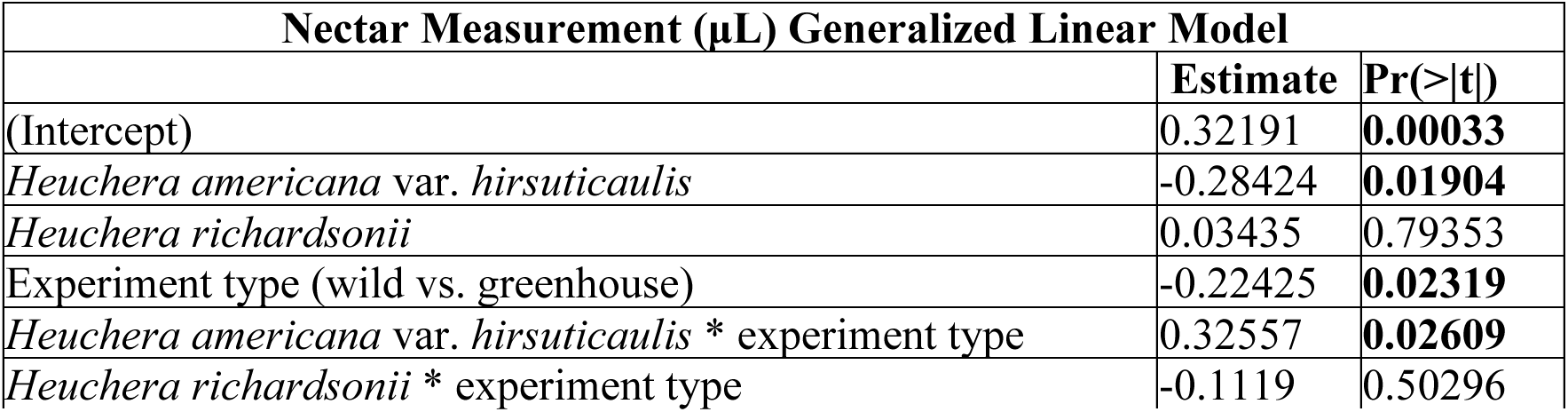
Results from the generalized linear model for nectar measurements. Default comparisons were made to the species *Heuchera americana* var. *americana* and experiment type greenhouse. Significant p-values are in bold.

Fruit sets in relative terms were similar among the parental taxa and lower in the hybrid, averaging 69% for *H. americana* var. *americana*, 43% for the hybrid, and 70% for *H. richardsonii*. Likewise, median fruit set proportions were 80%, 40%, and 75% respectively (Fig 7A). Although our metric of fruit set was substantially lower in the hybrid, fruit set was very variable among populations and an ANOVA indicated no discernible difference in median fruit set proportion between the taxa (*p* = 0.12) similar to results crossing these taxa under artificial conditions (Wells, 1979).

## Discussion

### Floral scent

We centrally sought to test whether conservatism in floral attractants is associated with hybridization. As we anticipated, floral scent profiles measured via TICs were significantly different among species as considered at the level of the genus as a whole, but appreciably similar and statistically indiscernible when evaluating only the hybridizing complex. There appears to be a geographic gradient in trace compounds present in the ranges of the parents *H. americana* and *H. richardsonii*, and particularly the two parental species are distinguished by a more monoterpene- or ketone-based fragrance respectively. However, the VOC makeup is remarkably consistent across the narrower range of the hybrid zone and variant VOC profiles are associated with the peripherals of this range. One interpretation of this proximity based bimodal distribution of trace compound diversity in the hybrid zone, as well as both parents in parts of their range, suggests that the major volatiles responsible for attracting pollinators exhibit evolutionary conservatism in this range, despite the effects of local homogenization or divergence of trace volatiles. The source of this conservatism is unclear, but suggests some type of evolutionary constraint, such as stabilizing selection. Additionally, the localized homogenization of trace volatiles in the hybrid zone could suggest that gene flow is ongoing, but limited in range.

The spring-blooming eastern USA species of *Heuchera* form a clade of seven species with similarly overlapping ranges and near-contemporaneous phenologies, but despite having similar potential contact the remaining five species generally do not exhibit the same propensity for hybridizing with one another as do *H. americana* and *H. richardsonii*. Similarly, outgroups from other areas of the range do not form similar large hybrid zones. Our hypothesis was that more distantly related species tend to possess more divergent floral attractants and therefore that attractants primarily mediate actualized gene flow. Floral volatiles appear to lack significant divergence and therefore are unable to delimit distinct biotic niches between *H. americana* and *H. richardsonii.* Since *Heuchera* comprises primarily narrow ecological specialists (Rosendahl et al., 1936; Wells, 1984; Folk and Freudenstein, 2015; Folk et al., 2018b; Engle-Wrye, 2023; Pantinople et al., 2024) and seeds are limited in dispersal range, limiting a maternal dispersal process (Wells 1984), it seems probable that pollinators are the key limiter of gene flow between these species and that gene flow is realized by the intersection of biotic niche and geographic proximity.

### Pollinator assemblages

Our initial expectation was that the focal plant taxa would be pollination generalists while their insect pollinators may themselves be specialists, such as in the case of the *Heuchera* specialist bee *Colletes aestivalis*. Over the course of two field seasons, however, we found that the hybridizing *Heuchera* specialized in attracting primarily three bee genera as effective pollinators: *Lasioglossum*, *Colletes*, and *Augochlorella*, and thus the plants are highly specialized at the level of pollinator function groups (small bees). *Colletes aestivalis* is an oligolectic *Heuchera* pollinator while the two other *Colletes* seen and the several species of *Lasioglossum* and *Augochlorella* observed are polylectic. These genera accounted for over 96% of all the visitors observed and when they were present, they almost exclusively visited *Heuchera* with very few observations of pollinating other plants present (five occasions from four populations involving *Rosa, Rubus*, *Scrophularia,* and *Tradescantia*) and metabarcoding data indicated only a minor non-*Heuchera* pollen load. Almost all captured insects were female, and they spent much of their time collecting pollen as well as nectaring. Other incidental or more broadly generalist pollinators, like Ichneumonidae and Syrphidae, were observed, but they were inconsistent in their presence, made up only a small percentage of the pollinator assemblage, carried minute to no pollen loads, and visited other non-*Heuchera* species frequently. Further evidence that the three bee genera are the effective pollinators include behavioral observations indicating association with the sexual parts, long visits (as long as several minutes) with both pollen and nectar foraging behavior, visual evidence of movement of pollen, large pollen loads, and metabarcoding data showing that the pollen loads are primarily *Heuchera* and thus that all pollinators were primarily foraging *Heuchera* at the time of capture. Overall evidence indicates that the hybrid complex specializes in attracting, and relies heavily upon, three bee genera (often simultaneously) while only occasionally attracting opportunistic generalist visitors.

During pollination observations, we noticed strong differences in the visible pollen load size present on captured pollinators. We also recorded field observations of the typical saffron- orange *Heuchera* pollen, which was distinct from all other angiosperms observed at the field sites and easily observed on the pollinators. *Heuchera richardsonii*, the hybrid, and *H. americana* var. *americana* all had pollinator assemblages with over 85% *Heuchera* pollen (Fig 8A), suggesting high pollinator fidelity at the time of capture. Every tested visitor except for Syrphidae had a detectable pollen presence identifiable as *Heuchera*. Syrphids and ichneumonids both spent significant amounts of time (e.g., > 1 min) per flower but had poor pollen deposition due to their landing behavior and leg positioning, which was different from bees; only Ichneumonidae carried visually notable *Heuchera* pollen loads. Similarly, many inflorescences were parasitized by aphids (cf. *Nasonovia*), and while these samples had detectable *Heuchera* pollen (Fig. 8 and Fig. 9), these pollen loads were miniscule. This taken in combination with their lack of pollinating behavior and limited dispersal likely indicates a lack of effective pollination. However, these taxa could act as pollinators of hardship, albeit less effective than the primary effective pollinator assemblage.

Although pollinator assemblages by host plant were consistent, suggesting the large morphological difference among parental taxa (Fig. 2) does not relate to visitation directly, there did appear to be a geographic gradient as to which of the three primary pollinator genera predominated any given population’s assemblage; see Figure 3. Southern populations were often primarily visited by *Augochlorella*, while more northern Appalachian populations were visited primarily by *Colletes* or *Lasioglossum*. Most populations west of the Appalachian Mountains and north of Alabama had *Lasioglossum* as their primary pollinator with a few exceptions with more frequent *Colletes* or *Augochlorella* visitations. Within the hybrid zone, *Lasioglossum* was the dominant taxon overall and was present without exception. Overall, there appeared to be three regions of predominating pollinator presences, which may reflect localized availability of the pollinators rather than changes in host plant attractants.

### Flowers, fruit set, and nectar

While numbers of inflorescences per population were consistent across the taxa of the hybrid complex, floral counts varied greatly and corresponded with species differences in flower size. *Heuchera richardsonii* had the largest and fewest flowers, the hybrid was intermediate in both, and *H. americana* had the smallest, but most numerous flower counts. Floral counts did not correlate with successful fruit set, implying that sheer numbers of flowers or population size does not predict fecundity in the study species. Fruit set did not significantly differ among the hybrid and parents, consistent with artificial crossing experiments (Wells, 1979), but exhibited a notable downward trend in the hybrid, which also displayed pollinator reward differences. As can be seen in Table 2, the parental host plants yielded around seven times the nectar volume in greenhouse conditions (∼ 0.3 uL vs 0.04 uL). Because nectar measurements were lowest in greenhouse settings but highest in the wild for the hybrid (indicating low foraging of hybrids) and experimental evidence does not suggest lowered fruit set in this hybrid under equal pollination conditions (Wells, 1979), the lower fruit set proportions seen in the hybrid could instead be associated with poor visitation. Notably, *H. richardsonii* and *H. americana* var. *americana* had both similar fruit set percentages (70% and 69% respectively) and similar visitation averages (169.4 and 119.3 per day per population, respectively), whereas the hybrid only had a fruit set of 43% and had the lowest visitation rate (87.2 per day per population). In sum, fruit set, while not significant, tended to be lower in hybrid *Heuchera* and this appears to be associated with lower visitation rates and significantly inferior pollinator rewards.

### Phylogenetic implications

This is the first population genetic study in *Heuchera*, and thus the first to directly study hybridization in this genus as a population-level process rather than from a phylogenetic point of view. The most important outcome of increased sampling was evidence that different populations within the hybrid zone have differing ancestry. Folk et al. (2017) posited one ancient hybridization event in the history of *H. americana* var. *hirsuticaulis*, but we would expect descendent populations to resolve as monophyletic in this instance. A simplistic interpretation of our PhyloNetworks results is that that *H. americana* is itself a species of hybrid origin with 8% introgression from *H. americana* var. *hirsuticaulis.* However, we interpret the results to indicate that *H. americana* var. *hirsuticaulis* is an ancient hybrid involving ancestral contributions from *H. americana* and *H. richardsonii.* Although PhyloNetworks is sensitive for detecting hybridization involving members of the study, limitations apply to detecting hybrid edges and polarity can be problematic. For example, we limited our test to a maximum of 5 hybridization events (*h_max_*=5) for computational reasons and to avoid overfitting the data, but it is possible, maybe even likely, that there are many more hybridization events throughout the history of this hybrid complex than are computationally feasible for PhyloNetworks to test and which would result in difficult interpretation were they overlapping. More notably, rooting is problematic given species delimitation problems (note the either derived [Fig. 4] or early-diverging [Fig. 5] position of *H. richardsonii* depending on analytical setup). Paraphyly of the species currently recognized as *H. americana* (Folk et al., 2018a) represents a further issue complicating interpretation of gene flow in this wide-ranging and morphologically complex species. Finally, *H. americana* var. *hirsuticaulis* is itself complex, and while in an ordination it appears as expected for a hybrid, it appears to contain two clusters both genetically and geographically closer to the adjacent parent (Fig. 3C). In a future contribution we will revisit species delimitation in the *H. americana* complex, which likely contains several taxa, only some of which engage the hybrid zone.

### Conclusions

To our knowledge this is the first direct investigation of floral attractants in a hybrid-non-hybrid contrast. We found that similar floral scent profiles attract similar pollinator assemblages in the focal *Heuchera* hybrid zone, while floral scent profiles of the outgroup *Heuchera* are divergent and associated with diversified pollinator assemblages. Visitation trends in the hybrid zone are weak and taxonomic IDs (thus morphology) are unrelated to visitation. Our results demonstrate that a lack of divergence in biotic niche is correlated with overlapping pollinator assemblages, identifying that the shape of the plant-pollinator mosaic is a key potential driver of gene flow. We conclude that biotic niche and its interaction with abiotic factors is a major potential factor that may explain disparities in interspecific gene flow between taxa.

## Acknowledgements

This work was funded by NSF DEB-2337784, an American Society of Plant Taxonomists (ASPT) Graduate Research Grant, an International Association for Plant Taxonomy (IAPT) Research Grant, a Society of Systematic Biologists Graduate Research Award, an Oskar H. Zernickow Endowed Fund, a Mississippi State University Strategic Research Initiative (SRI) Student Research & Scholarly Activity Funding Program, and a Mississippi State University Biology Faculty Fund Research Award.

## Notes

### Competing Interest Statement

The authors have declared no competing interest.

