## Supplemental figures and tables for "Investigating the landscape of plant-pollinator interactions in a hybrid zone"

### Supplement

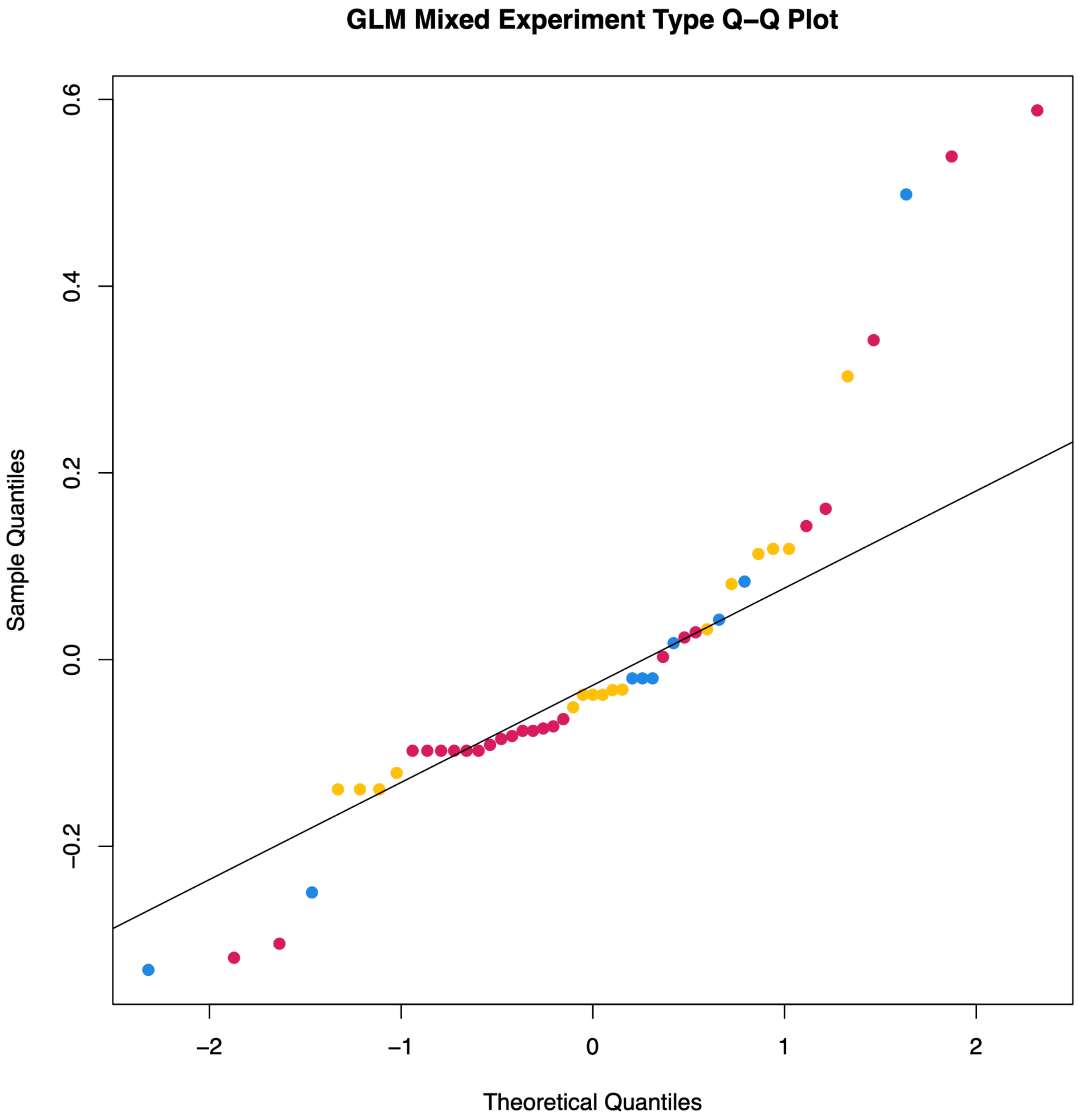

**Fig. S1 |** Q-Q plot of the residuals for the nectar measurements of mixed experimental type (greenhouse vs. wild growing conditions) generalized linear model. Colors of points in the plot match the colors used throughout the paper with red being *H. americana*, yellow being the hybrid, and blue being *H. richardsonii*.

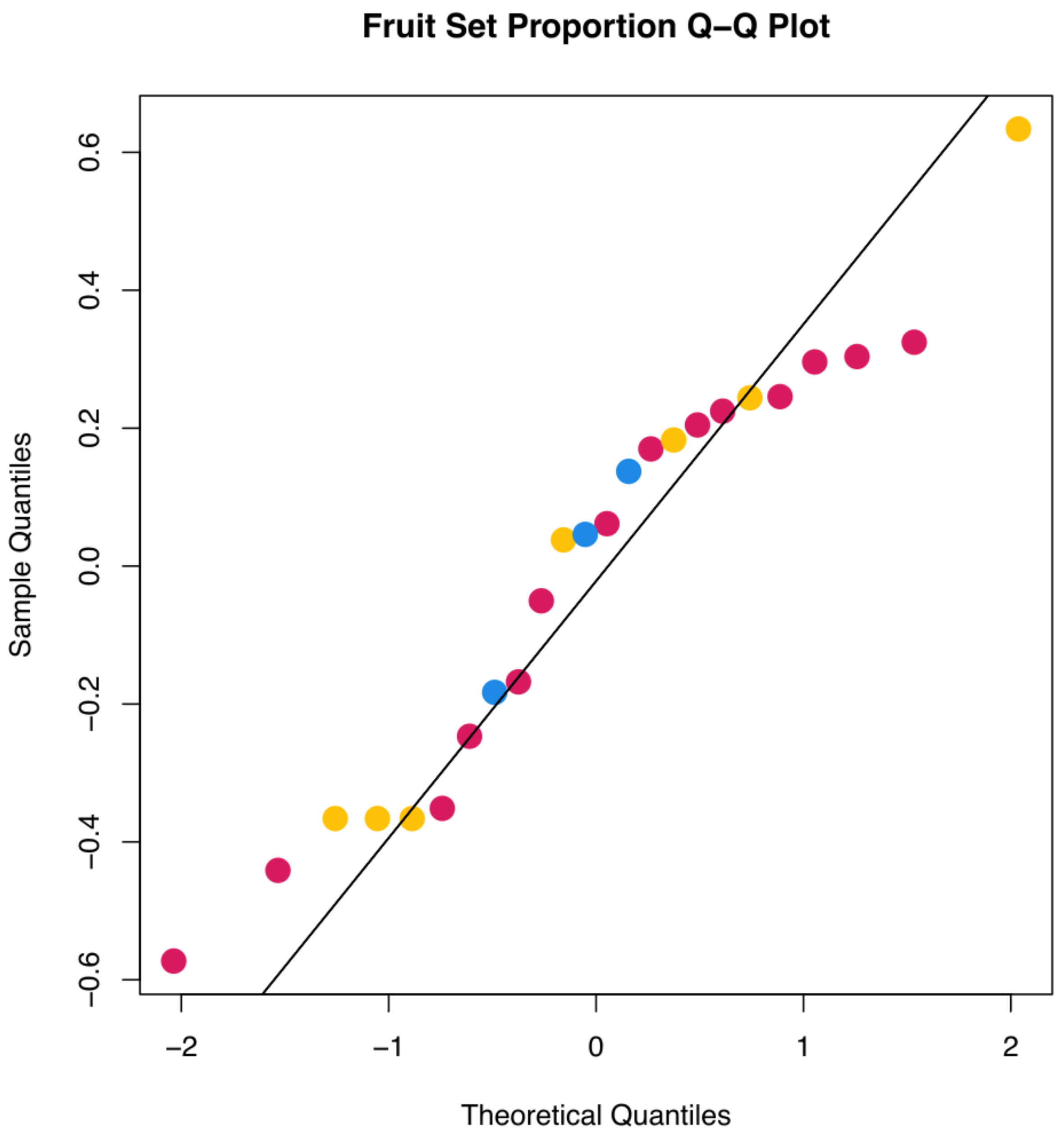

**Fig. S2 |** Fruit set proportion Q-Q plot. Colors of points in the plot match the colors used throughout the paper with red being *H. americana*, yellow being the hybrid, and blue being *H. richardsonii*.

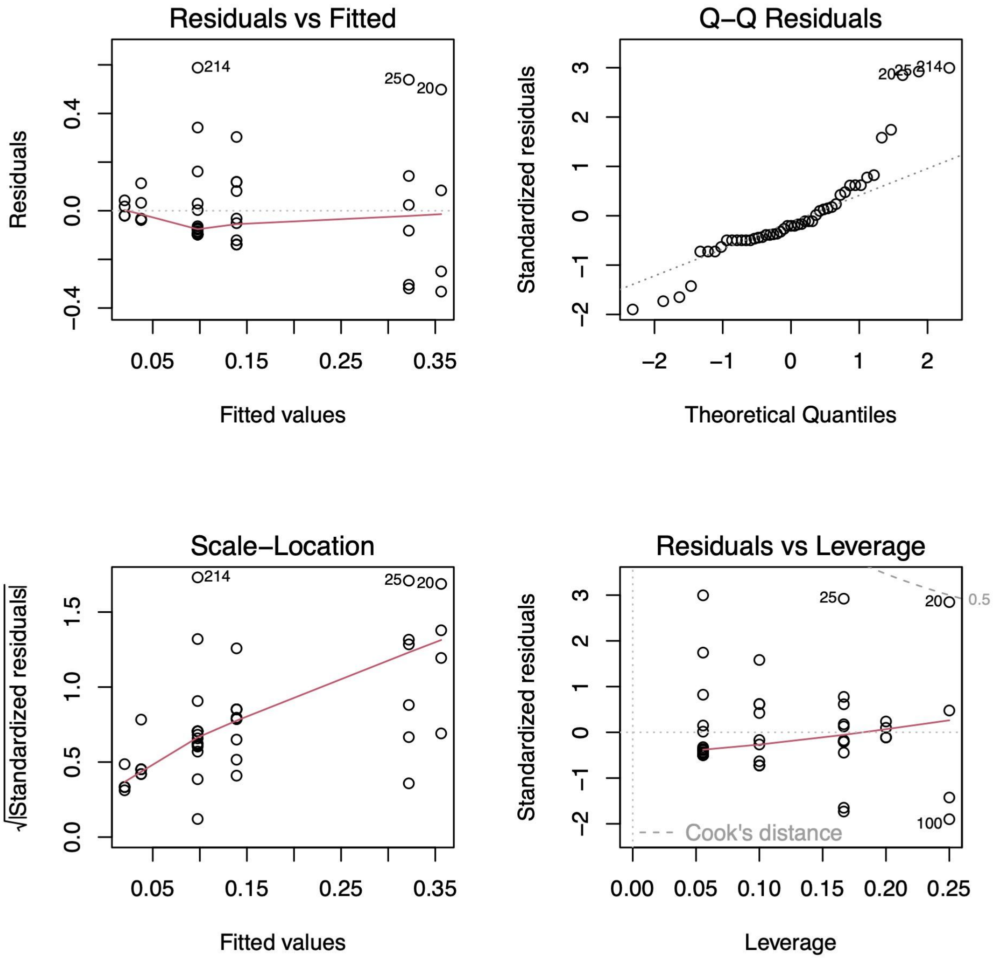

**Fig. S3 |** Two-way ANOVA diagnostic plots for fruit set proportion.

**Table S1.** Accession table of *Heuchera* specimens used for phylogenetic analyses.

| Accession | Morphological assignment | Collector + number | Provider acronym | Provider identifier | Country | State | County | Latitude | Longitude |
|---|---|---|---|---|---|---|---|---|---|
| A1-1 | *Heuchera americana var. americana* | Folk A1 | MISSA | MISSA034877 | USA | AL | Cleburne | 33.725 | -85.600833 |
| A1-2 | *Heuchera americana var. americana* | Folk A1 | MISSA | MISSA034877 | USA | AL | Cleburne | 33.725 | -85.600833 |
| A1-2 | *Heuchera americana var. americana* | Folk A1 | MISSA | MISSA034877 | USA | AL | Cleburne | 33.725 | -85.600833 |
| A1-3 | *Heuchera americana var. americana* | Folk A1 | MISSA | MISSA034877 | USA | AL | Cleburne | 33.725 | -85.600833 |
| A1-4 | *Heuchera americana var. americana* | Folk A1 | MISSA | MISSA034877 | USA | AL | Cleburne | 33.725 | -85.600833 |
| A10-1 | *Heuchera americana var. americana* | Folk A10 | MISSA | MISSA034513 | USA | IN | Floyd | 38.2152778 | -85.906944 |
| A10-2 | *Heuchera americana var. americana* | Folk A10 | MISSA | MISSA034513 | USA | IN | Floyd | 38.2152778 | -85.906944 |
| A10-3 | *Heuchera americana var. americana* | Folk A10 | MISSA | MISSA034513 | USA | IN | Floyd | 38.2152778 | -85.906944 |
| A10-4 | *Heuchera americana var. americana* | Folk A10 | MISSA | MISSA034513 | USA | IN | Floyd | 38.2152778 | -85.906944 |
| A10-5 | *Heuchera americana var. americana* | Folk A10 | MISSA | MISSA034513 | USA | IN | Floyd | 38.2152778 | -85.906944 |
| A10-6 | *Heuchera americana var. americana* | Folk A10 | MISSA | MISSA034513 | USA | IN | Floyd | 38.2152778 | -85.906944 |
| A10-7 | *Heuchera americana var. americana* | Folk A10 | MISSA | MISSA034513 | USA | IN | Floyd | 38.2152778 | -85.906944 |
| A10-8 | *Heuchera americana var. americana* | Folk A10 | MISSA | MISSA034513 | USA | IN | Floyd | 38.2152778 | -85.906944 |
| A11-1 | *Heuchera americana var. americana* | Folk A11 | | No voucher | USA | IN | Crawford | 38.2591667 | -86.461389 |
| A11-2 | *Heuchera americana var. americana* | Folk A11 | | No voucher | USA | IN | Crawford | 38.2591667 | -86.461389 |
| A11-3 | *Heuchera americana var. americana* | Folk A11 | | No voucher | USA | IN | Crawford | 38.2591667 | -86.461389 |
| A11-4 | *Heuchera americana var. americana* | Folk A11 | | No voucher | USA | IN | Crawford | 38.2591667 | -86.461389 |
| A11-5 | *Heuchera americana var. americana* | Folk A11 | | No voucher | USA | IN | Crawford | 38.2591667 | -86.461389 |
| A11-6 | *Heuchera americana var. americana* | Folk A11 | | No voucher | USA | IN | Crawford | 38.2591667 | -86.461389 |

| | | | | | | | | | |
|---|---|---|---|---|---|---|---|---|---|
| A11-7 | *Heuchera americana var. americana* | Folk A11 | | No voucher | USA | IN | Crawford | 38.2591667 | -86.461389 |
| A11-8 | *Heuchera americana var. americana* | Folk A11 | | No voucher | USA | IN | Crawford | 38.2591667 | -86.461389 |
| A13-1 | *Heuchera americana var. hirsuticaulis* | Folk A13 | MISSA | MISSA036288 | USA | IL | Saline | 37.60485 | -88.384667 |
| A13-2 | *Heuchera americana var. hirsuticaulis* | Folk A13 | MISSA | MISSA036288 | USA | IL | Saline | 37.60485 | -88.384667 |
| A13-3 | *Heuchera americana var. hirsuticaulis* | Folk A13 | MISSA | MISSA036288 | USA | IL | Saline | 37.60485 | -88.384667 |
| A13-4 | *Heuchera americana var. hirsuticaulis* | Folk A13 | MISSA | MISSA036288 | USA | IL | Saline | 37.60485 | -88.384667 |
| A13-5 | *Heuchera americana var. hirsuticaulis* | Folk A13 | MISSA | MISSA036288 | USA | IL | Saline | 37.60485 | -88.384667 |
| A13-6 | *Heuchera americana var. hirsuticaulis* | Folk A13 | MISSA | MISSA036288 | USA | IL | Saline | 37.60485 | -88.384667 |
| A13-7 | *Heuchera americana var. hirsuticaulis* | Folk A13 | MISSA | MISSA036288 | USA | IL | Saline | 37.60485 | -88.384667 |
| A13-8 | *Heuchera americana var. hirsuticaulis* | Folk A13 | MISSA | MISSA036288 | USA | IL | Saline | 37.60485 | -88.384667 |
| A13-9 | *Heuchera americana var. hirsuticaulis* | Folk A13 | MISSA | MISSA036288 | USA | IL | Saline | 37.60485 | -88.384667 |
| A14 | *Heuchera americana var. hirsuticaulis* | Folk A14 | MISSA | MISSA036286 | USA | IL | Jersey | 38.9730556 | -90.464444 |
| A15-10 | *Heuchera richardsonii* | Folk A15 | MISSA | MISSA036290 | USA | MO | Camden | 38.1491167 | -92.825311 |
| A15-11 | *Heuchera richardsonii* | Folk A15 | MISSA | MISSA036290 | USA | MO | Camden | 38.1491167 | -92.825311 |
| A15-12 | *Heuchera richardsonii* | Folk A15 | MISSA | MISSA036290 | USA | MO | Camden | 38.1491167 | -92.825311 |
| A15-13 | *Heuchera richardsonii* | Folk A15 | MISSA | MISSA036290 | USA | MO | Camden | 38.1491167 | -92.825311 |
| A15-15 | *Heuchera richardsonii* | Folk A15 | MISSA | MISSA036290 | USA | MO | Camden | 38.1491167 | -92.825311 |
| A15-2 | *Heuchera richardsonii* | Folk A15 | MISSA | MISSA036290 | USA | MO | Camden | 38.1491167 | -92.825311 |
| A15-3 | *Heuchera richardsonii* | Folk A15 | MISSA | MISSA036290 | USA | MO | Camden | 38.1491167 | -92.825311 |
| A15-4 | *Heuchera richardsonii* | Folk A15 | MISSA | MISSA036290 | USA | MO | Camden | 38.1491167 | -92.825311 |
| A15-5 | *Heuchera richardsonii* | Folk A15 | MISSA | MISSA036290 | USA | MO | Camden | 38.1491167 | -92.825311 |
| A15-6 | *Heuchera richardsonii* | Folk A15 | MISSA | MISSA036290 | USA | MO | Camden | 38.1491167 | -92.825311 |
| A15-7 | *Heuchera richardsonii* | Folk A15 | MISSA | MISSA036290 | USA | MO | Camden | 38.1491167 | -92.825311 |
| A15-8 | *Heuchera richardsonii* | Folk A15 | MISSA | MISSA036290 | USA | MO | Camden | 38.1491167 | -92.825311 |

| | | | | | | | | | |
|---|---|---|---|---|---|---|---|---|---|
| A15-9 | *Heuchera richardsonii* | Folk A15 | MISSA | MISSA036290 | USA | MO | Camden | 38.1491167 | -92.825311 |
| A16-1 | *Heuchera richardsonii* | Folk A16 | MISSA | MISSA036549 | USA | WS | Sauk | 43.4175 | -89.726944 |
| A16-10 | *Heuchera richardsonii* | Folk A16 | MISSA | MISSA036549 | USA | WS | Sauk | 43.4175 | -89.726944 |
| A16-11 | *Heuchera richardsonii* | Folk A16 | MISSA | MISSA036549 | USA | WS | Sauk | 43.4175 | -89.726944 |
| A16-2 | *Heuchera richardsonii* | Folk A16 | MISSA | MISSA036549 | USA | WS | Sauk | 43.4175 | -89.726944 |
| A16-3 | *Heuchera richardsonii* | Folk A16 | MISSA | MISSA036549 | USA | WS | Sauk | 43.4175 | -89.726944 |
| A16-4 | *Heuchera richardsonii* | Folk A16 | MISSA | MISSA036549 | USA | WS | Sauk | 43.4175 | -89.726944 |
| A16-6 | *Heuchera richardsonii* | Folk A16 | MISSA | MISSA036549 | USA | WS | Sauk | 43.4175 | -89.726944 |
| A16-8 | *Heuchera richardsonii* | Folk A16 | MISSA | MISSA036549 | USA | WS | Sauk | 43.4175 | -89.726944 |
| A16-9 | *Heuchera richardsonii* | Folk A16 | MISSA | MISSA036549 | USA | WS | Sauk | 43.4175 | -89.726944 |
| A16-9 | *Heuchera richardsonii* | Folk A16 | MISSA | MISSA036549 | USA | WS | Sauk | 43.4175 | -89.726944 |
| A17-1 | *Heuchera richardsonii* | Folk A17 | MISSA | MISSA036545 | USA | WS | Polk | 45.3975 | -92.648056 |
| A17-2 | *Heuchera richardsonii* | Folk A17 | MISSA | MISSA036545 | USA | WS | Polk | 45.3975 | -92.648056 |
| A17-3 | *Heuchera richardsonii* | Folk A17 | MISSA | MISSA036545 | USA | WS | Polk | 45.3975 | -92.648056 |
| A17-4 | *Heuchera richardsonii* | Folk A17 | MISSA | MISSA036545 | USA | WS | Polk | 45.3975 | -92.648056 |
| A17-5 | *Heuchera richardsonii* | Folk A17 | MISSA | MISSA036545 | USA | WS | Polk | 45.3975 | -92.648056 |
| A17-6 | *Heuchera richardsonii* | Folk A17 | MISSA | MISSA036545 | USA | WS | Polk | 45.3975 | -92.648056 |
| A17-7 | *Heuchera richardsonii* | Folk A17 | MISSA | MISSA036545 | USA | WS | Polk | 45.3975 | -92.648056 |
| A17-8 | *Heuchera richardsonii* | Folk A17 | MISSA | MISSA036545 | USA | WS | Polk | 45.3975 | -92.648056 |
| A17-9 | *Heuchera richardsonii* | Folk A17 | MISSA | MISSA036545 | USA | WS | Polk | 45.3975 | -92.648056 |
| A2-3 | *Heuchera americana var. americana* | Folk A2 | MISSA | MISSA034880 | USA | AL | Clay | 33.3705556 | -85.713056 |
| A2-3 | *Heuchera americana var. americana* | Folk A2 | MISSA | MISSA034880 | USA | AL | Clay | 33.3705556 | -85.713056 |
| A2-4 | *Heuchera americana var. americana* | Folk A2 | MISSA | MISSA034880 | USA | AL | Clay | 33.3705556 | -85.713056 |
| A2-4 | *Heuchera americana var. americana* | Folk A2 | MISSA | MISSA034880 | USA | AL | Clay | 33.3705556 | -85.713056 |
| A21-1 | *Heuchera richardsonii* | Folk A21 | MISSA | MISSA036546 | USA | MN | Lake | 47.7205556 | -91.777778 |
| A21-10 | *Heuchera richardsonii* | Folk A21 | MISSA | MISSA036546 | USA | MN | Lake | 47.7205556 | -91.777778 |
| A21-11 | *Heuchera richardsonii* | Folk A21 | MISSA | MISSA036546 | USA | MN | Lake | 47.7205556 | -91.777778 |

| | | | | | | | | | |
|---|---|---|---|---|---|---|---|---|---|
| A21-2 | *Heuchera richardsonii* | Folk A21 | MISSA | MISSA036546 | USA | MN | Lake | 47.7205556 | -91.777778 |
| A21-3 | *Heuchera richardsonii* | Folk A21 | MISSA | MISSA036546 | USA | MN | Lake | 47.7205556 | -91.777778 |
| A21-4 | *Heuchera richardsonii* | Folk A21 | MISSA | MISSA036546 | USA | MN | Lake | 47.7205556 | -91.777778 |
| A21-5 | *Heuchera richardsonii* | Folk A21 | MISSA | MISSA036546 | USA | MN | Lake | 47.7205556 | -91.777778 |
| A21-6 | *Heuchera richardsonii* | Folk A21 | MISSA | MISSA036546 | USA | MN | Lake | 47.7205556 | -91.777778 |
| A21-7 | *Heuchera richardsonii* | Folk A21 | MISSA | MISSA036546 | USA | MN | Lake | 47.7205556 | -91.777778 |
| A21-8 | *Heuchera richardsonii* | Folk A21 | MISSA | MISSA036546 | USA | MN | Lake | 47.7205556 | -91.777778 |
| A21-9 | *Heuchera richardsonii* | Folk A21 | MISSA | MISSA036546 | USA | MN | Lake | 47.7205556 | -91.777778 |
| A22-1 | *Heuchera glomerulata* | Folk A22 | MISSA | MISSA034909 | USA | AZ | Graham | 32.6499094 | -109.81976 |
| A26-1 | *Heuchera sanguinea* | Folk A26 | MISSA | MISSA036297 | USA | AZ | Graham | 32.6359861 | -109.82353 |
| A28-1 | *Heuchera americana var. americana* | Folk A28 | MISSA | MISSA036285 | USA | MS | Tishomingo | 34.58071 | -88.192537 |
| A28-2 | *Heuchera americana var. americana* | Folk A28 | MISSA | MISSA036285 | USA | MS | Tishomingo | 34.58071 | -88.192537 |
| A28-3 | *Heuchera americana var. americana* | Folk A28 | MISSA | MISSA036285 | USA | MS | Tishomingo | 34.58071 | -88.192537 |
| A28-4 | *Heuchera americana var. americana* | Folk A28 | MISSA | MISSA036285 | USA | MS | Tishomingo | 34.58071 | -88.192537 |
| A28-5 | *Heuchera americana var. americana* | Folk A28 | MISSA | MISSA036285 | USA | MS | Tishomingo | 34.58071 | -88.192537 |
| A28-6 | *Heuchera americana var. americana* | Folk A28 | MISSA | MISSA036285 | USA | MS | Tishomingo | 34.58071 | -88.192537 |
| A28-7 | *Heuchera americana var. americana* | Folk A28 | MISSA | MISSA036285 | USA | MS | Tishomingo | 34.58071 | -88.192537 |
| A28-8 | *Heuchera americana var. americana* | Folk A28 | MISSA | MISSA036285 | USA | MS | Tishomingo | 34.58071 | -88.192537 |
| A28-9 | *Heuchera americana var. americana* | Folk A28 | MISSA | MISSA036285 | USA | MS | Tishomingo | 34.58071 | -88.192537 |
| A29-1 | *Heuchera americana var. americana* | Folk A29 | MISSA | MISSA036878 | USA | MS | Tishomingo | 34.9294925 | -88.191126 |
| A29-10 | *Heuchera americana var. americana* | Folk A29 | MISSA | MISSA036902 | USA | MS | Tishomingo | 34.9294925 | -88.191126 |
| A29-2 | *Heuchera americana var. americana* | Folk A29 | MISSA | MISSA036902 | USA | MS | Tishomingo | 34.9294925 | -88.191126 |
| A29-3 | *Heuchera americana var. americana* | Folk A29 | MISSA | MISSA036902 | USA | MS | Tishomingo | 34.9294925 | -88.191126 |

| | | | | | | | | | |
|---|---|---|---|---|---|---|---|---|---|
| A29-4 | *Heuchera americana var. americana* | Folk A29 | MISSA | MISSA036902 | USA | MS | Tishomingo | 34.9294925 | -88.191126 |
| A29-5 | *Heuchera americana var. americana* | Folk A29 | MISSA | MISSA036902 | USA | MS | Tishomingo | 34.9294925 | -88.191126 |
| A29-6 | *Heuchera americana var. americana* | Folk A29 | MISSA | MISSA036902 | USA | MS | Tishomingo | 34.9294925 | -88.191126 |
| A29-7 | *Heuchera americana var. americana* | Folk A29 | MISSA | MISSA036902 | USA | MS | Tishomingo | 34.9294925 | -88.191126 |
| A29-8 | *Heuchera americana var. americana* | Folk A29 | MISSA | MISSA036902 | USA | MS | Tishomingo | 34.9294925 | -88.191126 |
| A29-9 | *Heuchera americana var. americana* | Folk A29 | MISSA | MISSA036902 | USA | MS | Tishomingo | 34.9294925 | -88.191126 |
| A3-1 | *Heuchera americana var. americana* | Folk A3 | MISSA | MISSA034881 | USA | AL | Coosa | 32.9541667 | -86.447222 |
| A3-2 | *Heuchera americana var. americana* | Folk A3 | MISSA | MISSA034881 | USA | AL | Coosa | 32.9541667 | -86.447222 |
| A3-2 | *Heuchera americana var. americana* | Folk A3 | MISSA | MISSA034881 | USA | AL | Coosa | 32.9541667 | -86.447222 |
| A3-3 | *Heuchera americana var. americana* | Folk A3 | MISSA | MISSA034881 | USA | AL | Coosa | 32.9541667 | -86.447222 |
| A3-3 | *Heuchera americana var. americana* | Folk A3 | MISSA | MISSA034881 | USA | AL | Coosa | 32.9541667 | -86.447222 |
| A3-4 | *Heuchera americana var. americana* | Folk A3 | MISSA | MISSA034881 | USA | AL | Coosa | 32.9541667 | -86.447222 |
| A30-1 | Folk A30 | MISSA | MISSA036300 | USA | MS | Tishomingo | 34.930459 | -88.189257 | |
| A31 | *Heuchera americana var. hirsuticaulis* | Engle-Wrye A31 | | No voucher | USA | MS | Panola | 34.408321 | -89.836198 |
| A32-1 | *Heuchera americana var. hirsuticaulis* | Engle-Wrye A32 | MISSA | MISSA036889 | USA | TN | Chester | 35.380564 | -88.832925 |
| A32-10 | *Heuchera americana var. hirsuticaulis* | Engle-Wrye A32 | MISSA | MISSA036889 | USA | TN | Chester | 35.380564 | -88.832925 |
| A32-2 | *Heuchera americana var. hirsuticaulis* | Engle-Wrye A32 | MISSA | MISSA036882 | USA | TN | Chester | 35.380564 | -88.832925 |
| A32-3 | *Heuchera americana var. hirsuticaulis* | Engle-Wrye A32 | MISSA | MISSA036889 | USA | TN | Chester | 35.380564 | -88.832925 |
| A32-4 | *Heuchera americana var. hirsuticaulis* | Engle-Wrye A32 | MISSA | MISSA036889 | USA | TN | Chester | 35.380564 | -88.832925 |
| A32-5 | *Heuchera americana var. hirsuticaulis* | Engle-Wrye A32 | MISSA | MISSA036889 | USA | TN | Chester | 35.380564 | -88.832925 |
| A32-6 | *Heuchera americana var. hirsuticaulis* | Engle-Wrye A32 | MISSA | MISSA036889 | USA | TN | Chester | 35.380564 | -88.832925 |

| | | | | | | | | | |
|---|---|---|---|---|---|---|---|---|---|
| A32-7 | *Heuchera americana var. hirsuticaulis* | Engle-Wrye A32 | MISSA | MISSA036889 | USA | TN | Chester | 35.380564 | -88.832925 |
| A32-8 | *Heuchera americana var. hirsuticaulis* | Engle-Wrye A32 | MISSA | MISSA036889 | USA | TN | Chester | 35.380564 | -88.832925 |
| A32-9 | *Heuchera americana var. hirsuticaulis* | Engle-Wrye A32 | MISSA | MISSA036889 | USA | TN | Chester | 35.380564 | -88.832925 |
| A33-1 | *Heuchera americana var. americana* | Engle-Wrye A33 | MISSA | MISSA036890 | USA | TN | Dickson | 36.036792 | -87.415364 |
| A33-2 | *Heuchera americana var. americana* | Engle-Wrye A33 | MISSA | MISSA036890 | USA | TN | Dickson | 36.036792 | -87.415364 |
| A33-3 | *Heuchera americana var. americana* | Engle-Wrye A33 | MISSA | MISSA036890 | USA | TN | Dickson | 36.036792 | -87.415364 |
| A33-4 | *Heuchera americana var. americana* | Engle-Wrye A33 | MISSA | MISSA036890 | USA | TN | Dickson | 36.036792 | -87.415364 |
| A33-5 | *Heuchera americana var. americana* | Engle-Wrye A33 | MISSA | MISSA036890 | USA | TN | Dickson | 36.036792 | -87.415364 |
| A33-6 | *Heuchera americana var. americana* | Engle-Wrye A33 | MISSA | MISSA036890 | USA | TN | Dickson | 36.036792 | -87.415364 |
| A34-1 | *Heuchera americana* | Engle-Wrye A34 | MISSA | MISSA036922 | USA | TN | Dickson | 36.036792 | -87.328511 |
| A34-10 | *Heuchera americana var. americana* | Engle-Wrye A34 | MISSA | MISSA036922 | USA | TN | Dickson | 36.036792 | -87.328511 |
| A34-2 | *Heuchera americana* | Engle-Wrye A34 | MISSA | MISSA036922 | USA | TN | Dickson | 36.036792 | -87.328511 |
| A34-3 | *Heuchera americana var. americana* | Engle-Wrye A34 | MISSA | MISSA036922 | USA | TN | Dickson | 36.036792 | -87.328511 |
| A34-4 | *Heuchera americana var. americana* | Engle-Wrye A34 | MISSA | MISSA036922 | USA | TN | Dickson | 36.036792 | -87.328511 |
| A34-5 | *Heuchera americana var. americana* | Engle-Wrye A34 | MISSA | MISSA036922 | USA | TN | Dickson | 36.036792 | -87.328511 |
| A34-6 | *Heuchera americana var. americana* | Engle-Wrye A34 | MISSA | MISSA036922 | USA | TN | Dickson | 36.036792 | -87.328511 |
| A34-7 | *Heuchera americana var. americana* | Engle-Wrye A34 | MISSA | MISSA036922 | USA | TN | Dickson | 36.036792 | -87.328511 |
| A34-8 | *Heuchera americana var. americana* | Engle-Wrye A34 | MISSA | MISSA036922 | USA | TN | Dickson | 36.036792 | -87.328511 |
| A34-9 | *Heuchera americana var. americana* | Engle-Wrye A34 | MISSA | MISSA036922 | USA | TN | Dickson | 36.036792 | -87.328511 |
| A35-1 | *Heuchera americana var. americana* | Engle-Wrye A35 | MISSA | MISSA036903 | USA | TN | Dickson | 36.311858 | -87.307933 |
| A35-10 | *Heuchera americana var. americana* | Engle-Wrye A35 | MISSA | MISSA036892 | USA | TN | Dickson | 36.311858 | -87.307933 |

| | | | | | | | | | |
|---|---|---|---|---|---|---|---|---|---|
| A35-2 | *Heuchera americana var. americana* | Engle-Wrye A35 | MISSA | MISSA036892 | USA | TN | Dickson | 36.311858 | -87.307933 |
| A35-3 | *Heuchera americana var. americana* | Engle-Wrye A35 | MISSA | MISSA036892 | USA | TN | Dickson | 36.311858 | -87.307933 |
| A35-4 | *Heuchera americana var. americana* | Engle-Wrye A35 | MISSA | MISSA036892 | USA | TN | Dickson | 36.311858 | -87.307933 |
| A35-5 | *Heuchera americana var. americana* | Engle-Wrye A35 | MISSA | MISSA036892 | USA | TN | Dickson | 36.311858 | -87.307933 |
| A35-6 | *Heuchera americana var. americana* | Engle-Wrye A35 | MISSA | MISSA036892 | USA | TN | Dickson | 36.311858 | -87.307933 |
| A35-7 | *Heuchera americana var. americana* | Engle-Wrye A35 | MISSA | MISSA036892 | USA | TN | Dickson | 36.311858 | -87.307933 |
| A35-8 | *Heuchera americana var. americana* | Engle-Wrye A35 | MISSA | MISSA036892 | USA | TN | Dickson | 36.311858 | -87.307933 |
| A35-9 | *Heuchera americana var. americana* | Engle-Wrye A35 | MISSA | MISSA036892 | USA | TN | Dickson | 36.311858 | -87.307933 |
| A36 | *Heuchera americana var. americana* | Engle-Wrye A36 | MISSA | MISSA036927 | USA | KY | Butler | 37.204169 | -86.736069 |
| A37-5 | *Heuchera missouriensis* | Engle-Wrye A37 | MISSA | MISSA036880 | USA | KY | Butler | 37.204169 | -86.736069 |
| A38-1 | *Heuchera americana var. hirsuticaulis* | Engle-Wrye A38 | MISSA | MISSA036910 | USA | KY | Logan | 36.888572 | -86.832992 |
| A38-2 | *Heuchera americana var. hirsuticaulis* | Engle-Wrye A38 | MISSA | MISSA036909 | USA | KY | Logan | 36.888572 | -86.832992 |
| A39-1 | *Heuchera americana var. hirsuticaulis* | Engle-Wrye A39 | MISSA | MISSA036894 | USA | KY | Trigg | 36.847206 | -88.072117 |
| A39-10 | *Heuchera americana var. hirsuticaulis* | Engle-Wrye A39 | MISSA | MISSA036894 | USA | KY | Trigg | 36.847206 | -88.072117 |
| A39-11 | *Heuchera americana var. hirsuticaulis* | Engle-Wrye A39 | MISSA | MISSA036894 | USA | KY | Trigg | 36.847206 | -88.072117 |
| A39-2 | *Heuchera americana var. hirsuticaulis* | Engle-Wrye A39 | MISSA | MISSA036894 | USA | KY | Trigg | 36.847206 | -88.072117 |
| A39-3 | *Heuchera americana var. hirsuticaulis* | Engle-Wrye A39 | MISSA | MISSA036894 | USA | KY | Trigg | 36.847206 | -88.072117 |
| A39-4 | *Heuchera americana var. hirsuticaulis* | Engle-Wrye A39 | MISSA | MISSA036894 | USA | KY | Trigg | 36.847206 | -88.072117 |
| A39-5 | *Heuchera americana var. hirsuticaulis* | Engle-Wrye A39 | MISSA | MISSA036894 | USA | KY | Trigg | 36.847206 | -88.072117 |
| A39-6 | *Heuchera americana var. hirsuticaulis* | Engle-Wrye A39 | MISSA | MISSA036894 | USA | KY | Trigg | 36.847206 | -88.072117 |

| | | | | | | | | | |
|---|---|---|---|---|---|---|---|---|---|
| A39-7 | *Heuchera americana var. hirsuticaulis* | Engle-Wrye A39 | MISSA | MISSA036894 | USA | KY | Trigg | 36.847206 | -88.072117 |
| A39-8 | *Heuchera americana var. hirsuticaulis* | Engle-Wrye A39 | MISSA | MISSA036894 | USA | KY | Trigg | 36.847206 | -88.072117 |
| A39-9 | *Heuchera americana var. hirsuticaulis* | Engle-Wrye A39 | MISSA | MISSA036894 | USA | KY | Trigg | 36.847206 | -88.072117 |
| A4-1 | *Heuchera americana var. americana* | Folk A4 | MISSA | MISSA034507 | USA | GA | Union | 34.7288889 | -84.082778 |
| A4-2 | *Heuchera americana var. americana* | Folk A4 | MISSA | MISSA034507 | USA | GA | Union | 34.7288889 | -84.082778 |
| A4-2 | *Heuchera americana var. americana* | Folk A4 | MISSA | MISSA034507 | USA | GA | Union | 34.7288889 | -84.082778 |
| A4-3 | *Heuchera americana var. americana* | Folk A4 | MISSA | MISSA034507 | USA | GA | Union | 34.7288889 | -84.082778 |
| A4-4 | *Heuchera americana var. americana* | Folk A4 | MISSA | MISSA034507 | USA | GA | Union | 34.7288889 | -84.082778 |
| A4-5 | *Heuchera americana var. americana* | Folk A4 | MISSA | MISSA034507 | USA | GA | Union | 34.7288889 | -84.082778 |
| A40 | *Heuchera americana var. hirsuticaulis* | Engle-Wrye A40 | MISSA | MISSA036919 | USA | IL | Union | 37.573553 | -89.439867 |
| A40-2 | *Heuchera americana var. hirsuticaulis* | Engle-Wrye A40 | MISSA | MISSA036893 | USA | IL | Union | 37.573553 | -89.439867 |
| A40-3 | *Heuchera americana var. hirsuticaulis* | Engle-Wrye A40 | MISSA | MISSA036919 | USA | IL | Union | 37.573553 | -89.439867 |
| A41-1 | *Heuchera americana var. hirsuticaulis* | Engle-Wrye A41 | MISSA | MISSA036925 | USA | MO | Wayne | 36.966256 | -90.234139 |
| A41-2 | *Heuchera americana var. hirsuticaulis* | Engle-Wrye A41 | MISSA | MISSA036888 | USA | MO | Wayne | 36.966256 | -90.234139 |
| A41-3 | *Heuchera americana var. hirsuticaulis* | Engle-Wrye A41 | MISSA | MISSA036911 | USA | MO | Wayne | 36.966256 | -90.234139 |
| A42-1 | *Heuchera richardsonii* | Engle-Wrye A42 | | no voucher | USA | MO | Jefferson | 38.454586 | -90.6237 |
| A42-2 | *Heuchera richardsonii* | Engle-Wrye A42 | | no voucher | USA | MO | Jefferson | 38.454586 | -90.6237 |
| A42-3 | *Heuchera richardsonii* | Engle-Wrye A42 | | no voucher | USA | MO | Jefferson | 38.454586 | -90.6237 |
| A42-4 | *Heuchera richardsonii* | Engle-Wrye A42 | | no voucher | USA | MO | Jefferson | 38.454586 | -90.6237 |
| A43-1 | *Heuchera richardsonii* | Engle-Wrye A43 | | no voucher | USA | IL | St. Louis | 38.630683 | -90.265731 |
| A43-2 | *Heuchera richardsonii* | Engle-Wrye A43 | | no voucher | USA | IL | St. Louis | 38.630683 | -90.265731 |
| A43-3 | *Heuchera richardsonii* | Engle-Wrye A43 | | no voucher | USA | IL | St. Louis | 38.630683 | -90.265731 |

| | | | | | | | | | |
|---|---|---|---|---|---|---|---|---|---|
| A44-1 | *Heuchera richardsonii* | Engle-Wrye A44 | MISSA | MISSA036881 | USA | MO | Boone | 38.830369 | -92.283972 |
| A44-10 | *Heuchera richardsonii* | Engle-Wrye A44 | MISSA | MISSA036881 | USA | MO | Boone | 38.830369 | -92.283972 |
| A44-2 | *Heuchera richardsonii* | Engle-Wrye A44 | MISSA | MISSA036881 | USA | MO | Boone | 38.830369 | -92.283972 |
| A44-3 | *Heuchera richardsonii* | Engle-Wrye A44 | MISSA | MISSA036881 | USA | MO | Boone | 38.830369 | -92.283972 |
| A44-4 | *Heuchera richardsonii* | Engle-Wrye A44 | MISSA | MISSA036881 | USA | MO | Boone | 38.830369 | -92.283972 |
| A44-5 | *Heuchera richardsonii* | Engle-Wrye A44 | MISSA | MISSA036881 | USA | MO | Boone | 38.830369 | -92.283972 |
| A44-6 | *Heuchera richardsonii* | Engle-Wrye A44 | MISSA | MISSA036881 | USA | MO | Boone | 38.830369 | -92.283972 |
| A44-7 | *Heuchera richardsonii* | Engle-Wrye A44 | MISSA | MISSA036881 | USA | MO | Boone | 38.830369 | -92.283972 |
| A44-8 | *Heuchera richardsonii* | Engle-Wrye A44 | MISSA | MISSA036881 | USA | MO | Boone | 38.830369 | -92.283972 |
| A44-9 | *Heuchera richardsonii* | Engle-Wrye A44 | MISSA | MISSA036881 | USA | MO | Boone | 38.830369 | -92.283972 |
| A45-1 | *Heuchera americana var. hirsuticaulis* | Engle-Wrye A45 | MISSA | MISSA036897 | USA | AR | Washington | 36.065467 | -94.13855 |
| A45-10 | *Heuchera americana var. hirsuticaulis* | Engle-Wrye A45 | MISSA | MISSA036897 | USA | AR | Washington | 36.065467 | -94.13855 |
| A45-2 | *Heuchera americana var. hirsuticaulis* | Engle-Wrye A45 | MISSA | MISSA036897 | USA | AR | Washington | 36.065467 | -94.13855 |
| A45-3 | *Heuchera americana var. hirsuticaulis* | Engle-Wrye A45 | MISSA | MISSA036897 | USA | AR | Washington | 36.065467 | -94.13855 |
| A45-4 | *Heuchera americana var. hirsuticaulis* | Engle-Wrye A45 | MISSA | MISSA036897 | USA | AR | Washington | 36.065467 | -94.13855 |
| A45-5 | *Heuchera americana var. hirsuticaulis* | Engle-Wrye A45 | MISSA | MISSA036897 | USA | AR | Washington | 36.065467 | -94.13855 |
| A45-6 | *Heuchera americana var. hirsuticaulis* | Engle-Wrye A45 | MISSA | MISSA036897 | USA | AR | Washington | 36.065467 | -94.13855 |
| A45-7 | *Heuchera americana var. hirsuticaulis* | Engle-Wrye A45 | MISSA | MISSA036897 | USA | AR | Washington | 36.065467 | -94.13855 |
| A45-8 | *Heuchera americana var. hirsuticaulis* | Engle-Wrye A45 | MISSA | MISSA036897 | USA | AR | Washington | 36.065467 | -94.13855 |
| A45-9 | *Heuchera americana var. hirsuticaulis* | Engle-Wrye A45 | MISSA | MISSA036897 | USA | AR | Washington | 36.065467 | -94.13855 |
| A46-1 | *Heuchera americana var. hirsuticaulis* | Engle-Wrye A46 | MISSA | MISSA036920 | USA | AR | Washington | 35.994786 | -94.132508 |
| A46-2 | *Heuchera americana var. hirsuticaulis* | Engle-Wrye A46 | MISSA | MISSA036924 | USA | AR | Washington | 35.994786 | -94.132508 |
| A46-3 | *Heuchera americana var. hirsuticaulis* | Engle-Wrye A46 | MISSA | MISSA036899 | USA | AR | Washington | 35.994786 | -94.132508 |

| | | | | | | | | | |
|---|---|---|---|---|---|---|---|---|---|
| A46-4 | *Heuchera americana var. hirsuticaulis* | Engle-Wrye A46 | MISSA | MISSA036899 | USA | AR | Washington | 35.994786 | -94.132508 |
| A46-5 | *Heuchera americana var. hirsuticaulis* | Engle-Wrye A46 | MISSA | MISSA036899 | USA | AR | Washington | 35.994786 | -94.132508 |
| A46-6 | *Heuchera americana var. hirsuticaulis* | Engle-Wrye A46 | MISSA | MISSA036899 | USA | AR | Washington | 35.994786 | -94.132508 |
| A46-7 | *Heuchera americana var. hirsuticaulis* | Engle-Wrye A46 | MISSA | MISSA036899 | USA | AR | Washington | 35.994786 | -94.132508 |
| A46-8 | *Heuchera americana var. hirsuticaulis* | Engle-Wrye A46 | MISSA | MISSA036899 | USA | AR | Washington | 35.994786 | -94.132508 |
| A47-1 | *Heuchera americana var. hirsuticaulis* | Engle-Wrye A47 | MISSA | MISSA036915 | USA | AR | Washington | 35.996722 | -94.129067 |
| A47-2 | *Heuchera americana var. hirsuticaulis* | Engle-Wrye A47 | MISSA | MISSA036914 | USA | AR | Washington | 35.996722 | -94.129067 |
| A47-3 | *Heuchera americana var. hirsuticaulis* | Engle-Wrye A47 | MISSA | MISSA036896 | USA | AR | Washington | 35.996722 | -94.129067 |
| A47-4 | *Heuchera americana var. hirsuticaulis* | Engle-Wrye A47 | MISSA | MISSA036915 | USA | AR | Washington | 35.996722 | -94.129067 |
| A47-5 | *Heuchera americana var. hirsuticaulis* | Engle-Wrye A47 | MISSA | MISSA036918 | USA | AR | Washington | 35.996722 | -94.129067 |
| A48 | *Heuchera americana var. hirsuticaulis* | Engle-Wrye A48 | MISSA | MISSA036883 | USA | AR | Pope | 35.304633 | -93.165575 |
| A49-1 | *Heuchera americana var. hirsuticaulis* | Engle-Wrye A49 | MISSA | MISSA036921 | USA | AR | Faulkner | 35.074283 | -92.538006 |
| A49-10 | *Heuchera americana var. hirsuticaulis* | Engle-Wrye A49 | MISSA | MISSA036901 | USA | AR | Faulkner | 35.074283 | -92.538006 |
| A49-11 | *Heuchera americana var. hirsuticaulis* | Engle-Wrye A49 | MISSA | MISSA036895 | USA | AR | Faulkner | 35.074283 | -92.538006 |
| A49-12 | *Heuchera americana var. hirsuticaulis* | Engle-Wrye A49 | MISSA | MISSA036901 | USA | AR | Faulkner | 35.074283 | -92.538006 |
| A49-2 | *Heuchera americana var. hirsuticaulis* | Engle-Wrye A49 | MISSA | MISSA036901 | USA | AR | Faulkner | 35.074283 | -92.538006 |
| A49-3 | *Heuchera americana var. hirsuticaulis* | Engle-Wrye A49 | MISSA | MISSA036901 | USA | AR | Faulkner | 35.074283 | -92.538006 |
| A49-4 | *Heuchera americana var. hirsuticaulis* | Engle-Wrye A49 | MISSA | MISSA036901 | USA | AR | Faulkner | 35.074283 | -92.538006 |
| A49-5 | *Heuchera americana var. hirsuticaulis* | Engle-Wrye A49 | MISSA | MISSA036901 | USA | AR | Faulkner | 35.074283 | -92.538006 |
| A49-6 | *Heuchera americana var. hirsuticaulis* | Engle-Wrye A49 | MISSA | MISSA036901 | USA | AR | Faulkner | 35.074283 | -92.538006 |

| | | | | | | | | | |
|---|---|---|---|---|---|---|---|---|---|
| A49-7 | *Heuchera americana var. hirsuticaulis* | Engle-Wrye A49 | MISSA | MISSA036901 | USA | AR | Faulkner | 35.074283 | -92.538006 |
| A49-8 | *Heuchera americana var. hirsuticaulis* | Engle-Wrye A49 | MISSA | MISSA036901 | USA | AR | Faulkner | 35.074283 | -92.538006 |
| A49-9 | *Heuchera americana var. hirsuticaulis* | Engle-Wrye A49 | MISSA | MISSA036901 | USA | AR | Faulkner | 35.074283 | -92.538006 |
| A5-1 | *Heuchera villosa var. villosa* | Folk A5 | MISSA | MISSA034894 | USA | GA | Lumpkin | 34.6772222 | -84 |
| A50-1 | *Heuchera caroliniana* | Folk A50 | MISSA | MISSA036277 | USA | SC | Pickens | 34.993338 | -80.081078 |
| A50-2 | *Heuchera caroliniana* | Folk A50 | MISSA | MISSA036277 | USA | SC | Pickens | 34.993338 | -80.081078 |
| A50-4 | *Heuchera caroliniana* | Folk A50 | MISSA | MISSA036277 | USA | SC | Pickens | 34.993338 | -80.081078 |
| A53-1 | *Heuchera americana var. americana* | Engle-Wrye A53 | | no voucher | USA | AL | Cleburne | 33.693964 | -85.55944 |
| A53-10 | *Heuchera americana var. americana* | Engle-Wrye A53 | | no voucher | USA | AL | Cleburne | 33.693964 | -85.55944 |
| A53-2 | *Heuchera americana var. americana* | Engle-Wrye A53 | | no voucher | USA | AL | Cleburne | 33.693964 | -85.55944 |
| A53-3 | *Heuchera americana var. americana* | Engle-Wrye A53 | | no voucher | USA | AL | Cleburne | 33.693964 | -85.55944 |
| A53-4 | *Heuchera americana var. americana* | Engle-Wrye A53 | | no voucher | USA | AL | Cleburne | 33.693964 | -85.55944 |
| A53-5 | *Heuchera americana var. americana* | Engle-Wrye A53 | | no voucher | USA | AL | Cleburne | 33.693964 | -85.55944 |
| A53-6 | *Heuchera americana var. americana* | Engle-Wrye A53 | | no voucher | USA | AL | Cleburne | 33.693964 | -85.55944 |
| A53-7 | *Heuchera americana var. americana* | Engle-Wrye A53 | | no voucher | USA | AL | Cleburne | 33.693964 | -85.55944 |
| A53-8 | *Heuchera americana var. americana* | Engle-Wrye A53 | | no voucher | USA | AL | Cleburne | 33.693964 | -85.55944 |
| A53-9 | *Heuchera americana var. americana* | Engle-Wrye A53 | | no voucher | USA | AL | Cleburne | 33.693964 | -85.55944 |
| A54-1 | *Heuchera americana var. americana* | Engle-Wrye A54 | | no voucher | USA | AL | Shelby | 33.369119 | -86.659483 |
| A54-2 | *Heuchera americana var. americana* | Engle-Wrye A54 | | no voucher | USA | AL | Shelby | 33.369119 | -86.659483 |
| A54-3 | *Heuchera americana var. americana* | Engle-Wrye A54 | | no voucher | USA | AL | Shelby | 33.369119 | -86.659483 |
| A55-1 | *Heuchera americana var. heteradenia* | Folk A55 | MISSA | MISSA037100 | USA | TN | Polk | 35.0996176 | -84.555909 |

| | | | | | | | | | |
|---|---|---|---|---|---|---|---|---|---|
| A55-2 | *Heuchera americana var. heteradenia* | Folk A55 | MISSA | MISSA037100 | USA | TN | Polk | 35.0996176 | -84.555909 |
| A55-3 | *Heuchera americana var. heteradenia* | Folk A55 | MISSA | MISSA037100 | USA | TN | Polk | 35.0996176 | -84.555909 |
| A55-4 | *Heuchera americana var. heteradenia* | Folk A55 | MISSA | MISSA037100 | USA | TN | Polk | 35.0996176 | -84.555909 |
| A55-5 | *Heuchera americana var. heteradenia* | Folk A55 | MISSA | MISSA037100 | USA | TN | Polk | 35.0996176 | -84.555909 |
| A55-6 | *Heuchera americana var. heteradenia* | Folk A55 | MISSA | MISSA037100 | USA | TN | Polk | 35.0996176 | -84.555909 |
| A56-1 | *Heuchera americana var. americana* | Engle-Wrye A56 | | no voucher | USA | NC | Cherokee | 35.1230527 | -83.989823 |
| A56-2 | *Heuchera americana var. americana* | Engle-Wrye A56 | | no voucher | USA | NC | Cherokee | 35.1230527 | -83.989823 |
| A56-3 | *Heuchera americana var. americana* | Engle-Wrye A56 | | no voucher | USA | NC | Cherokee | 35.1230527 | -83.989823 |
| A56-4 | *Heuchera americana var. americana* | Engle-Wrye A56 | | no voucher | USA | NC | Cherokee | 35.1230527 | -83.989823 |
| A56-5 | *Heuchera americana var. americana* | Engle-Wrye A56 | | no voucher | USA | NC | Cherokee | 35.1230527 | -83.989823 |
| A56-6 | *Heuchera americana var. americana* | Engle-Wrye A56 | | no voucher | USA | NC | Cherokee | 35.1230527 | -83.989823 |
| A56-7 | *Heuchera americana var. americana* | Engle-Wrye A56 | | no voucher | USA | NC | Cherokee | 35.1230527 | -83.989823 |
| A57 | *Heuchera americana var. americana* | Engle-Wrye A57 | | no voucher | USA | NC | Swain | 35.4648505 | -83.433531 |
| A58-1 | *Heuchera americana var. americana* | Engle-Wrye A58 | | no voucher | USA | NC | Jackson | 35.248278 | -83.086306 |
| A58-2 | *Heuchera americana var. americana* | Engle-Wrye A58 | | no voucher | USA | NC | Jackson | 35.248278 | -83.086306 |
| A58-3 | *Heuchera americana var. americana* | Engle-Wrye A58 | | no voucher | USA | NC | Jackson | 35.248278 | -83.086306 |
| A58-4 | *Heuchera americana var. americana* | Engle-Wrye A58 | | no voucher | USA | NC | Jackson | 35.248278 | -83.086306 |
| A59-1 | *Heuchera americana var. hispida* | Engle-Wrye A59 | MISSA | MISSA037098 | USA | NC | McDowell | 35.735357 | -82.116329 |
| A59-2 | *Heuchera americana var. hispida* | Engle-Wrye A59 | MISSA | MISSA037098 | USA | NC | McDowell | 35.735357 | -82.116329 |
| A59-3 | *Heuchera americana var. hispida* | Engle-Wrye A59 | MISSA | MISSA037098 | USA | NC | McDowell | 35.735357 | -82.116329 |

| | | | | | | | | | |
|---|---|---|---|---|---|---|---|---|---|
| A59-4 | *Heuchera americana var. hispida* | Engle-Wrye A59 | MISSA | MISSA037098 | USA | NC | McDowell | 35.735357 | -82.116329 |
| A59-5 | *Heuchera americana var. hispida* | Engle-Wrye A59 | MISSA | MISSA037098 | USA | NC | McDowell | 35.735357 | -82.116329 |
| A59-6 | *Heuchera americana var. hispida* | Engle-Wrye A59 | MISSA | MISSA037098 | USA | NC | McDowell | 35.735357 | -82.116329 |
| A59-7 | *Heuchera americana var. hispida* | Engle-Wrye A59 | MISSA | MISSA037098 | USA | NC | McDowell | 35.735357 | -82.116329 |
| A59-8 | *Heuchera americana var. hispida* | Engle-Wrye A59 | MISSA | MISSA037098 | USA | NC | McDowell | 35.735357 | -82.116329 |
| A59-9 | *Heuchera americana var. hispida* | Engle-Wrye A59 | MISSA | MISSA037098 | USA | NC | McDowell | 35.735357 | -82.116329 |
| A6-2 | *Heuchera americana var. americana* | Folk A6 | MISSA | MISSA034509 | USA | GA | White | 34.6194444 | -83.792222 |
| A6-3 | *Heuchera americana var. americana* | Folk A6 | MISSA | MISSA034509 | USA | GA | White | 34.6194444 | -83.792222 |
| A6-3 | *Heuchera americana var. americana* | Folk A6 | MISSA | MISSA034509 | USA | GA | White | 34.6194444 | -83.792222 |
| A60-1 | *Heuchera americana var. americana* | Engle-Wrye A60 | MISSA | no voucher | USA | NC | Alexander | 35.966166 | -81.114492 |
| A60-2 | *Heuchera americana var. americana* | Engle-Wrye A60 | MISSA | no voucher | USA | NC | Alexander | 35.966166 | -81.114492 |
| A60-3 | *Heuchera americana var. americana* | Engle-Wrye A60 | MISSA | no voucher | USA | NC | Alexander | 35.966166 | -81.114492 |
| A60-4 | *Heuchera americana var. americana* | Engle-Wrye A60 | MISSA | no voucher | USA | NC | Alexander | 35.966166 | -81.114492 |
| A62-1 | *Heuchera americana var. heteradenia* | Engle-Wrye A62 | MISSA | MISSA037106 | USA | TN | Sevier | 35.695778 | -83.389556 |
| A62-2 | *Heuchera americana var. heteradenia* | Engle-Wrye A62 | MISSA | MISSA037106 | USA | TN | Sevier | 35.695778 | -83.389556 |
| A62-3 | *Heuchera americana var. heteradenia* | Engle-Wrye A62 | MISSA | MISSA037106 | USA | TN | Sevier | 35.695778 | -83.389556 |
| A63-1 | *Heuchera americana var. heteradenia* | Engle-Wrye A63 | MISSA | MISSA037105 | USA | TN | Sevier | 35.695778 | -83.389556 |
| A63-10 | *Heuchera americana var. heteradenia* | Engle-Wrye A63 | MISSA | MISSA037105 | USA | TN | Sevier | 35.695778 | -83.389556 |
| A63-2 | *Heuchera americana var. heteradenia* | Engle-Wrye A63 | MISSA | MISSA037105 | USA | TN | Sevier | 35.695778 | -83.389556 |
| A63-3 | *Heuchera americana var. heteradenia* | Engle-Wrye A63 | MISSA | MISSA037105 | USA | TN | Sevier | 35.695778 | -83.389556 |

| | | | | | | | | | |
|---|---|---|---|---|---|---|---|---|---|
| A63-4 | *Heuchera americana var. heteradenia* | Engle-Wrye A63 | MISSA | MISSA037105 | USA | TN | Sevier | 35.695778 | -83.389556 |
| A63-5 | *Heuchera americana var. heteradenia* | Engle-Wrye A63 | MISSA | MISSA037105 | USA | TN | Sevier | 35.695778 | -83.389556 |
| A63-6 | *Heuchera americana var. heteradenia* | Engle-Wrye A63 | MISSA | MISSA037105 | USA | TN | Sevier | 35.695778 | -83.389556 |
| A63-7 | *Heuchera americana var. heteradenia* | Engle-Wrye A63 | MISSA | MISSA037105 | USA | TN | Sevier | 35.695778 | -83.389556 |
| A63-8 | *Heuchera americana var. heteradenia* | Engle-Wrye A63 | MISSA | MISSA037105 | USA | TN | Sevier | 35.695778 | -83.389556 |
| A63-9 | *Heuchera americana var. heteradenia* | Engle-Wrye A63 | MISSA | MISSA037105 | USA | TN | Sevier | 35.695778 | -83.389556 |
| A7-1 | *Heuchera americana var. americana* | Folk A7 | MISSA | To be processed | USA | SC | Abbeville | 34.0972222 | -82.351389 |
| A7-2 | *Heuchera americana var. americana* | Folk A7 | MISSA | To be processed | USA | SC | Abbeville | 34.0972222 | -82.351389 |
| A7-3 | *Heuchera americana var. americana* | Folk A7 | MISSA | To be processed | USA | SC | Abbeville | 34.0972222 | -82.351389 |
| A7-4 | *Heuchera americana var. americana* | Folk A7 | MISSA | To be processed | USA | SC | Abbeville | 34.0972222 | -82.351389 |
| A7-4 | *Heuchera americana var. americana* | Folk A7 | MISSA | To be processed | USA | SC | Abbeville | 34.0972222 | -82.351389 |
| A8 | *Heuchera americana var. americana* | Folk A8 | MISSA | MISSA036643 | USA | GA | Jasper | 33.2555556 | -83.6825 |
| A9-1 | *Heuchera americana var. hirsuticaulis* | Folk A9 | MISSA | MISSA034515 | USA | IN | Warren | 40.3380556 | -87.316389 |
| A9-3 | *Heuchera americana var. hirsuticaulis* | Folk A9 | MISSA | MISSA034515 | USA | IN | Warren | 40.3380556 | -87.316389 |
| A9-4 | *Heuchera americana var. hirsuticaulis* | Folk A9 | MISSA | MISSA034515 | USA | IN | Warren | 40.3380556 | -87.316389 |
| E1 | *Heuchera americana var. americana* | Bowers 13612 | UNA | UNA00034487 | USA | AL | Shelby | 33.183991 | -86.999559 |
| E12 | *Heuchera americana var. americana* | Williams s.n. | UNA | UNA00027468 | USA | AL | Jefferson | 33.5228184 | -86.916451 |
| E1415 | *Heuchera americana var. americana* | Radford 220808 | NCU | NCU00180859 | USA | SC | Abbeville | 34.09611 | -82.344593 |
| E1416 | *Heuchera americana var. americana* | Ahles 56273 | NCU | NCU00180860 | USA | SC | Aiken | 33.542909 | -81.99475 |
| E1419 | *Heuchera americana var. americana* | Nelson 7394 | NCU | NCU00180863 | USA | SC | Cherokee | 33.1595951 | -79.907046 |

| | | | | | | | | | |
|---|---|---|---|---|---|---|---|---|---|
| E1420 | *Heuchera americana var. americana* | Haesloop 26792 | NCU | NCU00180864 | USA | SC | Cherokee | 35.13592 | -81.593086 |
| E1421 | *Heuchera americana var. americana* | Leonard 4250 | NCU | NCU00180865 | USA | SC | Dorchester | 35.0372626 | -81.648076 |
| E1422 | *Heuchera americana var. americana* | Radford 22536 | NCU | NCU00180866 | USA | SC | Edgefield | 33.0756291 | -80.343113 |
| E1423 | *Heuchera americana var. americana* | Bell 7189 | NCU | NCU00180867 | USA | SC | Fairfield | 34.51148 | -81.297873 |
| E1426 | *Heuchera americana var. americana* | Radford 22388 | NCU | NCU00180851 | USA | SC | McCormick | 34.4646451 | -81.994404 |
| E1427 | *Heuchera americana var. americana* | Bell 7024 | NCU | NCU00180852 | USA | SC | Newberry | 33.8795325 | -82.279772 |
| E1429 | *Heuchera americana var. americana* | Knox 100 | NCU | NCU00180854 | USA | SC | Pickens | 34.8882592 | -82.719382 |
| E1430 | *Heuchera americana var. americana* | Freeman 57187 | NCU | NCU00180855 | USA | SC | Pickens | 34.8882592 | -82.719382 |
| E1432 | *Heuchera americana var. americana* | Bell 8451 | NCU | NCU00180857 | USA | SC | Union | 34.9498007 | -81.932016 |
| E1435 | *Heuchera caroliniana* | Ahles 27366 | NCU | NCU00053744 | USA | SC | Lancaster | 34.6859896 | -81.154507 |
| E1436 | *Heuchera caroliniana* | Wells 3310 | NCU | NCU00053745 | USA | SC | Lancaster | 34.6628067 | -80.700546 |
| E1438 | *Heuchera caroliniana* | Smith 67 | NCU | NCU00053743 | USA | SC | Darlington | 34.7622221 | -83.109965 |
| E1457 | *Heuchera americana var. americana* | Furr 582 | NCU | NCU00180880 | USA | KY | Bell | 33.762884 | -83.740416 |
| E1459 | *Heuchera americana var. hirsuticaulis* | Wilson 234 | NCU | NCU00180881 | USA | KY | Carlisle | 36.848 | -89.085 |
| E1469 | *Heuchera americana var. americana* | Furr 586 | NCU | NCU00180898 | USA | TN | Claiborne | 35.6719722 | -83.931447 |
| E1470 | *Heuchera americana var. americana* | Sharp 23198 | NCU | NCU00180897 | USA | TN | Coffee | 36.4673161 | -83.678638 |
| E1472 | *Heuchera americana var. americana* | Shaw s.n. | NCU | NCU00439719 | USA | TN | Fentress | 36.188076 | -85.002043 |
| E1473 | *Heuchera americana var. americana* | Lazell s.n. | NCU | NCU00180894 | USA | TN | Franklin | 36.360098 | -84.926231 |
| E1474 | *Heuchera americana var. americana* | Furr 337 | NCU | NCU00180900 | USA | TN | Grainger | 35.925206 | -86.868942 |
| E1475 | *Heuchera americana var. americana* | Clark 1760 | NCU | NCU00181081 | USA | TN | Grundy | 36.2810821 | -83.510702 |
| E1476 | *Heuchera americana var. americana* | Furr 366 | NCU | NCU00181082 | USA | TN | Hawkins | 35.3697894 | -85.710661 |

| | | | | | | | | | |
|---|---|---|---|---|---|---|---|---|---|
| E1477 | *Heuchera americana var. americana* | Wells 3286 | NCU | NCU00181083 | USA | TN | Jefferson | 36.429878 | -82.95689 |
| E1482 | *Heuchera americana var. americana* | Hess 1150 | NCU | NCU00181090 | USA | TN | Monroe | 35.4397238 | -84.239964 |
| E1486 | *Heuchera americana var. americana* | Furr 292 | NCU | NCU00181093 | USA | TN | Swain | 35.84857 | -84.522552 |
| E1490 | *Heuchera americana var. americana* | Wells 3292 | NCU | NCU00181097 | USA | VA | Botetourt | 37.5520825 | -79.802957 |
| E1491 | *Heuchera americana var. americana* | Kirkland 23220 | NCU | NCU00181098 | USA | VA | Brunswick | 37.5520825 | -79.802957 |
| E1492 | *Heuchera americana var. americana* | Ramsey 7535 | NCU | NCU00181099 | USA | VA | Buckingham | 36.7788996 | -77.867088 |
| E1493 | *Heuchera americana var. americana* | Howell 16501 | NCU | NCU00181100 | USA | VA | Buckingham | 37.5558826 | -78.554717 |
| E1494 | *Heuchera americana var. americana* | Ware 4210 | NCU | NCU00181101 | USA | VA | Charles City | 37.5558826 | -78.554717 |
| E1495 | *Heuchera americana var. americana* | Bradley 6826 | NCU | NCU00181104 | USA | VA | Fairfax | 37.3705777 | -77.06051 |
| E1496 | *Heuchera americana var. americana* | Diggs 226 | NCU | NCU00181105 | USA | VA | Fluvanna | 38.8462236 | -77.306373 |
| E1497 | *Heuchera pubescens* | Weiboldt 8150 | NCU | NCU00181106 | USA | VA | Franklin | 36.838161 | -79.727771 |
| E1498 | *Heuchera americana var. americana* | Wells 3363 | NCU | NCU00181107 | USA | VA | Giles | 37.0527719 | -79.881309 |
| E1499 | *Heuchera americana var. americana* | Kimsey 122 | NCU | NCU00181108 | USA | VA | Goochland | 37.3088528 | -80.709878 |
| E1500 | *Heuchera americana var. americana* | Massey 4014 | NCU | NCU00181111 | USA | VA | Halifax | 37.7204342 | -77.883798 |
| E1501 | *Heuchera americana var. americana* | Ramsey 4173 | NCU | NCU00181109 | USA | VA | Halifax | 36.747244 | -78.942563 |
| E1502 | *Heuchera americana var. americana* | Boufford 13886 | NCU | NCU00181110 | USA | VA | Halifax | 36.747244 | -78.942563 |
| E1503 | *Heuchera americana var. americana* | Wells 3291 | NCU | NCU00181113 | USA | VA | Hanover | 36.747244 | -78.942563 |
| E1504 | *Heuchera americana var. americana* | Dougherty s.n. | NCU | NCU00181115 | USA | VA | Richmond City | 37.744783 | -77.446417 |
| E1505 | *Heuchera americana var. americana* | Wells 3294 | NCU | NCU00181116 | USA | VA | Isle of Wright | 37.034167 | -76.619444 |
| E1506 | *Heuchera americana var. americana* | Borans 201 | NCU | NCU00181118 | USA | VA | James City | 37.271705 | -76.714104 |

| | | | | | | | | | |
|---|---|---|---|---|---|---|---|---|---|
| E1507 | *Heuchera americana var. americana* | Gillespie 635 | NCU | NCU00181122 | USA | VA | New Kent | 37.581311 | -77.062158 |
| E1508 | *Heuchera americana var. americana* | Mikula 5348 | NCU | NCU00181123 | USA | VA | Page | 37.4974701 | -76.999045 |
| E1510 | *Heuchera americana var. americana* | Corcoran 296 | NCU | NCU00181126 | USA | VA | Powhatan | 36.6818706 | -80.284636 |
| E1512 | *Heuchera americana var. americana* | Wells 3295 | NCU | NCU00181128 | USA | VA | Surry | 37.1118778 | -76.895924 |
| E1513 | *Heuchera americana var. americana* | Fur 439 | NCU | NCU00181129 | USA | VA | Washington | 37.1118778 | -76.895924 |
| E1514 | *Heuchera americana var. americana* | Salle 324 | NCU | NCU00181130 | USA | VA | York | 36.740236 | -81.942167 |
| E1515 | *Heuchera americana var. americana* | Massey 4627 | NCU | NCU00181131 | USA | AL | Jackson | 34.744036 | -86.30427 |
| E1516 | *Heuchera americana var. americana* | Lelong 4564 | NCU | NCU00181132 | USA | AL | Marion | 34.7692447 | -85.986794 |
| E1517 | *Heuchera americana var. americana* | Tucker 4003 | NCU | NCU00128865 | USA | AR | Johnson | 35.624429 | -93.291523 |
| E1518 | *Heuchera americana var. americana* | Merrill 1844 | NCU | NCU00128866 | USA | AR | Pulaski | 35.5600697 | -93.455011 |
| E1519 | *Heuchera americana var. americana* | Younors s.n. | NCU | NCU00181133 | USA | GA | Bulloch | 32.393408 | -81.74381 |
| E1520 | *Heuchera americana var. americana* | Thieret 22647 | NCU | NCU00181134 | USA | LA | Caddo | 32.577195 | -93.882423 |
| E1521 | *Heuchera americana var. americana* | Windler 3822 | NCU | NCU00181135 | USA | MD | Baltimore | 39.417425 | -76.541639 |
| E1522 | *Heuchera americana var. americana* | Hickey 348 | NCU | NCU00181136 | USA | MD | Frederick | 39.2908816 | -76.610759 |
| E1523 | *Heuchera americana var. americana* | Richards 3790 | NCU | NCU00142730 | USA | AR | Craighead | 36.142026 | -90.575279 |
| E1524 | *Heuchera americana var. hirsuticaulis* | Demaree 33248 | NCU | NCU00092970 | USA | AR | Craighead | 35.8348413 | -90.644114 |
| E1525 | *Heuchera americana var. hirsuticaulis* | Demaree 5138 | NCU | NCU00092973 | USA | AR | Franklin | 35.8348413 | -90.644114 |
| E1526 | *Heuchera americana var. hirsuticaulis* | Stephens 10575 | NCU | NCU00092972 | USA | AR | Franklin | 35.5110075 | -93.886485 |
| E1527 | *Heuchera americana var. hirsuticaulis* | Hardin 634 | NCU | NCU00092971 | USA | AR | Franklin | 35.5110075 | -93.886485 |
| E1529 | *Heuchera americana var. hirsuticaulis* | Iltis 5477 | NCU | NCU00092974 | USA | AR | Newton | 35.943319 | -93.070165 |

| | | | | | | | | | |
|---|---|---|---|---|---|---|---|---|---|
| E1531 | *Heuchera americana var. hirsuticaulis* | Hawkins 385 | NCU | NCU00142731 | USA | AR | Randolph | 36.306896 | -91.085753 |
| E1532 | *Heuchera americana var. hirsuticaulis* | Furr 649 | NCU | NCU00092977 | USA | AR | Searcy | 36.3412315 | -91.038702 |
| E1533 | *Heuchera americana var. hirsuticaulis* | Redfearn 31682 | NCU | NCU00128867 | USA | AR | Stone | 35.995665 | -92.297486 |
| E1534 | *Heuchera americana var. hirsuticaulis* | Demaree 49855 | NCU | NCU00092978 | USA | AR | Van Buren | 35.8317159 | -92.174763 |
| E1535 | *Heuchera americana var. hirsuticaulis* | Hughes 76 | NCU | NCU00092981 | USA | AR | Washington | 35.5759086 | -92.491614 |
| E1536 | *Heuchera americana var. hirsuticaulis* | Wells 3301 | NCU | NCU00092986 | USA | KY | Edmonson | 35.9672625 | -94.228066 |
| E1537 | *Heuchera americana var. hirsuticaulis* | Windler 2497 | NCU | NCU00092991 | USA | KY | Livingston | 37.1999181 | -86.220674 |
| E1538 | *Heuchera americana var. hirsuticaulis* | Ellis 01089 | NCU | NCU00092982 | USA | KY | Lyon | 37.1898627 | -88.334383 |
| E1539 | *Heuchera americana var. hirsuticaulis* | Ellis 01338 | NCU | NCU00092987 | USA | KY | Trigg | 36.9981232 | -88.065815 |
| E1540 | *Heuchera americana var. hirsuticaulis* | Ellis 01393 | NCU | NCU00092983 | USA | KY | Trigg | 36.7949221 | -87.878243 |
| E1541 | *Heuchera americana var. heteradenia* | Parrish 40 | NCU | NCU00092996 | USA | GA | Screven | 32.744751 | -81.617585 |
| E1542 | *Heuchera americana var. americana* | Park s.n. | NCU | NCU00092992 | USA | GA | Screven | 32.744751 | -81.617585 |
| E1543 | *Heuchera americana var. hirsuticaulis* | Clebsch s.n. | NCU | NCU00092988 | USA | TN | Montgomery | 36.439722 | -87.296667 |
| E1544 | *Heuchera americana var. americana* | Shaw s.n. | NCU | NCU00439730 | USA | TN | Polk | 35.103928 | -84.55709 |
| E1546 | *Heuchera americana var. hispida* | Sharp 319 | NCU | NCU00045132 | USA | VA | Augusta | | |
| E1547 | *Heuchera americana var. hispida* | Freer 3943 | NCU | NCU00045133 | USA | VA | Augusta | 38.1793809 | -79.152576 |
| E1548 | *Heuchera americana var. hispida* | Freer 1564 | NCU | NCU00045134 | USA | VA | Bedford | 38.1793809 | -79.152576 |
| E1549 | *Heuchera americana var. hispida* | Furr 497 | NCU | NCU00045135 | USA | VA | Botetourt | 37.545183 | -79.971503 |
| E1550 | *Heuchera americana var. hispida* | Furr 484 | NCU | NCU00045136 | USA | VA | Botetourt | 37.544884 | -79.971284 |
| E1551 | *Heuchera americana var. hispida* | Fernald 1596 | NCU | NCU00045137 | USA | VA | Botetourt | 37.482975 | -79.668629 |

| | | | | | | | | | |
|---|---|---|---|---|---|---|---|---|---|
| E1552 | *Heuchera americana var. hispida* | Wells 3313 | NCU | NCU00045138 | USA | VA | Craig | 37.534587 | -80.235484 |
| E1553 | *Heuchera americana var. hispida* | Furr 480 | NCU | NCU00045139 | USA | VA | Craig | 37.542294 | -79.969289 |
| E1554 | *Heuchera americana var. hispida* | Furr 474 | NCU | NCU00045140 | USA | VA | Craig | 37.48337 | -80.130283 |
| E1556 | *Heuchera americana var. hispida* | Hawill 14127 | NCU | NCU00045142 | USA | VA | Craig | 37.357343 | -80.439593 |
| E1557 | *Heuchera americana var. hispida* | Wells 3293 | NCU | NCU00045143 | USA | VA | Fauquier | 38.825515 | -77.719505 |
| E1558 | *Heuchera americana var. hispida* | Massey W3315 | NCU | NCU00045144 | USA | VA | Giles | 37.605918 | -80.240697 |
| E1559 | *Heuchera americana var. hispida* | Cooperrider 4456 | NCU | NCU00045145 | USA | VA | Giles | 37.270794 | -80.694765 |
| E1560 | *Heuchera americana var. hispida* | Furr 464 | NCU | NCU00045146 | USA | VA | Giles | 37.266857 | -80.656989 |
| E1561 | *Heuchera americana var. hispida* | Harvill 14272 | NCU | NCU00045147 | USA | VA | Giles | 37.605918 | -80.240697 |
| E1562 | *Heuchera americana var. hispida* | Harvill 16432 | NCU | NCU00045148 | USA | VA | Grayson | 36.576455 | -81.158355 |
| E1563 | *Heuchera americana var. hispida* | Harvill 19416 | NCU | NCU00045149 | USA | VA | Loudoun | 39.319112 | -77.710924 |
| E1564 | *Heuchera americana var. hispida* | Kral 10351 | NCU | NCU00045150 | USA | VA | Montgomery | 37.244208 | -80.604519 |
| E1565 | *Heuchera americana var. hispida* | Smyth 1055 | NCU | NCU00045151 | USA | VA | Montgomery | 38.1803833 | -81.328445 |
| E1566 | *Heuchera americana var. hispida* | Ahles 62433 | NCU | NCU00045153 | USA | VA | Pittsylvania | 37.067349 | -79.105487 |
| E1567 | *Heuchera americana var. hispida* | Furr 454 | NCU | NCU00045154 | USA | VA | Pulaski | 37.208469 | -80.738294 |
| E1568 | *Heuchera americana var. hispida* | Wood 5819 | NCU | NCU00045155 | USA | VA | Roanoke | 37.140928 | -80.118981 |
| E1569 | *Heuchera americana var. hispida* | Furr 393 | NCU | NCU00045157 | USA | VA | Scott | 37.270973 | -79.941431 |
| E1570 | *Heuchera americana var. hispida* | Uttal 10455 | NCU | NCU00045158 | USA | VA | Tazewell | 36.7117473 | -82.589305 |
| E1571 | *Heuchera americana var. hispida* | Furr 434 | NCU | NCU00045159 | USA | VA | Wise | 36.90057 | -82.310079 |
| E1572 | *Heuchera americana var. hispida* | Furr 449 | NCU | NCU00045160 | USA | VA | Wythe | 36.886036 | -81.192015 |

| | | | | | | | | | |
|---|---|---|---|---|---|---|---|---|---|
| E1573 | *Heuchera americana var. hispida* | Downs 3488 | NCU | NCU00196809 | USA | MD | Washington | 39.346451 | -77.726568 |
| E1574 | *Heuchera americana var. hispida* | Wells 3314 | NCU | NCU00053746 | USA | WV | Greenbrier | 37.806692 | 80.06763 |
| E1575 | *Heuchera americana var. hispida* | Wells 3312 | NCU | NCU00053747 | USA | WV | Monroe | 37.961175 | -80.450934 |
| E1576 | *Heuchera americana var. hispida* | Southern Appalachian Botanical Club 320 | NCU | NCU00053748 | USA | WV | Pocahontas | 37.5585821 | -80.519223 |
| E1578 | *Heuchera americana var. americana* | Bussey 614 | NCU | NCU00181137 | USA | AL | Clay | 37.5969073 | -81.536493 |
| E1579 | *Heuchera americana var. americana* | Orzell 9505 | NCU | NCU00181138 | USA | AL | Cleburne | 33.721111 | -85.604444 |
| E1580 | *Heuchera americana var. hirsuticaulis* | Lipscomb 1564 | NCU | NCU00128864 | USA | AR | Izard | 36.118644 | -92.153699 |
| E1581 | *Heuchera americana var. americana* | Hill 349 | NCU | NCU00181139 | USA | GA | Morgan | 33.661916 | -83.593355 |
| E1596 | *Heuchera pubescens* | Gupton 3549 | NCU | NCU00190348 | USA | VA | Bath | 38.038103 | 79.763563 |
| E1598 | *Heuchera pubescens* | Wells 3316 | NCU | NCU00190350 | USA | VA | Craig | 37.546354 | -79.976945 |
| E1599 | *Heuchera pubescens* | Ramsey 4693 | NCU | NCU00190352 | USA | VA | Franklin | 36.999573 | -79.878086 |
| E1600 | *Heuchera pubescens* | Mitchell 4107 | NCU | NCU00190353 | USA | VA | Franklin | 37.007717 | -79.889332 |
| E1601 | *Heuchera pubescens* | Kral 10306 | NCU | NCU00190354 | USA | VA | Montgomery | 37.215641 | -80.268486 |
| E1602 | *Heuchera pubescens* | Johnson 4565 | NCU | NCU00190355 | USA | VA | Patrick | 36.720388 | -80.327535 |
| E1603 | *Heuchera pubescens* | Ahles 60159 | NCU | NCU00190356 | USA | VA | Patrick | 36.717451 | -80.323062 |
| E1677 | *Heuchera americana var. hispida* | Kral 63868 | NCU | NCU00386386 | USA | VA | Montgomery | 37.191354 | -80.36668 |
| E1679 | *Heuchera americana var. americana* | Cusick 28987 | NCU | NCU00190481 | USA | KY | Greenup | 38.658386 | -83.04535 |
| E1680 | *Heuchera americana var. hirsuticaulis* | Ugent 81-70 | NCU | NCU00196305 | USA | IL | Calhoun | 38.999907 | -90.588731 |
| E1681 | *Heuchera americana var. hirsuticaulis* | Raven 27489 | NCU | NCU00196306 | USA | MO | Jefferson | 38.125 | -90.675 |
| E1682 | *Heuchera americana var. americana* | Wells 3298 | NCU | NCU00196308 | USA | NJ | Hunterdon | 40.475307 | -75.05958 |
| E1683 | *Heuchera americana var. americana* | Cusick 596 | NCU | NCU00196316 | USA | OH | Jefferson | 40.353147 | -80.691019 |
| E1684 | *Heuchera americana var. americana* | Cusick 7568 | NCU | NCU00196317 | USA | OH | Monroe | 39.65929 | -81.06818 |

| | | | | | | | | | |
|---|---|---|---|---|---|---|---|---|---|
| E1685 | *Heuchera americana var. americana* | Cusick 7961 | NCU | NCU00196318 | USA | OH | Noble | 39.7074 | -81.583942 |
| E1686 | *Heuchera americana var. americana* | Cooperrider 6745 | NCU | NCU00196319 | USA | OH | Portage | 41.216555 | -81.301325 |
| E1687 | *Heuchera americana var. americana* | Ziegler 371 | NCU | NCU00196320 | USA | OK | McCurtain | 34.09695 | -94.7043 |
| E1688 | *Heuchera americana var. americana* | Wells 3299 | NCU | NCU00196310 | USA | PA | Bucks | 40.563889 | -75.097729 |
| E1689 | *Heuchera americana var. americana* | Krouse s.n. | NCU | NCU00196312 | USA | PA | Fayette | 39.871742 | -79.492261 |
| E1690 | *Heuchera americana var. americana* | Wells 3297 | NCU | NCU00196313 | USA | PA | Lancaster | 39.940464 | -75.993633 |
| E1691 | *Heuchera americana var. americana* | Utech 82-188 | NCU | NCU00196315 | USA | PA | Westmoreland | 40.129167 | -79.291667 |
| E1692 | *Heuchera americana var. americana* | Correll 37142 | NCU | NCU00196493 | USA | TX | Bowie | 33.516333 | -94.147 |
| E1693 | *Heuchera americana var. hirsuticaulis* | Furr 592 | NCU | NCU00196495 | USA | IL | Alexander | 37.208057 | -89.428705 |
| E1694 | *Heuchera americana var. hirsuticaulis* | Winterringer 901 | NCU | NCU00196496 | USA | IL | Hardin | 37.4999684 | -88.237835 |
| E1697 | *Heuchera americana var. hirsuticaulis* | Bollwinkel FC 124 | NCU | NCU00196500 | USA | IL | Johnson | 37.4501818 | -88.884405 |
| E1698 | *Heuchera americana var. hirsuticaulis* | Neill 15228 | NCU | NCU00196550 | USA | IL | Saint Clair | 38.4616972 | -89.932435 |
| E1699 | *Heuchera americana var. hirsuticaulis* | Bartlett 2109 | NCU | NCU00196551 | USA | IN | Brown | 39.1682855 | -86.2297 |
| E1702 | *Heuchera americana var. hirsuticaulis* | Wells 3306 | NCU | NCU00196555 | USA | IN | Howard | 40.4787061 | -86.135034 |
| E1703 | *Heuchera americana var. hirsuticaulis* | Wells 3303 | NCU | NCU00196556 | USA | IN | Montgomery | 40.0361447 | -86.900708 |
| E1704 | *Heuchera americana var. hirsuticaulis* | Furr 594 | NCU | NCU00196559 | USA | MO | Boone | 39.0153926 | -92.330753 |
| E1705 | *Heuchera americana var. hirsuticaulis* | Furr 668 | NCU | NCU00196801 | USA | MO | Shannon | 37.1498052 | -91.432825 |
| E1706 | *Heuchera americana var. hirsuticaulis* | Redfearn 27414 | NCU | NCU00196802 | USA | MO | Shannon | 37.093791 | -91.209527 |
| E1707 | *Heuchera americana var. hirsuticaulis* | Dorr 387 | NCU | NCU00196803 | USA | MO | Saint Louis | 38.6319657 | -90.242876 |
| E1708 | *Heuchera americana var. hirsuticaulis* | Furr 665 | NCU | NCU00196804 | USA | MO | Taney | 36.6563729 | -93.066578 |

| | | | | | | | | | |
|---|---|---|---|---|---|---|---|---|---|
| E1709 | *Heuchera americana var. americana* | O'Dell 276 | NCU | NCU00196806 | USA | OH | Vinton | 39.2744355 | -82.47403 |
| E1711 | *Heuchera americana var. hirsuticaulis* | Wallis 6790-1 | NCU | NCU00196808 | USA | OK | Sequoyah | 35.5035863 | -94.736058 |
| E1780 | *Heuchera richardsonii* | Huang 3045 | NCU | NCU00196884 | USA | IA | Cedar | 41.7608608 | -91.12632 |
| E1781 | *Heuchera richardsonii* | Cooperrider 1120 | NCU | NCU00196885 | USA | IA | Jackson | 42.1420382 | -90.548007 |
| E1782 | *Heuchera richardsonii* | Walker 231 | NCU | NCU00196894 | USA | IA | Jackson | 42.1420382 | -90.548007 |
| E1783 | *Heuchera richardsonii* | Witlake 905 | NCU | NCU00196887 | USA | IA | Lyon | 43.3747071 | -96.208192 |
| E1784 | *Heuchera richardsonii* | DeBurh 1184 | NCU | NCU00196888 | USA | IA | Lyon | 43.3747071 | -96.208192 |
| E1785 | *Heuchera richardsonii* | Myron 13 | NCU | NCU00196889 | USA | IA | Pocahontas | 42.7262681 | -94.647738 |
| E1786 | *Heuchera richardsonii* | Schwab 209 | NCU | NCU00196891 | USA | IA | Story | 42.040106 | -93.634508 |
| E1787 | *Heuchera richardsonii* | Freckmann 1882 | NCU | NCU00196892 | USA | IA | Story | 42.0161612 | -93.489194 |
| E1788 | *Heuchera richardsonii* | Wagenknecht 287 | NCU | NCU00196893 | USA | IA | Washington | 41.3160082 | -91.734033 |
| E1789 | *Heuchera richardsonii* | Utech 1925 | NCU | NCU00196912 | USA | WI | Crawford | 43.19824 | -90.874083 |
| E1790 | *Heuchera richardsonii* | Rice 1902 | NCU | NCU00196913 | USA | WI | Rock | 42.606119 | -89.02125 |
| E1791 | *Heuchera richardsonii* | Rice W3351 | NCU | NCU00196915 | USA | WI | Rock | 42.606119 | -89.02125 |
| E1792 | *Heuchera richardsonii* | Rice W3349 | NCU | NCU00196917 | USA | WI | Rock | 42.824114 | -89.318987 |
| E1793 | *Heuchera richardsonii* | Rice W3350 | NCU | NCU00196918 | USA | WI | Rock | 42.59156 | -89.021029 |
| E1794 | *Heuchera richardsonii* | Jackson 628 | NCU | NCU00196898 | USA | CO | El Paso | 38.827383 | -104.52747 |
| E1795 | *Heuchera richardsonii* | Wells 3347 | NCU | NCU00196899 | USA | IN | Fulton | 41.0421416 | -86.287521 |
| E1796 | *Heuchera richardsonii* | Wells 3346 | NCU | NCU00196900 | USA | IN | Porter | 41.4457032 | -87.072497 |
| E1797 | *Heuchera richardsonii* | McGregor 15586 | NCU | NCU00196901 | USA | KS | Cherokee | 37.1718068 | -94.848207 |
| E1798 | *Heuchera richardsonii* | Kukla 91 | NCU | NCU00196902 | USA | MN | Clay | 46.8994904 | -96.50882 |
| E1799 | *Heuchera richardsonii* | Steyermark 84538 | NCU | NCU00196904 | USA | MO | Andrew | 39.905814 | -94.730975 |
| E1800 | *Heuchera richardsonii* | Palmer 51751 | NCU | NCU00196905 | USA | MO | Dade | 37.4323036 | -93.84063 |
| E1801 | *Heuchera richardsonii* | Redfearn 14465 | NCU | NCU00196906 | USA | MO | Sainte Geneveive | 37.781647 | -90.284423 |
| E1802 | *Heuchera richardsonii* | Stephens 23426 | NCU | NCU00196908 | USA | ND | Golden Valley | 46.9317944 | -96.947075 |
| E1804 | *Heuchera richardsonii* | Wallis 6866 | NCU | NCU00196910 | USA | OK | Ottawa | 36.835764 | -94.802681 |
| E1805 | *Heuchera richardsonii* | Uttal 9346 | NCU | NCU00196911 | USA | SD | Custer | 43.684943 | -103.46225 |

| | | | | | | | | | |
|---|---|---|---|---|---|---|---|---|---|
| E1852 | *Heuchera americana var. americana* | Radford 6853 | NCU | NCU00181741 | USA | NC | Bladen | 34.627971 | -78.562771 |
| E1853 | *Heuchera americana var. americana* | McCormick s.n. | NCU | NCU00075922 | USA | NC | Buncombe | 35.64484 | -82.282619 |
| E1854 | *Heuchera americana var. americana* | Bradford 0011 | NCU | NCU00181746 | USA | NC | Burke | 35.834001 | -81.711678 |
| E1855 | *Heuchera americana var. americana* | Bell 11904 | NCU | NCU00180302 | USA | NC | Caswell | 36.343309 | -79.438356 |
| E1856 | *Heuchera americana var. americana* | Crutchfield 1304 | NCU | NCU00180304 | USA | NC | Chatham | 35.757001 | -79.088078 |
| E1857 | *Heuchera americana var. americana* | Ahles 57961 | NCU | NCU00180308 | USA | NC | Durham | 36.073411 | -78.873341 |
| E1858 | *Heuchera americana var. americana* | Radford 13285 | NCU | NCU00180310 | USA | NC | Graham | 35.44307 | -83.937459 |
| E1859 | *Heuchera americana var. americana* | Radford 13212 | NCU | NCU00180311 | USA | NC | Graham | 36.0690258 | -79.400576 |
| E1860 | *Heuchera americana var. americana* | Radford 43889 | NCU | NCU00180312 | USA | NC | Granville | 36.084312 | -78.746671 |
| E1862 | *Heuchera americana var. americana* | Downs 13390 | NCU | NCU00014279 | USA | NC | Harnett | 35.468214 | -78.898074 |
| E1863 | *Heuchera americana var. americana* | Wells 3284 | NCU | NCU00181750 | USA | NC | Haywood | 35.645284 | -82.941249 |
| E1864 | *Heuchera americana var. americana* | Ramseur 3329 | NCU | NCU00181749 | USA | NC | Haywood | 35.409551 | -82.856243 |
| E1866 | *Heuchera americana var. americana* | Radford 5793 | NCU | NCU00181757 | USA | NC | Hertford | 36.468816 | -77.095399 |
| E1868 | *Heuchera americana var. americana* | Stewart 405 | NCU | NCU00180318 | USA | NC | Lee | 35.4691746 | -79.154764 |
| E1869 | *Heuchera americana var. americana* | Kessler 222 | NCU | NCU00180319 | USA | NC | Lee | 35.5386 | -79.246693 |
| E1870 | *Heuchera americana var. americana* | Wells 3282 | NCU | NCU00180320 | USA | NC | Lee | 35.4691746 | -79.154764 |
| E1871 | *Heuchera americana var. americana* | Kessler 242 | NCU | NCU00180321 | USA | NC | Lee | 35.536819 | -79.252642 |
| E1872 | *Heuchera americana var. americana* | Wells 3283 | NCU | NCU00180322 | USA | NC | Macon | 35.1436639 | -83.39773 |
| E1873 | *Heuchera americana var. americana* | Radford s.n. | NCU | NCU00180323 | USA | NC | Macon | 35.148812 | -83.300076 |
| E1874 | *Heuchera americana var. americana* | Wells 3287 | NCU | NCU00180326 | USA | NC | Madison | 35.8482034 | -82.693106 |

| | | | | | | | | | |
|---|---|---|---|---|---|---|---|---|---|
| E1875 | *Heuchera americana var. americana* | Wells 3285 | NCU | NCU00180328 | USA | NC | Madison | 35.737329 | -82.869151 |
| E1876 | *Heuchera americana var. americana* | Treiber 425 | NCU | NCU00180331 | USA | NC | Martin | 35.966359 | -77.209684 |
| E1877 | *Heuchera americana var. americana* | Leonard 4793 | NCU | NCU00180332 | USA | NC | McDowell | 35.6608869 | -82.048217 |
| E1879 | *Heuchera americana var. americana* | Wells 3288 | NCU | NCU00180334 | USA | NC | Mitchell | 36.0001181 | -82.134903 |
| E1880 | *Heuchera americana var. americana* | Ahles 43133 | NCU | NCU00180335 | USA | NC | Mitchell | 36.0001181 | -82.134903 |
| E1881 | *Heuchera americana var. americana* | Radford 42974 | NCU | NCU00180339 | USA | NC | Montgomery | 35.3299572 | -79.897902 |
| E1882 | *Heuchera americana var. americana* | Kessler 272 | NCU | NCU00078066 | USA | NC | Moore | 35.3054614 | -79.476124 |
| E1883 | *Heuchera americana var. americana* | Ahles 11797 | NCU | NCU00180340 | USA | NC | Nash | 35.992983 | -77.975531 |
| E1884 | *Heuchera americana var. americana* | Ahles 41790 | NCU | NCU00180341 | USA | NC | Northampton | 36.4168078 | -77.364223 |
| E1885 | *Heuchera americana var. americana* | Munch s.n. | NCU | NCU00061747 | USA | NC | Orange | 35.914781 | -79.039 |
| E1887 | *Heuchera americana var. americana* | Wells 3281 | NCU | NCU00180343 | USA | NC | Orange | 36.0605095 | -79.117268 |
| E1888 | *Heuchera americana var. americana* | Radford 7569 | NCU | NCU00180344 | USA | NC | Orange | 36.0605095 | -79.117268 |
| E1889 | *Heuchera americana var. americana* | Larke 1193 | NCU | NCU00180345 | USA | NC | Orange | 35.891961 | -79.037875 |
| E1893 | *Heuchera americana var. americana* | White s.n. | NCU | NCU00180353 | USA | NC | Orange | 36.0605095 | -79.117268 |
| E1894 | *Heuchera americana var. americana* | Wickland 904 | NCU | NCU00180357 | USA | NC | Randolph | 35.738333 | -80.020833 |
| E1895 | *Heuchera americana var. americana* | Sorrie 9714 | NCU | NCU00180358 | USA | NC | Richmond | 35.0288383 | -79.733326 |
| E1896 | *Heuchera americana var. americana* | Radford 11479 | NCU | NCU00180359 | USA | NC | Richmond | 35.0288383 | -79.733326 |
| E1899 | *Heuchera americana var. americana* | Boufford 13648 | NCU | NCU00180482 | USA | NC | Swain | 35.45819 | -83.466275 |
| E1900 | *Heuchera americana var. americana* | Ahles 56581 | NCU | NCU00180483 | USA | NC | Wake | 35.7979355 | -78.611831 |
| E1901 | *Heuchera americana var. americana* | Bell 2911 | NCU | NCU00180484 | USA | NC | Warren | 36.3901694 | -78.105212 |

| | | | | | | | | | |
|---|---|---|---|---|---|---|---|---|---|
| E1902 | *Heuchera americana var. americana* | Downs 13696 | NCU | NCU00169493 | USA | NC | Wilkes | 36.17371 | -81.169751 |
| E1903 | *Heuchera americana var. americana* | Stewart s.n. | NCU | NCU00180488 | USA | NC | Wilkes | 36.1998247 | -81.134135 |
| E1904 | *Heuchera americana var. americana* | Ahles 42660 | NCU | NCU00180490 | USA | NC | Yancey | 35.913356 | -82.455335 |
| E1905 | *Heuchera caroliniana* | Radford 1351 | NCU | NCU00053715 | USA | NC | Alexander | 35.972354 | -81.108692 |
| E1911 | *Heuchera caroliniana* | Ahles 41083 | NCU | NCU00053722 | USA | NC | Iredell | 35.604503 | -80.905932 |
| E1915 | *Heuchera caroliniana* | Williams s.n. | NCU | NCU00053725 | USA | NC | Mecklenburg | 35.2356385 | -80.813949 |
| E1916 | *Heuchera americana var. americana* | Sorrie 13301 | NCU | NCU00439687 | USA | NC | Moore | 35.3054614 | -79.476124 |
| E1921 | *Heuchera caroliniana* | Radford 11873 | NCU | NCU00053736 | USA | NC | Stanly | 35.3235477 | -80.239137 |
| E1924 | *Heuchera caroliniana* | Radford 12001 | NCU | NCU00053739 | USA | NC | Union | 34.9795158 | -80.512821 |
| E1926 | *Heuchera americana var. hispida* | Poindexter 08-460 | NCU | NCU00128800 | USA | NC | Alleghany | 36.5691944 | 81.1784167 |
| E1927 | *Heuchera americana var. hispida* | Poindexter 08-350 | NCU | NCU00128901 | USA | NC | Alleghany | 36.5703611 | 81.4518889 |
| E1928 | *Heuchera americana var. hispida* | Radford 13141 | NCU | NCU00180492 | USA | NC | Surry | 36.551292 | -80.909589 |
| E1936 | *Heuchera pubescens* | McCurdy 449 | NCU | NCU00064407 | USA | NC | Stokes | 36.4120995 | -80.228809 |
| E306-2 | *Heuchera alba* | USDA GRIN: Ames 34945 | Sent directly | | USA | WV | Grant | 39.0034 | -79.2204 |
| E35 | *Heuchera caroliniana* | Haesloop 504 | UNA | UNA00014676 | USA | NC | Stanly | 35.3235477 | -80.239137 |
| E36 | *Heuchera americana var. americana* | Horn 1644 | UNA | UNA00014678 | USA | SC | Newberry | 34.3266879 | -81.583009 |
| E438 | *Heuchera americana var. americana* | Browne 70K14.5 | MEM | | USA | KY | Pulaski | 37.1124781 | -84.593893 |
| E439 | *Heuchera americana var. americana* | Gentry 926 | MEM | | USA | KY | Henry | 38.447983 | -85.119117 |
| E440 | *Heuchera americana var. americana* | Bates 2064 | MEM | | USA | TN | Decatur | 35.6067875 | -88.108398 |
| E441 | *Heuchera americana var. hirsuticaulis* | Bates 1758 | MEM | | USA | TN | Hardeman | 35.1743058 | -88.996336 |
| E442 | *Heuchera americana var. hirsuticaulis* | Bates 2205 | MEM | | USA | TN | McNairy | 35.1663375 | -88.576617 |
| E443 | *Heuchera americana var. americana* | Athey 3076 | MEM | | USA | KY | Bell | 36.7370344 | -83.64917 |

| | | | | | | | | | |
|---|---|---|---|---|---|---|---|---|---|
| E490 | *Heuchera americana var. americana* | Holmes 10831 | TEX | 211832 | USA | TX | Bowie | 33.4198886 | -94.447963 |
| E491 | *Heuchera americana var. americana* | Correll 37142 | TEX | 459919 | USA | TX | Bowie | 33.4198886 | -94.447963 |
| E492 | *Heuchera americana var. americana* | Correll 31281 | TEX | 352864 | USA | TX | Bowie | 33.4198886 | -94.447963 |
| E498 | *Heuchera americana var. hirsuticaulis* | D'Arcy 4497 | TEX | | USA | AR | Yell | 35.221944 | -93.243611 |
| E499 | *Heuchera americana var. hirsuticaulis* | Kral 67083 | TEX | | USA | AR | Garland | 34.5488944 | -93.183854 |
| E500 | *Heuchera americana var. hirsuticaulis* | Krall 61777 | TEX | | USA | AR | Howard | 34.0744088 | -93.974578 |
| E501 | *Heuchera americana var. hirsuticaulis* | Kral 59876 | TEX | | USA | AR | Faulkner | 35.1470851 | -92.321905 |
| E502 | *Heuchera americana var. americana* | Orzell 9505 | TEX | | USA | AL | Cleburne | 33.721111 | -85.604444 |
| E503 | *Heuchera americana var. hirsuticaulis* | Raven 27489 | TEX | | USA | MO | Jefferson | 38.125 | -90.675 |
| E505 | *Heuchera americana var. americana* | Pyron 2492 | TEX | | USA | GA | Burke | 33.0482247 | -81.957598 |
| E506 | *Heuchera americana var. americana* | Moldenke 30033 | TEX | | USA | VA | Amelia | 37.3319664 | -78.008448 |
| E507 | *Heuchera americana var. americana* | Woodbury s.n. | TEX | | USA | NC | Rutherford | 35.4833 | -81.95 |
| E509 | *Heuchera americana var. americana* | Shoals 30506 | TEX | | USA | AR | Cleburne | 35.5298937 | -92.031282 |
| E510 | *Heuchera americana var. americana* | Brown 2135 | TEX | | USA | NC | Durham | 35.996653 | -78.901805 |
| E511 | *Heuchera americana var. americana* | Crawford 223 | TEX | | USA | AL | Fayette | 33.7303493 | -87.741909 |
| E513 | *Heuchera americana var. hirsuticaulis* | Randrianaivo 423 | TEX | | USA | MO | Saint Genevieve | 37.828889 | -90.225556 |
| E514 | *Heuchera americana var. americana* | Kral 63437 | TEX | | USA | GA | Floyd | 34.2421984 | -85.222925 |
| E515 | *Heuchera americana var. americana* | Wofford 96-38 | TEX | | USA | TN | Grainger | 36.2810821 | -83.510702 |
| E516 | *Heuchera americana var. americana* | Haesloop 504 | TEX | | USA | NC | Stanly | 35.3235477 | -80.239137 |
| E517 | *Heuchera americana var. americana* | Roller 1/60 | TEX | | USA | KY | Madison | 37.7298081 | -84.297474 |

| | | | | | | | | | |
|---|---|---|---|---|---|---|---|---|---|
| E518 | *Heuchera americana var. americana* | Moldenke 27133 | TEX | | USA | TN | Blount | 35.6719722 | -83.931447 |
| E519 | *Heuchera americana var. americana* | Moldenke 27000 | TEX | | USA | SC | McCormick | 33.911441 | -82.295433 |
| E521 | *Heuchera americana var. hirsuticaulis* | Hardin 634 | TEX | | USA | AR | Franklin | 35.5110075 | -93.886485 |
| E522 | *Heuchera americana var. americana* | McVaugh 8610 | TEX | | USA | AL | Randolph | 33.2651763 | -85.469396 |
| E523 | *Heuchera americana var. americana* | Bright 15210 | TEX | | USA | PA | Allegheny | 40.4597204 | -79.976041 |
| E524 | *Heuchera americana var. americana* | McNeilus 97-384 | TEX | | USA | TN | Campbell | 36.3865389 | -84.135162 |
| E542 | *Heuchera richardsonii* | Chase 9546 | TEX | | USA | IL | McHenry | 42.3294391 | -88.460571 |
| E543 | *Heuchera richardsonii* | Welch 9839 | TEX | | USA | IA | Dickinson | 43.3753018 | -95.165428 |
| E639 | *Heuchera richardsonii* | Breitung 1139 | TEX | | Canada | Saskatchewan | Wallwort | 52.57045 | -104.01686 |
| E640 | *Heuchera richardsonii* | Chase 14460 | TEX | | USA | IL | Peoria | 41.4194058 | -89.589101 |
| E641 | *Heuchera richardsonii* | Chase 11914 | TEX | | USA | IL | Will | 45.9009106 | -87.999475 |
| FL-25 | *Heuchera longiflora var. longiflora* | Floden 3208 | OS | | USA | VA | Lee | 36.6679833 | -83.230925 |
| FL-26 | *Heuchera longiflora var. longiflora* | Floden s.n. | OS | | USA | AL | Talladega | 33.1415972 | -86.258378 |
| H105-1 | *Heuchera richardsonii* | Folk 105 | OS | | USA | SD | Pennington | 43.8438361 | -102.43763 |
| H106-1 | *Heuchera richardsonii* | Folk 106 | OS | | USA | SD | Pennington | 44.4666667 | -102.0525 |
| H106-2 | *Heuchera richardsonii* | Folk 106 | OS | | USA | SD | Pennington | 44.4666667 | -102.0525 |
| H166-2 | *Heuchera richardsonii* | Folk 166 | OS | | USA | IA | Boone | 41.9966667 | -93.883611 |
| H167-1 | *Heuchera richardsonii* | Folk 167 | OS | | USA | IL | Porter | 41.3213889 | -87.006389 |
| H167-2 | *Heuchera richardsonii* | Folk 167 | OS | | USA | IL | Porter | 41.3213889 | -87.006389 |
| H177-2 | *Heuchera americana var. hirsuticaulis* | Folk 177 | OS | | USA | IL | Union | 37.5415333 | -89.426917 |
| H246 | *Heuchera americana var. americana* | Folk 246 | OS | | USA | OH | Delaware | 40.267491 | -82.951414 |
| H59-2 | *Heuchera americana var. hirsuticaulis* | Folk 59 | OS | | USA | MO | Shannon | 37.39023 | -91.45272 |
| H59-2 | *Heuchera americana var. hirsuticaulis* | Folk 59 | OS | | USA | MO | Shannon | 37.39023 | -91.45272 |

| | | | | | | | | | |
|---|---|---|---|---|---|---|---|---|---|
| H62-1 | *Heuchera americana var. hirsuticaulis* | Folk 62 | OS | | USA | MO | Scott | 37.19118 | 89.63458 |
| H62-2 | *Heuchera americana var. hirsuticaulis* | Folk 62 | OS | | USA | MO | Scott | 37.19118 | 89.63458 |
| H62-2 | *Heuchera americana var. hirsuticaulis* | Folk 62 | OS | | USA | MO | Scott | 37.19118 | -89.63458 |
| H62-4 | *Heuchera americana var. hirsuticaulis* | Folk 62 | OS | | USA | MO | Scott | 37.19118 | 89.63458 |
| H62-4 | *Heuchera americana var. hirsuticaulis* | Folk 62 | OS | | USA | MO | Scott | 37.19118 | -89.63458 |
| H62-5 | *Heuchera americana var. hirsuticaulis* | Folk 62 | OS | | USA | MO | Scott | 37.19118 | 89.63458 |
| H62-5 | *Heuchera americana var. hirsuticaulis* | Folk 62 | OS | | USA | MO | Scott | 37.19118 | -89.63458 |
| H62-6 | *Heuchera americana var. hirsuticaulis* | Folk 62 | OS | | USA | MO | Scott | 37.19118 | 89.63458 |
| H62-6 | *Heuchera americana var. hirsuticaulis* | Folk 62 | OS | | USA | MO | Scott | 37.19118 | -89.63458 |
| H71 | *Heuchera americana var. americana* | Folk 71 | OS | | USA | OH | Lawrence | 38.576956 | -82.335822 |
| H95 | *Heuchera americana var. americana* | Folk 95 | OS | | USA | NC | Buncombe | 35.625553 | -82.518968 |
| I173 | *Heuchera alba* | Vanderhorst 7485 | MISSA | | USA | WV | Grant | 38.927459 | -79.237163 |
| I174 | *Heuchera alba* | Byers 1570 | MISSA | | USA | WV | Grant | 39.000504 | -79.081646 |
| I175 | *Heuchera alba* | Streets 3790 | MISSA | | USA | WV | Pocahontas | 38.202504 | -80.194893 |
| I176 | *Heuchera alba* | Streets 3004 | MISSA | | USA | WV | Pendleton | 38.505711 | -79.209617 |
| I177 | *Heuchera alba* | Streets 5990 | MISSA | | USA | WV | Pocahontas | 38.365892 | -79.93746 |
| I178 | *Heuchera alba* | Streets 4090 | MISSA | | USA | WV | Grant | 38.984892 | -79.218757 |
| I179 | *Heuchera alba* | Streets 4768 | MISSA | | USA | WV | Pendleton | 38.636815 | -79.394064 |
| I180 | *Heuchera alba* | Streets 4778 | MISSA | | USA | WV | Pendleton | 38.666659 | -79.235843 |
| I2 | *Heuchera americana var. americana* | Folk I2 | OS | | USA | OH | Defiance | 41.174476 | -84.256819 |
| I20 | *Heuchera americana var. americana* | Folk I20 | OS | | USA | VA | Albemarle | 38.094964 | -78.489373 |
| I65 | *Heuchera richardsonii* | Folk I65 | OS | | USA | MI | Berrien | 41.868479 | -86.348772 |
| I67 | *Heuchera alba* | Folk I67 | OS | | USA | WV | Highland | 38.594238 | -79.18181 |
| I82 | *Heuchera richardsonii* | Folk I82 | OS | | Canada | Alberta | | 49.949883 | -113.97907 |

| | | | | | | | | | |
|---|---|---|---|---|---|---|---|---|---|
| I90-C | *Heuchera inconstans* | Folk I90 | OS | | USA | AZ | Coconino | 34.99126 | -111.73521 |
| L1-1 | *Heuchera longiflora var. aceroides* | Folk L-1 | OS | | USA | NC | Madison | 35.7467139 | -82.872478 |
| L1-3 | *Heuchera longiflora var. aceroides* | Folk L-1 | OS | | USA | NC | Madison | 35.7467139 | -82.872478 |
| L14-1 | *Heuchera longiflora var. longiflora* | Folk L-14 | OS | | USA | KY | Floyd | 37.6698389 | -82.913272 |
| L14-2 | *Heuchera longiflora var. longiflora* | Folk L-14 | OS | | USA | KY | Floyd | 37.6698389 | -82.913272 |
| L20-1 | *Heuchera longiflora var. longiflora* | Folk L-20 | OS | | USA | KY | Lee | 37.6304667 | -83.770767 |
| L20-2 | *Heuchera longiflora var. longiflora* | Folk L-20 | OS | | USA | KY | Lee | 37.6304667 | -83.770767 |
| L3-1 | *Heuchera longiflora var. aceroides* | Folk L-3 | OS | | USA | NC | Madison | 35.92865 | -82.750078 |
| L3-2 | *Heuchera longiflora var. aceroides* | Folk L-3 | OS | | USA | NC | Madison | 35.92865 | -82.750078 |
| L6-1 | *Heuchera longiflora var. longiflora* | Folk L-6 | OS | | USA | VA | Wise | 36.8558167 | -82.747758 |
| L6-2 | *Heuchera longiflora var. longiflora* | Folk L-6 | OS | | USA | VA | Wise | 36.8558167 | -82.747758 |
| L9-2 | *Heuchera longiflora var. longiflora* | Folk L-9 | OS | | USA | VA | Wise | 37.1037417 | -82.668386 |

**Table S2.** GC-MS temperature programming.

| **Table S2**. Temperature programming | | | | |
|---|---|---|---|---|
| | Rate °C/min | Value °C | Hold time min. | Runtime min. |
| (Initial) | | 35 | 3 | 3 |
| Ramp 1 | 15 | 150 | 5 | 15.667 |
| Ramp 2 | 50 | 250 | 5 | 22.667 |

| | | | | |
|---|---|---|---|---|
| Ramp 3 | 30 | 280 | 5 | 28.667 |

**Table S3**. Captured pollinator identifications.

| Barcode | Genus | Species | Male | Female | Day | Month | Year | Country | Host population | Pollinator accession |
|---|---|---|---|---|---|---|---|---|---|---|
| VECO020701 | *Colletes* | *aestivalis* | 0 | 1 | 3 | May | 2022 | USA | A1 | AN1 |
| VECO020702 | *Augochlorella* | *striata* | 0 | 1 | 3 | May | 2022 | USA | A1 | AU1 |
| VECO020703 | *Augochlorella* | *striata* | 0 | 1 | 3 | May | 2022 | USA | A1 | AU2 |
| VECO020704 | *Colletes* | *distincta* | 0 | 1 | 5 | Jun | 2022 | USA | A16 | AN1 |
| VECO020705 | *Colletes* | *inaequalis* | 1 | 0 | 5 | Jun | 2022 | USA | A16 | AN2 |
| VECO020706 | *Colletes* | *distincta* | 0 | 1 | 5 | Jun | 2022 | USA | A16 | AN3 |
| VECO020707 | *Colletes* | *distincta* | 0 | 1 | 5 | Jun | 2022 | USA | A16 | AN4 |
| VECO020708 | *Lasioglossum (Dialictus)* | *imitatum* | 0 | 1 | 5 | Jun | 2022 | USA | A16 | LA1 |
| VECO020709 | *Lasioglossum (Dialictus)* | *imitatum* | 0 | 1 | 5 | Jun | 2022 | USA | A16 | LA2 |
| VECO020710 | *Lasioglossum (Dialictus)* | *imitatum* | 0 | 1 | 5 | Jun | 2022 | USA | A16 | LA3 |
| VECO020711 | *Colletes* | *inaequalis?* | 1 | 0 | 11 | Oct | 2022 | USA | A17 | AN1 |
| VECO020712 | *Augochlorella* | *striata* | 0 | 1 | 7 | Jun | 2022 | USA | A17 | AU1 |
| VECO020713 | *Lasioglossum (Dialictus)* | *imitatum* | 0 | 1 | 7 | Jun | 2022 | USA | A17 | LA1 |

| | | | | | | | | | | |
|---|---|---|---|---|---|---|---|---|---|---|
| VECO020714 | *Lasioglossum (Dialictus)* | *imitatum* | 0 | 1 | 7 | Jun | 2022 | USA | A17 | LA2 |
| VECO020715 | *Lasioglossum (Dialictus)* | *imitatum* | 0 | 1 | 7 | Jun | 2022 | USA | A17 | LA3 |
| VECO020716 | *Lasioglossum (Dialictus)* | *creberrimum* | 0 | 1 | 7 | Jun | 2022 | USA | A17 | LA4 |
| VECO020717 | *Lasioglossum (Dialictus)* | *imitatum* | 0 | 1 | 7 | Jun | 2022 | USA | A17 | LA5 |
| VECO020718 | *Lasioglossum (Dialictus)* | *imitatum* | 0 | 1 | 7 | Jun | 2022 | USA | A17 | LA6 |
| VECO020719 | *Augochlorella* | *striata* | 0 | 1 | 2 | May | 2022 | USA | A3 | AU1 |
| VECO020720 | *Augochlorella* | *striata* | 0 | 1 | 2 | May | 2022 | USA | A3 | AU2 |
| VECO020721 | *Augochlorella* | *striata* | 1 | 0 | 13 | Aug | 2021 | USA | A30 | SM1A |
| VECO020722 | *Augochlorella* | *striata* | 0 | 1 | 17 | May | 2021 | USA | A34 | AU1A |
| VECO020723 | *Lasioglossum (Dialictus)* | *imitatum* | 0 | 1 | 17 | May | 2021 | USA | A34 | LA1A |
| VECO020724 | *Lasioglossum (Dialictus)* | *imitatum* | 0 | 1 | 17 | May | 2021 | USA | A34 | LA2 |
| VECO020725 | *Lasioglossum (Dialictus)* | *imitatum* | 0 | 1 | 17 | May | 2021 | USA | A35 | LA1A |
| VECO020726 | *Lasioglossum (Dialictus)* | *imitatum* | 0 | 1 | 16 | May | 2021 | USA | A47 | LA1A |
| VECO020727 | *Colletes* | *aestivalis* | 0 | 1 | 15 | May | 2022 | USA | A55 | AN1 |
| VECO020728 | *Colletes* | *aestivalis* | 0 | 1 | 15 | May | 2022 | USA | A55 | AN2 |
| VECO020729 | *Lasioglossum* | *fuscipenne* | 0 | 1 | 15 | May | 2022 | USA | A55 | AN3 |

| | | | | | | | | | | |
|---|---|---|---|---|---|---|---|---|---|---|
| VECO020730 | *Lasioglossum (Dialictus)* | *imitatum* | 0 | 1 | 15 | May | 2022 | USA | A55 | LA1 |
| VECO020731 | *Lasioglossum (Dialictus)* | *imitatum* | 0 | 1 | 15 | May | 2022 | USA | A55 | LA2 |
| VECO020732 | *Colletes* | *distincta* | 0 | 1 | 11 | May | 2022 | USA | A58 | AN1 |
| VECO020733 | *Colletes* | *inaequalis* | 1 | 0 | 11 | May | 2022 | USA | A58 | AN2 |
| VECO020734 | *Colletes* | *distincta* | 0 | 1 | 11 | May | 2022 | USA | A58 | AN3 |
| VECO020735 | *Lasioglossum (Dialictus)* | *imitatum* | 0 | 1 | 11 | May | 2022 | USA | A58 | LA1 |
| VECO020736 | *Lasioglossum (Dialictus)* | *imitatum* | 0 | 1 | 11 | May | 2022 | USA | A58 | LA2 |
| VECO020737 | *Colletes* | *aestivalis* | 0 | 1 | 13 | May | 2022 | USA | A60 | AN1 |
| VECO020738 | *Colletes* | *aestivalis* | 0 | 1 | 13 | May | 2022 | USA | A60 | AN2 |
| VECO020739 | *Colletes* | *aestivalis* | 0 | 1 | 13 | May | 2022 | USA | A60 | AN3 |
| VECO020740 | *Lasioglossum (Dialictus)* | *imitatum* | 0 | 1 | 14 | May | 2022 | USA | A62 | LA1 |
| VECO020741 | *Lasioglossum (Dialictus)* | *imitatum* | 0 | 1 | 14 | May | 2022 | USA | A62 | LA2 |
| VECO020742 | *Lasioglossum (Dialictus)* | *imitatum* | 0 | 1 | 12 | May | 2022 | USA | CG | LA1 |
| VECO020743 | *Lasioglossum (Dialictus)* | *imitatum* | 0 | 1 | 8 | Jun | 2022 | USA | H166 | LA1 |
| VECO020744 | *Lasioglossum (Dialictus)* | *imitatum* | 0 | 1 | 8 | Jun | 2022 | USA | H166 | LA2 |
| VECO020745 | *Lasioglossum (Dialictus)* | *imitatum* | 0 | 1 | 4 | Jun | 2022 | USA | H167 | LA1 |

| | | | | | | | | | | |
|---|---|---|---|---|---|---|---|---|---|---|
| VECO020746 | *Lasioglossum (Dialictus)* | *imitatum* | 0 | 1 | 4 | Jun | 2022 | USA | H167 | LA2 |
| VECO020747 | *Lasioglossum (Dialictus)* | *imitatum* | 0 | 1 | 4 | Jun | 2022 | USA | H167 | LA3 |
| VECO020748 | *Augochlora* | *pura* | 0 | 1 | 11 | Aug | 2021 | USA | A30 | AU1 |
| VECO020749 | *Augochlora* | *pura* | 0 | 1 | 11 | Aug | 2021 | USA | A30 | AU2A |
| VECO020750 | *Augochlorella* | *striata* | 0 | 1 | 11 | Aug | 2021 | USA | A30 | AU3 |
| VECO020751 | *Augochlorella* | *striata* | 0 | 1 | 11 | Aug | 2021 | USA | A30 | AU4 |
| VECO020752 | *Augochlorella* | *striata* | 1 | 0 | 11 | Aug | 2021 | USA | A30 | AU5A |
| VECO020753 | *Bombus* | *impatiens* | 1 | 0 | 11 | Aug | 2021 | USA | A30 | BO1A |
| VECO020754 | *Bombus* | *impatiens* | 1 | 0 | 11 | Aug | 2021 | USA | A30 | BO2 |
| VECO020755 | *Lasioglossum (Dialictus)* | *imitatum* | 0 | 1 | 11 | Aug | 2021 | USA | A30 | LA1 |
| VECO020756 | *Lasioglossum (Dialictus)* | *apopkense* | 0 | 1 | 11 | Aug | 2021 | USA | A30 | LA2 |
| VECO020757 | *Colletes* | *sp.* | 0 | 1 | 14 | May | 2022 | USA | H98 | AN1 |
| VECO020758 | *Colletes* | *distincta* | 0 | 1 | 14 | May | 2022 | USA | H98 | AN2 |
| VECO020759 | *Lasioglossum (Dialictus)* | *imitatum* | 0 | 1 | 14 | May | 2022 | USA | H98 | LA1 |
| VECO020760 | *Lasioglossum (Dialictus)* | *creberrimum* | 0 | 1 | 2 | Jun | 2022 | USA | MERAMEE | LA1 |
| VECO020761 | *Lasioglossum (Dialictus)* | *imitatum* | 0 | 1 | 2 | Jun | 2022 | USA | MERAMEE | LA2 |

| | | | | | | | | | | |
|---|---|---|---|---|---|---|---|---|---|---|
| VECO020762 | *Augochlorella* | *striata* | 0 | 1 | 2 | Jun | 2022 | USA | MERAMEE | AU1 |
| VECO020763 | *Lasioglossum (Dialictus)* | *imitatum* | 0 | 1 | 4 | Aug | 2021 | USA | NA | LA1 |
| VECO020764 | *Lasioglossum (Dialictus)* | *imitatum* | 0 | 1 | 15 | May | 2021 | USA | ONDA | LA1 |
| VECO020766 | *Augochlorella* | *striata* | 0 | 1 | 2 | Jun | 2022 | USA | SAUK | AU1 |
| VECO020767 | *Colletes* | *aestivalis* | 0 | 1 | 2 | Jun | 2022 | USA | SM | AN1 |
| VECO020768 | *Lasioglossum (Dialictus)* | *imitatum* | 0 | 1 | 2 | Jun | 2022 | USA | SM | LA1 |
| VECO020769 | *Augochlorella* | *striata* | 0 | 1 | 4 | Jun | 2022 | USA | STONE BARN | AU1 |
| VECO020770 | *Augochlorella* | *striata* | 0 | 1 | 4 | Jun | 2022 | USA | STONE BARN | AU2 |
| VECO020771 | *Augochlorella* | *striata* | 0 | 1 | 4 | Jun | 2022 | USA | STONE BARN | AU3 |
| VECO020772 | *Lasioglossum (Dialictus)* | *imitatum* | 0 | 1 | 4 | Jun | 2022 | USA | STONE BARN | LA1 |
| VECO020773 | *Lasioglossum* | *macoupinense* | 0 | 1 | 4 | Jun | 2022 | USA | STONE BARN | LA2 |
| VECO020774 | *Lasioglossum (Dialictus)* | *imitatum* | 0 | 1 | 4 | Jun | 2022 | USA | STONE BARN | LA3 |
| VECO020775 | *Lasioglossum (Dialictus)* | *imitatum* | 0 | 1 | 15 | May | 2021 | USA | ONDA | LA2 |
| VECO020776 | *Lasioglossum (Dialictus)* | *imitatum* | 0 | 1 | 15 | May | 2021 | USA | ONDA | LA2 |
| VECO020777 | *Lasioglossum (Dialictus)* | *imitatum* | 0 | 1 | 15 | May | 2021 | USA | ONDA | LA2 |
| VECO020778 | *Lasioglossum (Dialictus)* | *imitatum* | 0 | 1 | 15 | May | 2021 | USA | ONDA | LA2 |

| | | | | | | | | | | |
|---|---|---|---|---|---|---|---|---|---|---|
| VECO020779 | *Lasioglossum (Hemihalictus)* | *imitatum* | 0 | 1 | 17 | May | 2021 | USA | A34 | LA1A |
| VECO020780 | *Lasioglossum (Dialictus)* | *imitatum* | 0 | 1 | 16 | May | 2021 | USA | A47 | LA1A |
| VECO020781 | *Augochlorella* | *striata* | 1 | 0 | 13 | Aug | 2021 | USA | A30 | SM1A |
| VECO020782 | *Bombus* | *impatiens* | 1 | 0 | 11 | Aug | 2021 | USA | A30 | BO2 |
| VECO020783 | *Augochlora* | *pura* | 0 | 1 | 11 | Aug | 2021 | USA | A30 | AU1 |
| VECO020785 | *Lasioglossum (Dialictus)* | *imitatum* | 0 | 1 | 15 | May | 2021 | USA | ONDA | LA1 |
| VECO022055 | *Lasioglossum (Dialictus)* | *imitatum* | 0 | 1 | 15 | May | 2021 | USA | ONDA | LA1 |
| VECO020765 | *Lasioglossum (Dialictus)* | *imitatum* | 0 | 1 | 15 | May | 2021 | USA | ONDA | LA2 |
| VECO022054 | *Lasioglossum (Dialictus)* | *apopkense* | 0 | 1 | 11 | Aug | 2021 | USA | A30 | LA2 |
| VECO020784 | *Lasioglossum (Dialictus)* | *sp.* | 0 | 1 | 11 | Aug | 2021 | USA | A30 | LA2 |

**Table S4**. Pollinator observation data.

| Locality | Host | Date | Observers | Total time (hrs) | Person-hours | Times | Flowers observed | Inflorescences observed | Pollinators | Observed pollinator number | Pollinator rate per population per hour | Pollinator rate per population per day (8 hours) | Pollinator rate per flower per day (8 hours) | Pollination rate per inflorescence per day (8 hours) | Fruit set | Weather | Persons present |
|---|---|---|---|---|---|---|---|---|---|---|---|---|---|---|---|---|---|
| Yonah Mt, GA | Heuchera americana var. americana | 4/20/21 | 1 | 2 | 2 | 12:30 PM - 2:30 PM | 10 | | None | | | | | | n/a | | Folk |
| Little Park Trail, SC | Heuchera caroliniana | 4/21/21 | 1 | 2 | 2 | 12:30 PM - 2:30 PM | 20 | | None | | | | | | n/a | | Folk |
| Sardis Lake, MS | Heuchera americana var. hirsuticaulis | 5/1/21 | 5 | 1.5 | 7.5 | 3:00 PM - 4:30 PM | 321 | | None | | | | | | 0 | | N.J. Engle-Wrye, H.K. Engle-Wrye, Folk, Siniscalchi, Dahal |
| Tishomingo State Park, MS | Heuchera americana var. americana | 5/7/21 | 1 | 5 | 5 | 8:30 AM - 10:30 AM; 12:30 PM - 3:30 PM | 84 | | None | | | | | | 0.23404255 | First observation sunny but brisk. Second sunny and cool with a light breeze. | Folk |
| Mt Sequoyah Woods Tr, AR | Heuchera americana var. hirsuticaulis | 5/15/21 | 2 | 0.25 | 0.5 | 9:00 AM - 9:15 AM | 210 | 7 | None | | | | | | | Overcast | Folk and Engle-Wrye |
| Lake Wilson | Heuchera americana var. hirsuticaulis | 5/15/21 | 2 | 1.5 | 3 | 10:00 AM - 11:30 AM | 100 | | None | | | | | | 0 | Cool overcast, later fitful sun. Fruit set unclear -- early. | Folk and Engle-Wrye |
| Onda Mountain Road | Heuchera americana var. hirsuticaulis | 5/15/21 | 2 | 2 | 4 | 2:00 PM - 4:00 PM | 6,200 | 124 | Lasioglossum (32), syrphid (2), Augochlorini (1), orange-black wasp (1) | 40 | 20 | 160 | 0.02580645 | 1.29032258 | 1 | Fitfully sunny, very gentle breeze | Folk and Engle-Wrye |
| Lake Wilson | Heuchera americana var. hirsuticaulis | 5/16/21 | 2 | 1 | 2 | 11:45 AM - 12:45 PM | 116 | 7 | Lasioglossum (12) | 12 | 12 | 96 | 0.82758621 | 13.7142857 | 0 | Mostly overcast but fitfully sunny | Folk and Engle-Wrye |
| Cunningham Road A35 | Heuchera americana var. americana | 5/17/21 | 2 | 2 | 4 | 10:00 AM - 12:00 PM | 600 | 12 | Lasioglossum (4) | 4 | 2 | 16 | 0.02666667 | 1.33333333 | 0.73684211 | Mostly overcast but fitfully sunny | Folk and Engle-Wrye |
| Franklin Road A34 | Heuchera americana var. americana | 5/17/21 | 2 | 3.5 | 7 | 1:15 PM - 4:45 PM | 1150 | 23 | Lasioglossum (11), Augochlorini (9) | 20 | 5.71428571 | 45.7142857 | 0.03975155 | 1.98757764 | 0.32394366 | Mostly overcast changing to fitfully sunny. | Folk and Engle-Wrye |

| | | | | | | | | | | | | | | | | | |
|---|---|---|---|---|---|---|---|---|---|---|---|---|---|---|---|---|---|
| Cleveland Botanical Garden | Heuchera villosa var. macrorhiza | 8/4/21 | 1 | 0.25 | 0.25 | 11:15 AM - 11:30 AM | 500 | 10 | Lasioglossum (5) | 5 | 20 | 160 | 0.32 | 16 | not taken | Sunny, moderate temperature | Folk |
| Cleveland Botanical Garden | Heuchera villosa var. villosa | 8/4/21 | 1 | 0.25 | 0.25 | 11:30 AM - 11:45 AM | 500 | 10 | Lasioglossum (5) | 5 | 20 | 160 | 0.32 | 16 | not taken | Sunny, moderate temperature | Folk |
| 1285 Neptune Ave, OH | Heuchera villosa var. villosa | 8/4/21 | 1 | 0.25 | 0.25 | 4:00 PM - 4:15 PM | 100 | 3 | Lasioglossum (3) | 3 | 12 | 96 | 0.48 | 32 | not taken | Sunny, moderate temperature | Folk |
| H30 Brogdan Hollow, JP Coleman State Park, Upstream Site | Heuchera villosa var. macrorhiza | 8/11/21 | 1 | 1 | 1 | 11:20 AM - 12:20 PM | 1300 | 13 | Bombus (15), long metallic bee [Augochlora pura] (6), Augochlorini [A. persimilis] (1), tiny Lasioglossum (8), long red eyed dipteran (1) | 31 | 31 | 248 | 0.496 | 24.8 | not taken | Sunny and very warm, a few very brief periods of cloud cover | Folk |
| H30 Brogdan Hollow, JP Coleman State Park, Downstream Site | Heuchera villosa var. macrorhiza | 8/11/21 | 1 | 0.5 | 0.5 | 12:45 PM - 1:15 PM | 3200 | 32 | Bombus (5), long metallic bee [Augochlora pura] (11), Augochlorini [A. persimilis] (1), tiny Lasioglossum (20) | 37 | 74 | 592 | 1.184 | 59.2 | 0.95652174 | Sunny and very warm | Folk |
| H30 Brogdan Hollow, JP Coleman State Park, Downstream Site | Heuchera villosa var. macrorhiza | 8/13/21 | 1 | 2 | 2 | 12:30 PM - 2:30 PM | 3200 | 32 | Long metallic bee [Augochlora pura] (22), Augochlorini [A. persimilis] (1), unknown small metallic bee (2), tiny Lasioglossum (3) | 28 | 14 | 112 | 0.224 | 11.2 | not taken | Sunny until 12:33, then a hard rain for ten minutes (bees cleared out in 2), and bees back by 1:20 before the flowers were quite dry. Another very light rain at 1:25 to 1:30. Dreary clouds with occasional bleary sunlight. | Folk |
| Downtown Minneapolis | Heuchera villosa var. macrorhiza | 10/7/21 | 1 | 0.25 | 0.25 | 3:15 PM - 3:30 PM | 500 | 10 | Bombus (3) | 3 | 12 | 96 | 0.192 | 9.6 | not taken | Sunny, cool | Folk |
| A3 Clay Creek at CR 55 | Heuchera americana var. americana | 5/2/22 | 2 | 2 | 4 | 12:20 PM - 2:20 PM | 1820 | 26 | Augochlorini (5), small moth (1) | 6 | 3 | 24 | 0.048 | 2.4 | 0.42857143 | Fitfully sunny, warm, almost no wind | Engle-Wrye and Folk |
| A2 Cheaha Mountain | Heuchera americana var. americana | 5/2/22 | 2 | 1.58333333 | 3.16666667 | 5:50 PM - 7:25 PM | 420 | 10 | None | 0 | 0 | 0 | 0 | 0 | | 70 F, no wind, fitfully sunny | Engle-Wrye and Folk |

| | | | | | | | | | | | | | | | | | |
|---|---|---|---|---|---|---|---|---|---|---|---|---|---|---|---|---|---|
| A2 Cheaha Mountain | Heuchera americana var. americana | 5/3/22 | 2 | 1 | 2 | 7:20 AM - 8:20 AM | 420 | 10 | None | 0 | 0 | 0 | 0 | 0 | | 60 F, moderately windy, sunny | Engle-Wrye and Folk |
| A1 Pine Glen Campground | Heuchera americana var. americana | 5/3/22 | 2 | 2.21666667 | 4.43333333 | 11:45 AM - 1:58 PM | 660 | 33 | Augochlorini (13), small bumblebee [Colletes] (9), thin black beetle (1) | 23 | 10.3759399 | 83.0075188 | 0.16601504 | 8.30075188 | 0.9 | 75 F, fitfully sunny, slightest breeze | Engle-Wrye and Folk |
| A53 Henry Creek | Heuchera americana var. americana | 5/3/22 | 2 | 1 | 2 | 2:45 PM - 3:45 PM | 750 | 25 | Augochlorini (1), small black syrphid [?] (1). | 2 | 2 | 16 | 0.032 | 1.6 | | 75 F, fitfully sunny, slightest breeze | Engle-Wrye and Folk |
| A1 Pine Glen Campground | Heuchera americana var. americana | 5/3/22 | 2 | 0.58333333 | 1.16666667 | 9:20 PM - 9:55 PM | 660 | 33 | None | 0 | 0 | 0 | 0 | 0 | | Fully dark, 70 F, slightest breeze. | Engle-Wrye and Folk |
| A1 Pine Glen Campground | Heuchera americana var. americana | 5/4/22 | 2 | 0.25 | 0.5 | 6:45 AM - 7:00 AM | 660 | 33 | None | 0 | 0 | 0 | 0 | 0 | | 68 F, scattered clouds, slightest breeze | Engle-Wrye and Folk |
| A54 Ivy Branch | Heuchera americana var. americana | 5/4/22 | 2 | 1 | 2 | 10:55 AM - 11:55 AM | 120 | 4 | None | 0 | 0 | 0 | 0 | 0 | 0.1025641 | 74 F, sunny, site is part shade, almost no wind | Engle-Wrye and Folk |
| Oak Mountain Park Rd | Heuchera americana var. americana | 5/4/22 | 2 | 1 | 2 | 1:15 PM - 2:15 PM | 175 | 7 | None | 0 | 0 | 0 | 0 | 0 | 0.50746269 | 83 F, full sun, sight was part shade | Engle-Wrye and Folk |
| A55 Ocoee Gorge | Heuchera americana var. americana | 5/10/22 | 2 | 0.5 | 1 | 5:50 PM - 6:20 PM | 9310 | 133 | Syrphids (4) | 4 | 8 | 64 | 0.128 | 6.4 | 0.88 | Sunny and warm, 78 F, slightest breeze | Engle-Wrye and Folk |
| A56 Valley River | Heuchera americana var. americana | 5/11/22 | 2 | 0.5 | 1 | 8:40 AM - 9:10 AM | 750 | 25 | None | 0 | 0 | 0 | 0 | 0 | 0.8452381 | Sunny, cool 72 F, slightest breeze | Engle-Wrye and Folk |
| A57 Deep Creek Trail | Heuchera americana var. americana | 5/11/22 | 2 | 1 | 2 | 11:30 AM - 12:30 PM | 1375 | 55 | Lasioglossum (143) | 143 | 143 | 1144 | 2.288 | 114.4 | 0.97142857 | Sunny, warm, 78 F, almost no breeze | Engle-Wrye and Folk |
| A57 Deep Creek Trail | Heuchera americana var. americana | 5/11/22 | 2 | 0.41666667 | 0.83333333 | 12:45 PM - 1:10 PM | 1375 | 55 | Lasioglossum (14), Colletes (3) | 17 | 40.8 | 326.4 | 0.6528 | 32.64 | | Sunny, warm, 78 F, almost no breeze | Engle-Wrye and Folk |
| A58 Canada Rd | Heuchera americana var. americana | 5/11/22 | 2 | 1.08333333 | 2.16666667 | 2:40 PM - 3:45 PM | 180 | 9 | Lasioglossum (9), Colletes (23), syrphid (1) | 33 | 30.4615385 | 243.692308 | 0.48738462 | 24.3692308 | 1 | Sunny, warm, 83 F, almost no breeze | Engle-Wrye and Folk |
| Craven Gap Trail | Heuchera americana var. americana | 5/12/22 | 2 | 0.25 | 0.5 | 12:05 PM - 12:20 PM | 80 | 4 | Colletes (2), Lasioglossum (2) | 4 | 16 | 128 | 0.256 | 12.8 | | 75 F, fitfully sunny, almost no wind | Engle-Wrye and Folk |
| A59 Buck Creek | Heuchera americana var. americana | 5/12/22 | 2 | 0.58333333 | 1.16666667 | 3:20 PM - 3:55 PM | 180 | 9 | Colletes (8) | 8 | 13.7142857 | 109.714286 | 0.21942857 | 10.9714286 | | Fitfully sunny/shade, moderate breeze. | Engle-Wrye and Folk |

| | | | | | | | | | | | | | | | | | |
|---|---|---|---|---|---|---|---|---|---|---|---|---|---|---|---|---|---|
| A60 Rocky Face Park | Heuchera americana var. americana | 5/13/22 | 2 | 3 | 6 | 11:00 AM - 2:00 PM | 1560 | 39 | Colletes (44), Ichneumonid wasp (1) | 45 | 15 | 120 | 0.24 | 12 | 0.97916667 | Cloudy, just rained, transition to overcast at 11:15 AM, when it was 68 degrees F, then transitioned to fitfully sunny at 11:35 AM. | Engle-Wrye and Folk |
| H98 Hwy 209 | Heuchera longiflora var. aceroides | 5/14/22 | 2 | 1 | 2 | 9:45 AM - 10:45 PM | 325 | 13 | Colletes (4), Lasioglossum (2) | 6 | 6 | 48 | 0.096 | 4.8 | 0.75 | Sunny, moderate breeze, 78 degrees F | Engle-Wrye and Folk |
| A62 Porters Creek | Heuchera americana var. americana | 5/14/22 | 2 | 1 | 2 | 3:00 PM - 4:00 PM | 390 | 13 | Lasioglossum (7), Colletes (1), syrphid (1) | 9 | 9 | 72 | 0.144 | 7.2 | 0.625 | Fitfully sunny becoming overcast, 75 degrees F | Engle-Wrye and Folk |
| A63 Schoolhouse Gap Rd | Heuchera americana var. americana | 5/15/22 | 2 | 2.08333333 | 4.16666667 | 8:00 AM - 10:05 AM | 300 | 12 | Black syrphid (2) | 2 | 0.96 | 7.68 | 0.01536 | 0.768 | 0.92105263 | Cloudy becoming sunny but site completely shaded in the morning, slightest breeze, 68 F | Engle-Wrye and Folk |
| A55 Ocoee Gorge | Heuchera americana var. americana | 5/15/22 | 2 | 1 | 2 | 12:35 PM - 1:35 PM | 9310 | 133 | Lasioglossum (10), Colletes (12), ichneumonid wasp (4), black moth with orange markings (1), syrphid (1) | 28 | 28 | 224 | 0.448 | 22.4 | see above | 79 F, sunny, partly shaded site, slightest breeze but with substantial car wind | Engle-Wrye and Folk |
| Taum Sauk Mountain | Heuchera americana var. hirsuticaulis | 6/2/22 | 2 | 1.06666667 | 2.13333333 | 10:20 AM - 11:24 AM | 80 | 4 | Colletes (2), Lasioglossum (1), Augochlorini (1) | 4 | 3.75 | 30 | 0.06 | 3 | 0.40414508 | sunny, 75 F, slight breeze | Engle-Wrye and Folk |
| Camper Spring/ meramec river | Heuchera americana var. hirsuticaulis | 6/2/22 | 2 | 1.06666667 | 2.13333333 | 1:55 PM - 2:59 PM | 300 | 20 | Lasioglossum (17), Augochlorini (3) | 20 | 18.75 | 150 | 0.3 | 15 | 0.61016949 | 78 F, slightest to no breeze, slightly overcast and fitfully sunny | Engle-Wrye and Folk |
| H167 River trail | Heuchera richardsonii | 6/4/22 | 2 | 1.01666667 | 2.03333333 | 10:20 AM - 11:21 | 350 | 14 | Lasioglossum (49), Augochlorini (2) | 51 | 50.1639344 | 401.311475 | 0.80262295 | 40.1311475 | 0.83333333 | 70 F, overcast to fitfully sunny, slightest breeze | Engle-Wrye and Folk |
| Stone Barn Savanna | Heuchera richardsonii | 6/4/22 | 2 | 1 | 2 | 2:25 PM - 3:25 PM | 750 | 25 | Colletes (2), Augochlorini (11), Lasioglossum (6) | 19 | 19 | 152 | 0.304 | 15.2 | 0.51282051 | 61 F, overcast, moderate breeze, | Engle-Wrye and Folk |

| | | | | | | | | | | | | | | | | | |
|---|---|---|---|---|---|---|---|---|---|---|---|---|---|---|---|---|---|
| | | | | | | | | | | | | | | | | then fitfully sunny moderately breezy | |
| A16 Devil's Lake | Heuchera richardsonii | 6/5/22 | 2 | 3 | 6 | 10:45 AM - 1:45 PM | 846 | 47 | Colletes (42), Ichneumonid (2), Lasioglossum (5) | 49 | 16.3333333 | 130.666667 | 0.26133333 | 13.0666667 | 0.74193548 | 60 F overcast, slightest to no breeze. 67 F fitfully sunny, slight breeze. 70 F partly to fully sunny with slight wind. | Engle-Wrye and Folk |
| A64 Bardon Peak Overlook | Heuchera richardsonii | 6/6/22 | 2 | 0.58333333 | 1.16666667 | 3:45 PM - 4:20 PM | 45 | 9 | Ant robbing nectar sometimes touching stamens (6) | 6 | 10.2857143 | 82.2857143 | 0.16457143 | 8.22857143 | too early | 60 F, sunny, windy | Engle-Wrye and Folk |
| A17 Ice Age Natl senic trail | Heuchera richardsonii | 6/7/22 | 2 | 2 | 4 | 10:30 AM - 12:30 PM | 2050 | 82 | Lasioglossum (56), Augochlorini (2), Colletes (3) | 61 | 30.5 | 244 | 0.488 | 24.4 | too early | 65 F, sunny, slight breeze | Engle-Wrye and Folk |
| H166 Ledges state park | Heuchera richardsonii | 6/8/22 | 2 | 2.5 | 5 | 8:40 AM - 11:10 AM | 30 | 2 | Lasioglossum (2) | 2 | 0.8 | 6.4 | 0.0128 | 0.64 | too early | 70 F, almost no breeze, fitfully sunny. slight breeze after 10:10 | Engle-Wrye and Folk |
| A44 3 creeks | Heuchera americana var. hirsuticaulis | 6/8/22 | 2 | 0.53333333 | 1.06666667 | 4:45 PM - 5:17 PM | 38 | 4 | 0 | 0 | 0 | 0 | 0 | 0 | 0.54901961 | 70 F, fitfully sunny to shady, slightest breeze | Engle-Wrye and Folk |

**Table S5**. Nectar data.

| Old Species IDs from notes | Updated species (for pollination paper) | Accession | Nectar (mm) | Average nectar (mm) | Inner diameter | Average nectar (uL) | Date | Time | Experiment type |
|---|---|---|---|---|---|---|---|---|---|
| Heuchera caroliniana | Heuchera caroliniana | A50 | 0.3 | 0.3 | 0.4 | 0.037699112 | 4/21/21 | 12:30 | Wild |
| Heuchera caroliniana | Heuchera caroliniana | A50 | 0.3 | | | | 4/21/21 | 12:30 | Wild |
| Heuchera caroliniana | Heuchera caroliniana | A50 | 0.3 | | | | 4/21/21 | 12:30 | Wild |
| Heuchera americana var. hirsuticaulis | Heuchera americana var. hirsuticaulis | None (Sardis Lake) | 0 | 0 | 0.4 | 0 | 5/1/21 | 3:00 PM | Wild |
| Heuchera americana var. hirsuticaulis | Heuchera americana var. hirsuticaulis | None (Sardis Lake) | 0 | | | | 5/1/21 | 3:00 PM | Wild |
| Heuchera americana var. hirsuticaulis | Heuchera americana var. hirsuticaulis | None (Sardis Lake) | 0 | | | | 5/1/21 | 3:00 PM | Wild |
| Heuchera americana var. hirsuticaulis | Heuchera americana var. hirsuticaulis | None (Sardis Lake) | 0 | | | | 5/1/21 | 3:00 PM | Wild |
| Heuchera americana var. hirsuticaulis | Heuchera americana var. hirsuticaulis | None (Sardis Lake) | 0 | | | | 5/1/21 | 3:00 PM | Wild |
| Heuchera americana var. hirsuticaulis | Heuchera americana var. hirsuticaulis | None (Sardis Lake) | 0 | | | | 5/1/21 | 3:00 PM | Wild |
| Heuchera alba | Heuchera alba | E306 | 18.5 | 20.7 | 0.4 | 2.601238717 | 5/5/21 | 9:30 AM | Greenhouse |
| Heuchera alba | Heuchera alba | E306 | 14 | | | | 5/5/21 | 9:30 AM | Greenhouse |
| Heuchera alba | Heuchera alba | E306 | 7 | | | | 5/5/21 | 9:30 AM | Greenhouse |
| Heuchera alba | Heuchera alba | E306 | 25.5 | | | | 5/5/21 | 9:30 AM | Greenhouse |
| Heuchera alba | Heuchera alba | E306 | 38.5 | | | | 5/5/21 | 9:30 AM | Greenhouse |
| Heuchera longiflora var. aceroides | Heuchera longiflora var. aceroides | H99 | 5.5 | 5.1 | 0.4 | 0.640884901 | 5/5/21 | 9:30 AM | Greenhouse |
| Heuchera longiflora var. aceroides | Heuchera longiflora var. aceroides | H99 | 7 | | | | 5/5/21 | 9:30 AM | Greenhouse |
| Heuchera longiflora var. aceroides | Heuchera longiflora var. aceroides | H99 | 2 | | | | 5/5/21 | 9:30 AM | Greenhouse |
| Heuchera longiflora var. aceroides | Heuchera longiflora var. aceroides | H99 | 6 | | | | 5/5/21 | 9:30 AM | Greenhouse |
| Heuchera longiflora var. aceroides | Heuchera longiflora var. aceroides | H99 | 5 | | | | 5/5/21 | 9:30 AM | Greenhouse |
| Heuchera richardsonii | Heuchera richardsonii | A21-3 | 8 | 6.8 | 0.4 | 0.854513202 | 5/5/21 | 9:30 AM | Greenhouse |

| | | | | | | | | | |
|---|---|---|---|---|---|---|---|---|---|
| Heuchera richardsonii | Heuchera richardsonii | A21-3 | 3 | | | | 5/5/21 | 9:30 AM | Greenhouse |
| Heuchera richardsonii | Heuchera richardsonii | A21-3 | 6 | | | | 5/5/21 | 9:30 AM | Greenhouse |
| Heuchera richardsonii | Heuchera richardsonii | A21-3 | 11 | | | | 5/5/21 | 9:30 AM | Greenhouse |
| Heuchera richardsonii | Heuchera richardsonii | A21-3 | 6 | | | | 5/5/21 | 9:30 AM | Greenhouse |
| Heuchera americana var. americana | Heuchera americana var. americana | A10-1 | 5.5 | 6.85 | 0.4 | 0.860796387 | 5/5/21 | 9:30 AM | Greenhouse |
| Heuchera americana var. americana | Heuchera americana var. americana | A10-1 | 9 | | | | 5/5/21 | 9:30 AM | Greenhouse |
| Heuchera americana var. americana | Heuchera americana var. americana | A10-1 | 4.75 | | | | 5/5/21 | 9:30 AM | Greenhouse |
| Heuchera americana var. americana | Heuchera americana var. americana | A10-1 | 5 | | | | 5/5/21 | 9:30 AM | Greenhouse |
| Heuchera americana var. americana | Heuchera americana var. americana | A10-1 | 10 | | | | 5/5/21 | 9:30 AM | Greenhouse |
| Heuchera americana var. hirsuticaulis | Heuchera americana var. hirsuticaulis | A38-1 | 0 | 0 | 0.4 | 0 | 5/5/21 | 9:30 AM | Greenhouse |
| Heuchera americana var. hirsuticaulis | Heuchera americana var. hirsuticaulis | A38-1 | 0 | | | | 5/5/21 | 9:30 AM | Greenhouse |
| Heuchera americana var. hirsuticaulis | Heuchera americana var. hirsuticaulis | A38-1 | 0 | | | | 5/5/21 | 9:30 AM | Greenhouse |
| Heuchera americana var. hirsuticaulis | Heuchera americana var. hirsuticaulis | A38-1 | 0 | | | | 5/5/21 | 9:30 AM | Greenhouse |
| Heuchera americana var. hirsuticaulis | Heuchera americana var. hirsuticaulis | A38-1 | 0 | | | | 5/5/21 | 9:30 AM | Greenhouse |
| Heuchera americana var. hirsuticaulis | Heuchera americana var. hirsuticaulis | A9-2 | 0 | 0 | 0.4 | 0 | 5/5/21 | 9:30 AM | Greenhouse |
| Heuchera americana var. hirsuticaulis | Heuchera americana var. hirsuticaulis | A9-2 | 0 | | | | 5/5/21 | 9:30 AM | Greenhouse |
| Heuchera americana var. hirsuticaulis | Heuchera americana var. hirsuticaulis | A9-2 | 0 | | | | 5/5/21 | 9:30 AM | Greenhouse |
| Heuchera americana var. hirsuticaulis | Heuchera americana var. hirsuticaulis | A9-2 | 0 | | | | 5/5/21 | 9:30 AM | Greenhouse |
| Heuchera americana var. hirsuticaulis | Heuchera americana var. hirsuticaulis | A9-2 | 0 | | | | 5/5/21 | 9:30 AM | Greenhouse |
| Heuchera americana var. americana | Heuchera americana var. americana | A8 | 7 | 3.7 | 0.4 | 0.464955713 | 5/5/21 | 10:00 AM | Greenhouse |
| Heuchera americana var. americana | Heuchera americana var. americana | A8 | 2 | | | | 5/5/21 | 10:00 AM | Greenhouse |
| Heuchera americana var. americana | Heuchera americana var. americana | A8 | 3 | | | | 5/5/21 | 10:00 AM | Greenhouse |
| Heuchera americana var. americana | Heuchera americana var. americana | A8 | 3 | | | | 5/5/21 | 10:00 AM | Greenhouse |

| | | | | | | | | | |
|---|---|---|---|---|---|---|---|---|---|
| Heuchera americana var. americana | Heuchera americana var. americana | A8 | 3.5 | | | | 5/5/21 | 10:00 AM | Greenhouse |
| Heuchera americana var. hirsuticaulis | Heuchera americana var. hirsuticaulis | A49-1 | 1 | 1.2 | 0.4 | 0.150796447 | 5/5/21 | 10:00 AM | Greenhouse |
| Heuchera americana var. hirsuticaulis | Heuchera americana var. hirsuticaulis | A49-1 | 2 | | | | 5/5/21 | 10:00 AM | Greenhouse |
| Heuchera americana var. hirsuticaulis | Heuchera americana var. hirsuticaulis | A49-1 | 0 | | | | 5/5/21 | 10:00 AM | Greenhouse |
| Heuchera americana var. hirsuticaulis | Heuchera americana var. hirsuticaulis | A49-1 | 2 | | | | 5/5/21 | 10:00 AM | Greenhouse |
| Heuchera americana var. hirsuticaulis | Heuchera americana var. hirsuticaulis | A49-1 | 1 | | | | 5/5/21 | 10:00 AM | Greenhouse |
| Heuchera soltisii | Heuchera soltisii | I85 | 7.5 | 6.35 | 0.4 | 0.797964534 | 5/5/21 | 3:00 PM | Greenhouse |
| Heuchera soltisii | Heuchera soltisii | I85 | 5.75 | | | | 5/5/21 | 3:00 PM | Greenhouse |
| Heuchera soltisii | Heuchera soltisii | I85 | 3 | | | | 5/5/21 | 3:00 PM | Greenhouse |
| Heuchera soltisii | Heuchera soltisii | I85 | 6 | | | | 5/5/21 | 3:00 PM | Greenhouse |
| Heuchera soltisii | Heuchera soltisii | I85 | 9.5 | | | | 5/5/21 | 3:00 PM | Greenhouse |
| Heuchera wootonii | Heuchera wootonii | H22 | 8 | 10.3 | 0.4 | 1.294336173 | 5/5/21 | 3:00 PM | Greenhouse |
| Heuchera wootonii | Heuchera wootonii | H22 | 11.25 | | | | 5/5/21 | 3:00 PM | Greenhouse |
| Heuchera wootonii | Heuchera wootonii | H22 | 10.75 | | | | 5/5/21 | 3:00 PM | Greenhouse |
| Heuchera wootonii | Heuchera wootonii | H22 | 10.5 | | | | 5/5/21 | 3:00 PM | Greenhouse |
| Heuchera wootonii | Heuchera wootonii | H22 | 11 | | | | 5/5/21 | 3:00 PM | Greenhouse |
| Heuchera elegans | Heuchera elegans | I42 | 0 | 0 | 0.4 | 0 | 5/5/21 | 3:00 PM | Greenhouse |
| Heuchera elegans | Heuchera elegans | I42 | 0 | | | | 5/5/21 | 3:00 PM | Greenhouse |
| Heuchera elegans | Heuchera elegans | I42 | 0 | | | | 5/5/21 | 3:00 PM | Greenhouse |
| Heuchera elegans | Heuchera elegans | I42 | 0 | | | | 5/5/21 | 3:00 PM | Greenhouse |
| Heuchera elegans | Heuchera elegans | I42 | 0 | | | | 5/5/21 | 3:00 PM | Greenhouse |
| Heuchera rubescens | Heuchera rubescens | I112 | 9.75 | 18.99 | 0.4 | 2.38635378 | 5/5/21 | 3:00 PM | Greenhouse |
| Heuchera rubescens | Heuchera rubescens | I112 | 11 | | | | 5/5/21 | 3:00 PM | Greenhouse |

| | | | | | | | | | |
|---|---|---|---|---|---|---|---|---|---|
| Heuchera rubescens | Heuchera rubescens | I112 | 21 | | | | 5/5/21 | 3:00 PM | Greenhouse |
| Heuchera rubescens | Heuchera rubescens | I112 | 28.7 | | | | 5/5/21 | 3:00 PM | Greenhouse |
| Heuchera rubescens | Heuchera rubescens | I112 | 24.5 | | | | 5/5/21 | 3:00 PM | Greenhouse |
| Heuchera brevistaminea | Heuchera brevistaminea | H37 | 0 | 0 | 0.4 | 0 | 5/5/21 | 3:00 PM | Greenhouse |
| Heuchera brevistaminea | Heuchera brevistaminea | H37 | 0 | | | | 5/5/21 | 3:00 PM | Greenhouse |
| Heuchera brevistaminea | Heuchera brevistaminea | H37 | 0 | | | | 5/5/21 | 3:00 PM | Greenhouse |
| Heuchera brevistaminea | Heuchera brevistaminea | H37 | 0 | | | | 5/5/21 | 3:00 PM | Greenhouse |
| Heuchera brevistaminea | Heuchera brevistaminea | H37 | 0 | | | | 5/5/21 | 3:00 PM | Greenhouse |
| Heuchera caespitosa | Heuchera caespitosa | H48 | 0 | 0 | 0.4 | 0 | 5/5/21 | 3:00 PM | Greenhouse |
| Heuchera caespitosa | Heuchera caespitosa | H48 | 0 | | | | 5/5/21 | 3:00 PM | Greenhouse |
| Heuchera caespitosa | Heuchera caespitosa | H48 | 0 | | | | 5/5/21 | 3:00 PM | Greenhouse |
| Heuchera caespitosa | Heuchera caespitosa | H48 | 0 | | | | 5/5/21 | 3:00 PM | Greenhouse |
| Heuchera caespitosa | Heuchera caespitosa | H48 | 0 | | | | 5/5/21 | 3:00 PM | Greenhouse |
| Heuchera rubescens var. truncata | Heuchera rubescens var. truncata | H153 | 0 | 0.14 | 0.4 | 0.017592919 | 5/5/21 | 3:00 PM | Greenhouse |
| Heuchera rubescens var. truncata | Heuchera rubescens var. truncata | H153 | 0.1 | | | | 5/5/21 | 3:00 PM | Greenhouse |
| Heuchera rubescens var. truncata | Heuchera rubescens var. truncata | H153 | 0.5 | | | | 5/5/21 | 3:00 PM | Greenhouse |
| Heuchera rubescens var. truncata | Heuchera rubescens var. truncata | H153 | 0 | | | | 5/5/21 | 3:00 PM | Greenhouse |
| Heuchera rubescens var. truncata | Heuchera rubescens var. truncata | H153 | 0.1 | | | | 5/5/21 | 3:00 PM | Greenhouse |
| Heuchera mexicana var. mexicana | Heuchera mexicana var. mexicana | I51 | 0 | 0.06 | 0.4 | 0.007539822 | 5/5/21 | 3:00 PM | Greenhouse |
| Heuchera mexicana var. mexicana | Heuchera mexicana var. mexicana | I51 | 0.1 | | | | 5/5/21 | 3:00 PM | Greenhouse |
| Heuchera mexicana var. mexicana | Heuchera mexicana var. mexicana | I51 | 0.1 | | | | 5/5/21 | 3:00 PM | Greenhouse |
| Heuchera mexicana var. mexicana | Heuchera mexicana var. mexicana | I51 | 0.1 | | | | 5/5/21 | 3:00 PM | Greenhouse |
| Heuchera mexicana var. mexicana | Heuchera mexicana var. mexicana | I51 | 0 | | | | 5/5/21 | 3:00 PM | Greenhouse |

| | | | | | | | | | |
|---|---|---|---|---|---|---|---|---|---|
| Heuchera parvifolia | Heuchera parvifolia | E234 | 3 | 3.25 | 0.4 | 0.408407045 | 5/5/21 | 3:30 PM | Greenhouse |
| Heuchera parvifolia | Heuchera parvifolia | E234 | 3.5 | | | | 5/5/21 | 3:30 PM | Greenhouse |
| Heuchera parvifolia | Heuchera parvifolia | E234 | 3.75 | | | | 5/5/21 | 3:30 PM | Greenhouse |
| Heuchera parvifolia | Heuchera parvifolia | E234 | 3 | | | | 5/5/21 | 3:30 PM | Greenhouse |
| Heuchera parvifolia | Heuchera parvifolia | E234 | 3 | | | | 5/5/21 | 3:30 PM | Greenhouse |
| Heuchera richardsonii | Heuchera richardsonii | H105 | 3.5 | 3.5 | 0.4 | 0.439822972 | 5/5/21 | 3:30 PM | Greenhouse |
| Heuchera richardsonii | Heuchera richardsonii | H105 | 5 | | | | 5/5/21 | 3:30 PM | Greenhouse |
| Heuchera richardsonii | Heuchera richardsonii | H105 | 2 | | | | 5/5/21 | 3:30 PM | Greenhouse |
| Heuchera richardsonii | Heuchera richardsonii | H105 | 3 | | | | 5/5/21 | 3:30 PM | Greenhouse |
| Heuchera richardsonii | Heuchera richardsonii | H105 | 4 | | | | 5/5/21 | 3:30 PM | Greenhouse |
| Heuchera richardsonii | Heuchera richardsonii | A42 | 0.1 | 0.19 | 0.4 | 0.023876104 | 5/5/21 | 3:30 PM | Greenhouse |
| Heuchera richardsonii | Heuchera richardsonii | A42 | 0.75 | | | | 5/5/21 | 3:30 PM | Greenhouse |
| Heuchera richardsonii | Heuchera richardsonii | A42 | 0 | | | | 5/5/21 | 3:30 PM | Greenhouse |
| Heuchera richardsonii | Heuchera richardsonii | A42 | 0.1 | | | | 5/5/21 | 3:30 PM | Greenhouse |
| Heuchera richardsonii | Heuchera richardsonii | A42 | 0 | | | | 5/5/21 | 3:30 PM | Greenhouse |
| Heuchera americana var. americana | Heuchera americana var. americana | A28 (Tishomingo State Park) | 0 | 0 | 0.4 | 0 | 5/7/21 | 3:00 PM | Wild |
| Heuchera americana var. americana | Heuchera americana var. americana | A28 (Tishomingo State Park) | 0 | | | | 5/7/21 | 3:00 PM | Wild |
| Heuchera americana var. americana | Heuchera americana var. americana | A28 (Tishomingo State Park) | 0 | | | | 5/7/21 | 3:00 PM | Wild |
| Heuchera americana var. americana | Heuchera americana var. americana | A28 (Tishomingo State Park) | 0 | | | | 5/7/21 | 3:00 PM | Wild |
| Heuchera americana var. americana | Heuchera americana var. americana | A28 (Tishomingo State Park) | 0 | | | | 5/7/21 | 3:00 PM | Wild |
| Heuchera americana var. americana | Heuchera americana var. americana | A28 (Tishomingo State Park) | 0 | | | | 5/7/21 | 3:00 PM | Wild |
| Heuchera americana var. americana | Heuchera americana var. americana | A28 (Tishomingo State Park) | 0 | | | | 5/7/21 | 3:00 PM | Wild |
| Heuchera americana var. americana | Heuchera americana var. americana | A28 (Tishomingo State Park) | 0 | | | | 5/7/21 | 3:00 PM | Wild |

| | | | | | | | | | |
|---|---|---|---|---|---|---|---|---|---|
| Heuchera americana var. americana | Heuchera americana var. americana | A28 (Tishomingo State Park) | 0 | | | | 5/7/21 | 3:00 PM | Wild |
| Heuchera americana var. americana | Heuchera americana var. americana | A28 (Tishomingo State Park) | 0 | | | | 5/7/21 | 3:00 PM | Wild |
| Heuchera americana var. hirsuticaulis | Heuchera americana var. hirsuticaulis | Mt Sequoyah Woods Trail | 3 | 2.05 | 0.4 | 0.257610598 | 5/15/21 | 9:15 AM | Wild |
| Heuchera americana var. hirsuticaulis | Heuchera americana var. hirsuticaulis | Mt Sequoyah Woods Trail | 2 | | | | 5/15/21 | 9:15 AM | Wild |
| Heuchera americana var. hirsuticaulis | Heuchera americana var. hirsuticaulis | Mt Sequoyah Woods Trail | 1.5 | | | | 5/15/21 | 9:15 AM | Wild |
| Heuchera americana var. hirsuticaulis | Heuchera americana var. hirsuticaulis | Mt Sequoyah Woods Trail | 2.75 | | | | 5/15/21 | 9:15 AM | Wild |
| Heuchera americana var. hirsuticaulis | Heuchera americana var. hirsuticaulis | Mt Sequoyah Woods Trail | 1 | | | | 5/15/21 | 9:15 AM | Wild |
| Heuchera americana var. hirsuticaulis | Heuchera americana var. hirsuticaulis | Lake Wilson | 0.75 | 1.75 | 0.4 | 0.219911486 | 5/15/21 | 11:30 AM | Wild |
| Heuchera americana var. hirsuticaulis | Heuchera americana var. hirsuticaulis | Lake Wilson | 2.5 | | | | 5/15/21 | 11:30 AM | Wild |
| Heuchera americana var. hirsuticaulis | Heuchera americana var. hirsuticaulis | Lake Wilson | 3 | | | | 5/15/21 | 11:30 AM | Wild |
| Heuchera americana var. hirsuticaulis | Heuchera americana var. hirsuticaulis | Lake Wilson | 1 | | | | 5/15/21 | 11:30 AM | Wild |
| Heuchera americana var. hirsuticaulis | Heuchera americana var. hirsuticaulis | Lake Wilson | 1.5 | | | | 5/15/21 | 11:30 AM | Wild |
| Heuchera americana var. hirsuticaulis | Heuchera americana var. hirsuticaulis | Lake Wilson | 0 | 0.14 | 0.4 | 0.017592919 | 5/15/21 | 11:30 AM | Wild |
| Heuchera americana var. hirsuticaulis | Heuchera americana var. hirsuticaulis | Lake Wilson | 0.1 | | | | 5/15/21 | 11:30 AM | Wild |
| Heuchera americana var. hirsuticaulis | Heuchera americana var. hirsuticaulis | Lake Wilson | 0.1 | | | | 5/15/21 | 11:30 AM | Wild |
| Heuchera americana var. hirsuticaulis | Heuchera americana var. hirsuticaulis | Lake Wilson | 0 | | | | 5/15/21 | 11:30 AM | Wild |
| Heuchera americana var. hirsuticaulis | Heuchera americana var. hirsuticaulis | Lake Wilson | 0.5 | | | | 5/15/21 | 11:30 AM | Wild |
| Heuchera americana var. hirsuticaulis | Heuchera americana var. hirsuticaulis | Onda Mountain Rd | 1 | 0.85 | 0.4 | 0.10681415 | 5/15/21 | 4:00 PM | Wild |
| Heuchera americana var. hirsuticaulis | Heuchera americana var. hirsuticaulis | Onda Mountain Rd | 1 | | | | 5/15/21 | 4:00 PM | Wild |
| Heuchera americana var. hirsuticaulis | Heuchera americana var. hirsuticaulis | Onda Mountain Rd | 1.25 | | | | 5/15/21 | 4:00 PM | Wild |
| Heuchera americana var. hirsuticaulis | Heuchera americana var. hirsuticaulis | Onda Mountain Rd | 0.5 | | | | 5/15/21 | 4:00 PM | Wild |
| Heuchera americana var. hirsuticaulis | Heuchera americana var. hirsuticaulis | Onda Mountain Rd | 0.5 | | | | 5/15/21 | 4:00 PM | Wild |
| Heuchera americana var. hirsuticaulis | Heuchera americana var. hirsuticaulis | Onda Mountain Rd | 1 | 0.7 | 0.4 | 0.087964594 | 5/15/21 | 4:00 PM | Wild |

| | | | | | | | | | |
|---|---|---|---|---|---|---|---|---|---|
| Heuchera americana var. hirsuticaulis | Heuchera americana var. hirsuticaulis | Onda Mountain Rd | 1 | | | | 5/15/21 | 4:00 PM | Wild |
| Heuchera americana var. hirsuticaulis | Heuchera americana var. hirsuticaulis | Onda Mountain Rd | 0.25 | | | | 5/15/21 | 4:00 PM | Wild |
| Heuchera americana var. hirsuticaulis | Heuchera americana var. hirsuticaulis | Onda Mountain Rd | 0.5 | | | | 5/15/21 | 4:00 PM | Wild |
| Heuchera americana var. hirsuticaulis | Heuchera americana var. hirsuticaulis | Onda Mountain Rd | 0.75 | | | | 5/15/21 | 4:00 PM | Wild |
| Heuchera americana var. americana | Heuchera americana var. americana | Cunningham Rd A35 | 0.25 | 0.8 | 0.4 | 0.100530965 | 5/17/21 | 12:00 PM | Wild |
| Heuchera americana var. americana | Heuchera americana var. americana | Cunningham Rd A35 | 2 | | | | 5/17/21 | 12:00 PM | Wild |
| Heuchera americana var. americana | Heuchera americana var. americana | Cunningham Rd A35 | 1 | | | | 5/17/21 | 12:00 PM | Wild |
| Heuchera americana var. americana | Heuchera americana var. americana | Cunningham Rd A35 | 0.25 | | | | 5/17/21 | 12:00 PM | Wild |
| Heuchera americana var. americana | Heuchera americana var. americana | Cunningham Rd A35 | 0.5 | | | | 5/17/21 | 12:00 PM | Wild |
| Heuchera americana var. americana | Heuchera americana var. americana | Frankin Rd A34 | 0.1 | 0.1 | 0.4 | 0.012566371 | 5/17/21 | 16:45 | Wild |
| Heuchera americana var. americana | Heuchera americana var. americana | Frankin Rd A34 | 0.1 | | | | 5/17/21 | 16:45 | Wild |
| Heuchera americana var. americana | Heuchera americana var. americana | Frankin Rd A34 | 0.1 | | | | 5/17/21 | 16:45 | Wild |
| Heuchera americana var. americana | Heuchera americana var. americana | Frankin Rd A34 | 0.1 | | | | 5/17/21 | 16:45 | Wild |
| Heuchera americana var. americana | Heuchera americana var. americana | Frankin Rd A34 | 0.1 | | | | 5/17/21 | 16:45 | Wild |
| Heuchera americana var. americana | Heuchera americana var. americana | Frankin Rd A34 | 0.1 | 0.19 | 0.4 | 0.023876104 | 5/17/21 | 16:45 | Wild |
| Heuchera americana var. americana | Heuchera americana var. americana | Frankin Rd A34 | 0 | | | | 5/17/21 | 16:45 | Wild |
| Heuchera americana var. americana | Heuchera americana var. americana | Frankin Rd A34 | 0.75 | | | | 5/17/21 | 16:45 | Wild |
| Heuchera americana var. americana | Heuchera americana var. americana | Frankin Rd A34 | 0.1 | | | | 5/17/21 | 16:45 | Wild |
| Heuchera americana var. americana | Heuchera americana var. americana | Frankin Rd A34 | 0 | | | | 5/17/21 | 16:45 | Wild |
| Heuchera glomerulata | Heuchera glomerulata | A23-1 | 3.75 | 3.1 | 0.4 | 0.389557489 | 4/7/22 | 9:10 | Greenhouse |
| Heuchera glomerulata | Heuchera glomerulata | A23-1 | 2.5 | | | | 4/7/22 | 9:10 | Greenhouse |
| Heuchera glomerulata | Heuchera glomerulata | A23-1 | 2.25 | | | | 4/7/22 | 9:10 | Greenhouse |
| Heuchera glomerulata | Heuchera glomerulata | A23-1 | 3 | | | | 4/7/22 | 9:10 | Greenhouse |

| | | | | | | | | | |
|---|---|---|---|---|---|---|---|---|---|
| Heuchera glomerulata | Heuchera glomerulata | A23-1 | 4 | | | | 4/7/22 | 9:10 | Greenhouse |
| Heuchera cylindrica var. cylindrica | Heuchera cylindrica var. cylindrica | E2890-1 | 5.5 | 8.35 | 0.4 | 1.049291946 | 4/7/22 | 9:10 | Greenhouse |
| Heuchera cylindrica var. cylindrica | Heuchera cylindrica var. cylindrica | E2890-1 | 8.5 | | | | 4/7/22 | 9:10 | Greenhouse |
| Heuchera cylindrica var. cylindrica | Heuchera cylindrica var. cylindrica | E2890-1 | 9 | | | | 4/7/22 | 9:10 | Greenhouse |
| Heuchera cylindrica var. cylindrica | Heuchera cylindrica var. cylindrica | E2890-1 | 10.75 | | | | 4/7/22 | 9:10 | Greenhouse |
| Heuchera cylindrica var. cylindrica | Heuchera cylindrica var. cylindrica | E2890-1 | 8 | | | | 4/7/22 | 9:10 | Greenhouse |
| Heuchera parishii | Heuchera parishii | E2895-2 | 0 | 0 | 0.4 | 0 | 4/7/22 | 9:10 | Greenhouse |
| Heuchera parishii | Heuchera parishii | E2895-2 | 0 | | | | 4/7/22 | 9:10 | Greenhouse |
| Heuchera parishii | Heuchera parishii | E2895-2 | 0 | | | | 4/7/22 | 9:10 | Greenhouse |
| Heuchera parishii | Heuchera parishii | E2895-2 | 0 | | | | 4/7/22 | 9:10 | Greenhouse |
| Heuchera parishii | Heuchera parishii | E2895-2 | 0 | | | | 4/7/22 | 9:10 | Greenhouse |
| Mukdenia rossii | Mukdenia rossii | Muk | 0 | 0 | 0.4 | 0 | 4/7/22 | 9:10 | Greenhouse |
| Mukdenia rossii | Mukdenia rossii | Muk | 0 | | | | 4/7/22 | 9:10 | Greenhouse |
| Mukdenia rossii | Mukdenia rossii | Muk | 0 | | | | 4/7/22 | 9:10 | Greenhouse |
| Mukdenia rossii | Mukdenia rossii | Muk | 0 | | | | 4/7/22 | 9:10 | Greenhouse |
| Mukdenia rossii | Mukdenia rossii | Muk | 0 | | | | 4/7/22 | 9:10 | Greenhouse |
| Heuchera hallii | Heuchera hallii | E2899-2 | 2 | 2.25 | 0.4 | 0.282743339 | 4/7/22 | 9:40 | Greenhouse |
| Heuchera hallii | Heuchera hallii | E2899-2 | 2.5 | | | | 4/7/22 | 9:40 | Greenhouse |
| Heuchera abramsii | Heuchera abramsii | E2896-2 | 0 | 0 | 0.4 | 0 | 4/7/22 | 9:40 | Greenhouse |
| Heuchera abramsii | Heuchera abramsii | E2896-2 | 0 | | | | 4/7/22 | 9:40 | Greenhouse |
| Heuchera abramsii | Heuchera abramsii | E2896-2 | 0 | | | | 4/7/22 | 9:40 | Greenhouse |
| Heuchera abramsii | Heuchera abramsii | E2896-2 | 0 | | | | 4/7/22 | 9:40 | Greenhouse |
| Heuchera abramsii | Heuchera abramsii | E2896-2 | 0 | | | | 4/7/22 | 9:40 | Greenhouse |

| | | | | | | | | | |
|---|---|---|---|---|---|---|---|---|---|
| Heuchera cylindrica var. glabella | Heuchera cylindrica var. glabella | E2890-3 | 3.25 | 5.333333333 | 0.4 | 0.670206433 | 4/7/22 | 9:40 | Greenhouse |
| Heuchera cylindrica var. glabella | Heuchera cylindrica var. glabella | E2890-3 | 5.75 | | | | 4/7/22 | 9:40 | Greenhouse |
| Heuchera cylindrica var. glabella | Heuchera cylindrica var. glabella | E2890-3 | 7 | | | | 4/7/22 | 9:40 | Greenhouse |
| Heuchera americana var. hirsuticaulis | Heuchera americana var. hirsuticaulis | A32-2 | 0 | 0 | 0.4 | 0 | 4/7/22 | 9:40 | Greenhouse |
| Heuchera americana var. hirsuticaulis | Heuchera americana var. hirsuticaulis | A32-2 | 0 | | | | 4/7/22 | 9:40 | Greenhouse |
| Heuchera americana var. hirsuticaulis | Heuchera americana var. hirsuticaulis | A32-2 | 0 | | | | 4/7/22 | 9:40 | Greenhouse |
| Heuchera americana var. hirsuticaulis | Heuchera americana var. hirsuticaulis | A32-2 | 0 | | | | 4/7/22 | 9:40 | Greenhouse |
| Heuchera americana var. hirsuticaulis | Heuchera americana var. hirsuticaulis | A32-2 | 0 | | | | 4/7/22 | 9:40 | Greenhouse |
| Heuchera americana var. hirsuticaulis | Heuchera americana var. hirsuticaulis | A9-2 | 0.1 | 0.04 | 0.4 | 0.005026548 | 4/7/22 | 9:40 | Greenhouse |
| Heuchera americana var. hirsuticaulis | Heuchera americana var. hirsuticaulis | A9-2 | 0 | | | | 4/7/22 | 9:40 | Greenhouse |
| Heuchera americana var. hirsuticaulis | Heuchera americana var. hirsuticaulis | A9-2 | 0 | | | | 4/7/22 | 9:40 | Greenhouse |
| Heuchera americana var. hirsuticaulis | Heuchera americana var. hirsuticaulis | A9-2 | 0.1 | | | | 4/7/22 | 9:40 | Greenhouse |
| Heuchera americana var. hirsuticaulis | Heuchera americana var. hirsuticaulis | A9-2 | 0 | | | | 4/7/22 | 9:40 | Greenhouse |
| Heuchera americana var. hirsuticaulis | Heuchera americana var. hirsuticaulis | A46-2 | 0.5 | 0.558333333 | 0.4 | 0.070162236 | 4/7/22 | 9:40 | Greenhouse |
| Heuchera americana var. hirsuticaulis | Heuchera americana var. hirsuticaulis | A46-2 | 0.1 | | | | 4/7/22 | 9:40 | Greenhouse |
| Heuchera americana var. hirsuticaulis | Heuchera americana var. hirsuticaulis | A46-2 | 1.5 | | | | 4/7/22 | 9:40 | Greenhouse |
| Heuchera americana var. hirsuticaulis | Heuchera americana var. hirsuticaulis | A46-2 | 0.75 | | | | 4/7/22 | 9:40 | Greenhouse |
| Heuchera americana var. hirsuticaulis | Heuchera americana var. hirsuticaulis | A46-2 | 0.5 | | | | 4/7/22 | 9:40 | Greenhouse |
| Heuchera americana var. hirsuticaulis | Heuchera americana var. hirsuticaulis | A46-2 | 0 | | | | 4/7/22 | 9:40 | Greenhouse |
| Heuchera longiflora var. longiflora | Heuchera longiflora var. longiflora | Prairie boulder | 1 | 2.95 | 0.4 | 0.370707933 | 4/7/22 | 9:40 | Greenhouse |
| Heuchera longiflora var. longiflora | Heuchera longiflora var. longiflora | Prairie boulder | 3 | | | | 4/7/22 | 9:40 | Greenhouse |
| Heuchera longiflora var. longiflora | Heuchera longiflora var. longiflora | Prairie boulder | 3 | | | | 4/7/22 | 9:40 | Greenhouse |
| Heuchera longiflora var. longiflora | Heuchera longiflora var. longiflora | Prairie boulder | 2.75 | | | | 4/7/22 | 9:40 | Greenhouse |

| | | | | | | | | | |
|---|---|---|---|---|---|---|---|---|---|
| Heuchera longiflora var. longiflora | Heuchera longiflora var. longiflora | Prairie boulder | 5 | | | | 4/7/22 | 9:40 | Greenhouse |
| Heuchera caroliniana | Heuchera caroliniana | A50:D | 0.75 | 0.85 | 0.4 | 0.10681415 | 4/13/22 | 3:50 | Greenhouse |
| Heuchera caroliniana | Heuchera caroliniana | A50:D | 1 | | | | 4/13/22 | 3:50 | Greenhouse |
| Heuchera caroliniana | Heuchera caroliniana | A50:D | 0.75 | | | | 4/13/22 | 3:50 | Greenhouse |
| Heuchera caroliniana | Heuchera caroliniana | A50:D | 1 | | | | 4/13/22 | 3:50 | Greenhouse |
| Heuchera caroliniana | Heuchera caroliniana | A50:D | 0.75 | | | | 4/13/22 | 3:50 | Greenhouse |
| Heuchera longiflora var. longiflora | Heuchera longiflora var. longiflora | A51-1 | 3 | 2.916666667 | 0.4 | 0.366519143 | 4/13/22 | 3:50 | Greenhouse |
| Heuchera longiflora var. longiflora | Heuchera longiflora var. longiflora | A51-1 | 4 | | | | 4/13/22 | 3:50 | Greenhouse |
| Heuchera longiflora var. longiflora | Heuchera longiflora var. longiflora | A51-1 | 1.75 | | | | 4/13/22 | 3:50 | Greenhouse |
| Heuchera americana var. calycosa | Heuchera americana var. americana | A3 | 2.5 | 5.458333333 | 0.4 | 0.685914396 | 5/2/22 | 2:20 | Wild |
| Heuchera americana var. calycosa | Heuchera americana var. americana | A3 | 8.25 | | | | 5/2/22 | 2:20 | Wild |
| Heuchera americana var. calycosa | Heuchera americana var. americana | A3 | 6.5 | | | | 5/2/22 | 2:20 | Wild |
| Heuchera americana var. calycosa | Heuchera americana var. americana | A3 | 11.75 | | | | 5/2/22 | 2:20 | Wild |
| Heuchera americana var. calycosa | Heuchera americana var. americana | A3 | 2.25 | | | | 5/2/22 | 2:20 | Wild |
| Heuchera americana var. calycosa | Heuchera americana var. americana | A3 | 1.5 | | | | 5/2/22 | 2:20 | Wild |
| Heuchera americana var. calycosa | Heuchera americana var. americana | A2 | 0 | 0.208333333 | 0.4 | 0.026179939 | 5/2/22 | 7:20 PM | Wild |
| Heuchera americana var. calycosa | Heuchera americana var. americana | A2 | 0.25 | | | | 5/2/22 | 7:20 PM | Wild |
| Heuchera americana var. calycosa | Heuchera americana var. americana | A2 | 0 | | | | 5/2/22 | 7:20 PM | Wild |
| Heuchera americana var. calycosa | Heuchera americana var. americana | A2 | 0.75 | | | | 5/2/22 | 7:20 PM | Wild |
| Heuchera americana var. calycosa | Heuchera americana var. americana | A2 | 0 | | | | 5/2/22 | 7:20 PM | Wild |
| Heuchera americana var. calycosa | Heuchera americana var. americana | A2 | 0.25 | | | | 5/2/22 | 7:20 PM | Wild |
| Heuchera americana var. calycosa | Heuchera americana var. americana | A53 | 0.25 | 0.27 | 0.4 | 0.033929201 | 5/3/22 | 3:45 PM | Wild |
| Heuchera americana var. calycosa | Heuchera americana var. americana | A53 | 0.5 | | | | 5/3/22 | 3:45 PM | Wild |

| | | | | | | | | | |
|---|---|---|---|---|---|---|---|---|---|
| Heuchera americana var. calycosa | Heuchera americana var. americana | A53 | 0.25 | | | | 5/3/22 | 3:45 PM | Wild |
| Heuchera americana var. calycosa | Heuchera americana var. americana | A53 | 0.25 | | | | 5/3/22 | 3:45 PM | Wild |
| Heuchera americana var. calycosa | Heuchera americana var. americana | A53 | 0.1 | | | | 5/3/22 | 3:45 PM | Wild |
| Heuchera americana var. calycosa | Heuchera americana var. americana | A1 | 1 | 2.0625 | 0.4 | 0.259181394 | 5/4/22 | 7:00 AM | Wild |
| Heuchera americana var. calycosa | Heuchera americana var. americana | A1 | 1 | | | | 5/4/22 | 7:00 AM | Wild |
| Heuchera americana var. calycosa | Heuchera americana var. americana | A1 | 5 | | | | 5/4/22 | 7:00 AM | Wild |
| Heuchera americana var. calycosa | Heuchera americana var. americana | A1 | 1.25 | | | | 5/4/22 | 7:00 AM | Wild |
| Heuchera americana var. calycosa | Heuchera americana var. americana | A54 | 0.75 | 0.17 | 0.4 | 0.02136283 | 5/4/22 | 7:00 AM | Wild |
| Heuchera americana var. calycosa | Heuchera americana var. americana | A54 | 0.1 | | | | 5/4/22 | 7:00 AM | Wild |
| Heuchera americana var. calycosa | Heuchera americana var. americana | A54 | 0 | | | | 5/4/22 | 7:00 AM | Wild |
| Heuchera americana var. calycosa | Heuchera americana var. americana | A54 | 0 | | | | 5/4/22 | 7:00 AM | Wild |
| Heuchera americana var. calycosa | Heuchera americana var. americana | A54 | 0 | | | | 5/4/22 | 7:00 AM | Wild |
| Heuchera americana var. calycosa | Heuchera americana var. americana | Oak Mountain Park Rd | 0.75 | 0.17 | 0.4 | 0.02136283 | 5/4/22 | 2:15 PM | Wild |
| Heuchera americana var. calycosa | Heuchera americana var. americana | Oak Mountain Park Rd | 0.1 | | | | 5/4/22 | 2:15 PM | Wild |
| Heuchera americana var. calycosa | Heuchera americana var. americana | Oak Mountain Park Rd | 0 | | | | 5/4/22 | 2:15 PM | Wild |
| Heuchera americana var. calycosa | Heuchera americana var. americana | Oak Mountain Park Rd | 0 | | | | 5/4/22 | 2:15 PM | Wild |
| Heuchera americana var. calycosa | Heuchera americana var. americana | Oak Mountain Park Rd | 0 | | | | 5/4/22 | 2:15 PM | Wild |
| Heuchera americana var. calycosa | Heuchera americana var. americana | Oak Mountain Park Rd | 0.75 | | | | 5/4/22 | 2:15 PM | Wild |
| Heuchera americana var. heteradenia | Heuchera americana var. americana | A55 | 0.3 | 1.01 | 0.4 | 0.126920343 | 5/10/22 | 6:20 PM | Wild |
| Heuchera americana var. heteradenia | Heuchera americana var. americana | A55 | 0.5 | | | | 5/10/22 | 6:20 PM | Wild |
| Heuchera americana var. heteradenia | Heuchera americana var. americana | A55 | 1.75 | | | | 5/10/22 | 6:20 PM | Wild |
| Heuchera americana var. heteradenia | Heuchera americana var. americana | A55 | 0.75 | | | | 5/10/22 | 6:20 PM | Wild |
| Heuchera americana var. heteradenia | Heuchera americana var. americana | A55 | 1.75 | | | | 5/10/22 | 6:20 PM | Wild |

| | | | | | | | | | |
|---|---|---|---|---|---|---|---|---|---|
| Heuchera americana var. americana | Heuchera americana var. americana | A56 | 0.25 | 0.05 | 0.4 | 0.006283185 | 5/11/22 | 9:10 AM | Wild |
| Heuchera americana var. americana | Heuchera americana var. americana | A56 | 0 | | | | 5/11/22 | 9:10 AM | Wild |
| Heuchera americana var. americana | Heuchera americana var. americana | A56 | 0 | | | | 5/11/22 | 9:10 AM | Wild |
| Heuchera americana var. americana | Heuchera americana var. americana | A56 | 0 | | | | 5/11/22 | 9:10 AM | Wild |
| Heuchera americana var. americana | Heuchera americana var. americana | A56 | 0 | | | | 5/11/22 | 9:10 AM | Wild |
| Heuchera americana var. americana | Heuchera americana var. americana | A58 | 0 | 0 | 0.4 | 0 | 5/11/22 | 3:45 PM | Wild |
| Heuchera americana var. americana | Heuchera americana var. americana | A58 | 0 | | | | 5/11/22 | 3:45 PM | Wild |
| Heuchera americana var. americana | Heuchera americana var. americana | A58 | 0 | | | | 5/11/22 | 3:45 PM | Wild |
| Heuchera americana var. americana | Heuchera americana var. americana | A58 | 0 | | | | 5/11/22 | 3:45 PM | Wild |
| Heuchera americana var. americana | Heuchera americana var. americana | A58 | 0 | | | | 5/11/22 | 3:45 PM | Wild |
| Heuchera americana var. americana | Heuchera americana var. americana | Craven Gap Trail | 0 | 0 | 0.4 | 0 | 5/12/22 | 12:20 PM | Wild |
| Heuchera americana var. americana | Heuchera americana var. americana | Craven Gap Trail | 0 | | | | 5/12/22 | 12:20 PM | Wild |
| Heuchera americana var. americana | Heuchera americana var. americana | Craven Gap Trail | 0 | | | | 5/12/22 | 12:20 PM | Wild |
| Heuchera americana var. americana | Heuchera americana var. americana | Craven Gap Trail | 0 | | | | 5/12/22 | 12:20 PM | Wild |
| Heuchera americana var. americana | Heuchera americana var. americana | Craven Gap Trail | 0 | | | | 5/12/22 | 12:20 PM | Wild |
| Heuchera americana var. americana | Heuchera americana var. americana | A59 | 0 | 0 | 0.4 | 0 | 5/12/22 | 3:55 PM | Wild |
| Heuchera americana var. americana | Heuchera americana var. americana | A59 | 0 | | | | 5/12/22 | 3:55 PM | Wild |
| Heuchera americana var. americana | Heuchera americana var. americana | A59 | 0 | | | | 5/12/22 | 3:55 PM | Wild |
| Heuchera americana var. americana | Heuchera americana var. americana | A59 | 0 | | | | 5/12/22 | 3:55 PM | Wild |
| Heuchera americana var. americana | Heuchera americana var. americana | A59 | 0 | | | | 5/12/22 | 3:55 PM | Wild |
| Heuchera americana var. americana | Heuchera americana var. americana | A60 Rocky Face Park | 0 | 0 | 0.4 | 0 | 5/13/22 | 2:00 PM | Wild |
| Heuchera americana var. americana | Heuchera americana var. americana | A60 Rocky Face Park | 0 | | | | 5/13/22 | 2:00 PM | Wild |
| Heuchera americana var. americana | Heuchera americana var. americana | A60 Rocky Face Park | 0 | | | | 5/13/22 | 2:00 PM | Wild |

| | | | | | | | | | |
|---|---|---|---|---|---|---|---|---|---|
| Heuchera americana var. americana | Heuchera americana var. americana | A60 Rocky Face Park | 0 | | | | 5/13/22 | 2:00 PM | Wild |
| Heuchera americana var. americana | Heuchera americana var. americana | A60 Rocky Face Park | 0 | | | | 5/13/22 | 2:00 PM | Wild |
| Heuchera americana var. aceroides | Heuchera longiflora var. aceroides | H98 | 0 | 0 | 0.4 | 0 | 5/14/22 | 10:45 PM | Wild |
| Heuchera americana var. aceroides | Heuchera longiflora var. aceroides | H98 | 0 | | | | 5/14/22 | 10:45 PM | Wild |
| Heuchera americana var. aceroides | Heuchera longiflora var. aceroides | H98 | 0 | | | | 5/14/22 | 10:45 PM | Wild |
| Heuchera americana var. aceroides | Heuchera longiflora var. aceroides | H98 | 0 | | | | 5/14/22 | 10:45 PM | Wild |
| Heuchera americana var. aceroides | Heuchera longiflora var. aceroides | H98 | 0 | | | | 5/14/22 | 10:45 PM | Wild |
| Heuchera americana var. heteradenia | Heuchera americana var. americana | A62 | 5 | 3.5 | 0.4 | 0.439822972 | 5/14/22 | 4:00 PM | Wild |
| Heuchera americana var. heteradenia | Heuchera americana var. americana | A62 | 4 | | | | 5/14/22 | 4:00 PM | Wild |
| Heuchera americana var. heteradenia | Heuchera americana var. americana | A62 | 1 | | | | 5/14/22 | 4:00 PM | Wild |
| Heuchera americana var. heteradenia | Heuchera americana var. americana | A62 | 1 | | | | 5/14/22 | 4:00 PM | Wild |
| Heuchera americana var. heteradenia | Heuchera americana var. americana | A62 | 6.5 | | | | 5/14/22 | 4:00 PM | Wild |
| Heuchera americana var. heteradenia | Heuchera americana var. americana | A63 Schoolhouse Gap Rd | 0 | 0 | 0.4 | 0 | 5/15/22 | 9:30 AM | Wild |
| Heuchera americana var. heteradenia | Heuchera americana var. americana | A63 Schoolhouse Gap Rd | 0 | | | | 5/15/22 | 9:30 AM | Wild |
| Heuchera americana var. heteradenia | Heuchera americana var. americana | A63 Schoolhouse Gap Rd | 0 | | | | 5/15/22 | 9:30 AM | Wild |
| Heuchera americana var. heteradenia | Heuchera americana var. americana | A63 Schoolhouse Gap Rd | 0 | | | | 5/15/22 | 9:30 AM | Wild |
| Heuchera americana var. heteradenia | Heuchera americana var. americana | A63 Schoolhouse Gap Rd | 0 | | | | 5/15/22 | 9:30 AM | Wild |
| Heuchera americana var. hirsuticaulis | Heuchera americana var. hirsuticaulis | Mt Sequoyah Woods Tr, AR | 3 | 2.05 | 0.4 | 0.257610598 | 5/15/21 | 9:15 AM | Wild |
| Heuchera americana var. hirsuticaulis | Heuchera americana var. hirsuticaulis | Mt Sequoyah Woods Tr, AR | 2 | | | | 5/15/21 | 9:15 AM | Wild |
| Heuchera americana var. hirsuticaulis | Heuchera americana var. hirsuticaulis | Mt Sequoyah Woods Tr, AR | 1.5 | | | | 5/15/21 | 9:15 AM | Wild |
| Heuchera americana var. hirsuticaulis | Heuchera americana var. hirsuticaulis | Mt Sequoyah Woods Tr, AR | 2.75 | | | | 5/15/21 | 9:15 AM | Wild |
| Heuchera americana var. hirsuticaulis | Heuchera americana var. hirsuticaulis | Mt Sequoyah Woods Tr, AR | 1 | | | | 5/15/21 | 9:15 AM | Wild |
| Heuchera americana var. hispida | Heuchera americana var. americana | A59 | 0.5 | 1.91 | 0.4 | 0.240017679 | 5/12/23 | 2:30 PM | Greenhouse |

| | | | | | | | | | |
|---|---|---|---|---|---|---|---|---|---|
| Heuchera americana var. hispida | Heuchera americana var. americana | A59 | 0.3 | | | | 5/12/23 | 2:30 PM | Greenhouse |
| Heuchera americana var. hispida | Heuchera americana var. americana | A59 | 2.5 | | | | 5/12/23 | 2:30 PM | Greenhouse |
| Heuchera americana var. hispida | Heuchera americana var. americana | A59 | 2.75 | | | | 5/12/23 | 2:30 PM | Greenhouse |
| Heuchera americana var. hispida | Heuchera americana var. americana | A59 | 3.5 | | | | 5/12/23 | 2:30 PM | Greenhouse |
| Heuchera americana var. calycosa | Heuchera americana var. americana | A55 | 0.5 | 0.14 | 0.4 | 0.017592919 | 5/12/23 | 2:30 PM | Greenhouse |
| Heuchera americana var. calycosa | Heuchera americana var. americana | A55 | 0.1 | | | | 5/12/23 | 2:30 PM | Greenhouse |
| Heuchera americana var. calycosa | Heuchera americana var. americana | A55 | 0.1 | | | | 5/12/23 | 2:30 PM | Greenhouse |
| Heuchera americana var. calycosa | Heuchera americana var. americana | A55 | 0 | | | | 5/12/23 | 2:30 PM | Greenhouse |
| Heuchera americana var. calycosa | Heuchera americana var. americana | A55 | 0 | | | | 5/12/23 | 2:30 PM | Greenhouse |
| Heuchera richardsonii | Heuchera richardsonii | A64 Duluth | 0.75 | 0.85 | 0.4 | 0.10681415 | 5/12/23 | 2:30 PM | Greenhouse |
| Heuchera richardsonii | Heuchera richardsonii | A64 Duluth | 1 | | | | 5/12/23 | 2:30 PM | Greenhouse |
| Heuchera richardsonii | Heuchera richardsonii | A64 Duluth | 0.5 | | | | 5/12/23 | 2:30 PM | Greenhouse |
| Heuchera richardsonii | Heuchera richardsonii | A64 Duluth | 0.25 | | | | 5/12/23 | 2:30 PM | Greenhouse |
| Heuchera richardsonii | Heuchera richardsonii | A64 Duluth | 1.75 | | | | 5/12/23 | 2:30 PM | Greenhouse |
| Heuchera americana var. heteradenia | Heuchera americana var. americana | A63 | 0 | 0.02 | 0.4 | 0.002513274 | 5/12/23 | 2:30 PM | Greenhouse |
| Heuchera americana var. heteradenia | Heuchera americana var. americana | A63 | 0 | | | | 5/12/23 | 2:30 PM | Greenhouse |
| Heuchera americana var. heteradenia | Heuchera americana var. americana | A63 | 0 | | | | 5/12/23 | 2:30 PM | Greenhouse |
| Heuchera americana var. heteradenia | Heuchera americana var. americana | A63 | 0 | | | | 5/12/23 | 2:30 PM | Greenhouse |
| Heuchera americana var. heteradenia | Heuchera americana var. americana | A63 | 0.1 | | | | 5/12/23 | 2:30 PM | Greenhouse |
| Heuchera americana var. americana | Heuchera americana var. americana | A8 | 2.5 | 2.75 | 0.4 | 0.345575192 | 5/12/23 | 2:30 PM | Greenhouse |
| Heuchera americana var. americana | Heuchera americana var. americana | A8 | 2.5 | | | | 5/12/23 | 2:30 PM | Greenhouse |
| Heuchera americana var. americana | Heuchera americana var. americana | A8 | 2.25 | | | | 5/12/23 | 2:30 PM | Greenhouse |
| Heuchera americana var. americana | Heuchera americana var. americana | A8 | 4 | | | | 5/12/23 | 2:30 PM | Greenhouse |

| | | | | | | | | | |
|---|---|---|---|---|---|---|---|---|---|
| Heuchera americana var. americana | Heuchera americana var. americana | A8 | 2.5 | | | | 5/12/23 | 2:30 PM | Greenhouse |
| Heuchera americana var. hirsuticaulis | Heuchera americana var. hirsuticaulis | Taum Sauk Mountain State Park | 0 | 0 | 0.4 | 0 | 6/2/22 | 11:24 AM | Wild |
| Heuchera americana var. hirsuticaulis | Heuchera americana var. hirsuticaulis | Taum Sauk Mountain State Park | 0 | | | | 6/2/22 | 11:24 AM | Wild |
| Heuchera americana var. hirsuticaulis | Heuchera americana var. hirsuticaulis | Taum Sauk Mountain State Park | 0 | | | | 6/2/22 | 11:24 AM | Wild |
| Heuchera americana var. hirsuticaulis | Heuchera americana var. hirsuticaulis | Taum Sauk Mountain State Park | 0 | | | | 6/2/22 | 11:24 AM | Wild |
| Heuchera americana var. hirsuticaulis | Heuchera americana var. hirsuticaulis | Taum Sauk Mountain State Park | 0 | | | | 6/2/22 | 11:24 AM | Wild |
| Heuchera richardsonii var. affinis | Heuchera americana var. hirsuticaulis | camper spring/ meramec river | 0 | 0 | 0.4 | 0 | 6/2/22 | 2:59 PM | Wild |
| Heuchera richardsonii var. affinis | Heuchera americana var. hirsuticaulis | camper spring/ meramec river | 0 | | | | 6/2/22 | 2:59 PM | Wild |
| Heuchera richardsonii var. affinis | Heuchera americana var. hirsuticaulis | camper spring/ meramec river | 0 | | | | 6/2/22 | 2:59 PM | Wild |
| Heuchera richardsonii var. affinis | Heuchera americana var. hirsuticaulis | camper spring/ meramec river | 0 | | | | 6/2/22 | 2:59 PM | Wild |
| Heuchera richardsonii var. affinis | Heuchera americana var. hirsuticaulis | camper spring/ meramec river | 0 | | | | 6/2/22 | 2:59 PM | Wild |
| Heuchera richardsonii var. affinis | Heuchera richardsonii | H167 Starved rock pool | 0 | 0 | 0.4 | 0 | 6/4/22 | 11:21 AM | Wild |
| Heuchera richardsonii var. affinis | Heuchera richardsonii | H167 Starved rock pool | 0 | | | | 6/4/22 | 11:21 AM | Wild |
| Heuchera richardsonii var. affinis | Heuchera richardsonii | H167 Starved rock pool | 0 | | | | 6/4/22 | 11:21 AM | Wild |
| Heuchera richardsonii var. affinis | Heuchera richardsonii | H167 Starved rock pool | 0 | | | | 6/4/22 | 11:21 AM | Wild |
| Heuchera richardsonii var. affinis | Heuchera richardsonii | H167 Starved rock pool | 0 | | | | 6/4/22 | 11:21 AM | Wild |
| Heuchera richardsonii var. affinis | Heuchera richardsonii | Stone Barn Savanna | 0 | 0 | 0.4 | 0 | 6/4/22 | 3:25 PM | Wild |
| Heuchera richardsonii var. affinis | Heuchera richardsonii | Stone Barn Savanna | 0 | | | | 6/4/22 | 3:25 PM | Wild |
| Heuchera richardsonii var. affinis | Heuchera richardsonii | Stone Barn Savanna | 0 | | | | 6/4/22 | 3:25 PM | Wild |
| Heuchera richardsonii var. affinis | Heuchera richardsonii | Stone Barn Savanna | 0 | | | | 6/4/22 | 3:25 PM | Wild |
| Heuchera richardsonii var. affinis | Heuchera richardsonii | Stone Barn Savanna | 0 | | | | 6/4/22 | 3:25 PM | Wild |
| Heuchera richardsonii var. grayana | Heuchera richardsonii | A16 Devil's Lake | 1.5 | 0.3 | 0.4 | 0.037699112 | 6/5/22 | 1:45 PM | Wild |
| Heuchera richardsonii var. grayana | Heuchera richardsonii | A16 Devil's Lake | 0 | | | | 6/5/22 | 1:45 PM | Wild |

| | | | | | | | | | |
|---|---|---|---|---|---|---|---|---|---|
| Heuchera richardsonii var. grayana | Heuchera richardsonii | A16 Devil's Lake | 0 | | | | 6/5/22 | 1:45 PM | Wild |
| Heuchera richardsonii var. grayana | Heuchera richardsonii | A16 Devil's Lake | 0 | | | | 6/5/22 | 1:45 PM | Wild |
| Heuchera richardsonii var. grayana | Heuchera richardsonii | A16 Devil's Lake | 0 | | | | 6/5/22 | 1:45 PM | Wild |
| Heuchera richardsonii var. grayana | Heuchera richardsonii | A64 Bardon Peak Overlook | 0 | 0 | 0.4 | 0 | 6/6/22 | 3:45 PM | Wild |
| Heuchera richardsonii var. grayana | Heuchera richardsonii | A64 Bardon Peak Overlook | 0 | | | | 6/6/22 | 3:45 PM | Wild |
| Heuchera richardsonii var. grayana | Heuchera richardsonii | A64 Bardon Peak Overlook | 0 | | | | 6/6/22 | 3:45 PM | Wild |
| Heuchera richardsonii var. grayana | Heuchera richardsonii | A64 Bardon Peak Overlook | 0 | | | | 6/6/22 | 3:45 PM | Wild |
| Heuchera richardsonii var. grayana | Heuchera richardsonii | A64 Bardon Peak Overlook | 0 | | | | 6/6/22 | 3:45 PM | Wild |
| Heuchera richardsonii var. grayana | Heuchera richardsonii | A17 Ice Age Natl Scenic Trail | 0 | 0.5 | 0.4 | 0.062831853 | 6/7/22 | 12:00 PM | Wild |
| Heuchera richardsonii var. grayana | Heuchera richardsonii | A17 Ice Age Natl Scenic Trail | 0 | | | | 6/7/22 | 12:00 PM | Wild |
| Heuchera richardsonii var. grayana | Heuchera richardsonii | A17 Ice Age Natl Scenic Trail | 0.75 | | | | 6/7/22 | 12:00 PM | Wild |
| Heuchera richardsonii var. grayana | Heuchera richardsonii | A17 Ice Age Natl Scenic Trail | 0.5 | | | | 6/7/22 | 12:00 PM | Wild |
| Heuchera richardsonii var. grayana | Heuchera richardsonii | A17 Ice Age Natl Scenic Trail | 1.25 | | | | 6/7/22 | 12:00 PM | Wild |
| Heuchera richardsonii var. affinis | Heuchera americana var. hirsuticaulis | A44 | 2.25 | 3.52 | 0.4 | 0.442336246 | 6/8/22 | 4:45 | Wild |
| Heuchera richardsonii var. affinis | Heuchera americana var. hirsuticaulis | A44 | 0.1 | | | | 6/8/22 | 4:45 | Wild |
| Heuchera richardsonii var. affinis | Heuchera americana var. hirsuticaulis | A44 | 2.5 | | | | 6/8/22 | 4:45 | Wild |
| Heuchera richardsonii var. affinis | Heuchera americana var. hirsuticaulis | A44 | 0.25 | | | | 6/8/22 | 4:45 | Wild |
| Heuchera richardsonii var. affinis | Heuchera americana var. hirsuticaulis | A44 | 12.5 | | | | 6/8/22 | 4:45 | Wild |

**Table S6**. Fruit data.

| Old Species IDs from notes | Updated species (for pollination paper) | Accession | Fruits | Aborted | Percentage | Date |
|---|---|---|---|---|---|---|
| Heuchera alba | Heuchera alba | H64 | 14 | 2 | 0.875 | 19 July 2011 |
| Heuchera americana var. americana | Heuchera americana var. americana | A28 | 11 | 36 | 0.2340425532 | 7 May 2021 |
| Heuchera americana var. brevipetela | Heuchera americana var. americana | A35 | 28 | 10 | 0.7368421053 | 5/17/21 |
| Heuchera americana | Heuchera americana var. americana | A34 | 48 | 23 | 0.676056338 | 5/17/21 |
| Heuchera americana var. calycosa | Heuchera americana var. americana | A3 | 15 | 20 | 0.4285714286 | 5/2/22 |
| Heuchera americana var. calycosa | Heuchera americana var. americana | A1 | 18 | 2 | 0.9 | 5/3/22 |
| Heuchera americana var. calycosa | Heuchera americana var. americana | A54 | 4 | 35 | 0.1025641026 | 5/4/22 |
| Heuchera americana var. calycosa | Heuchera americana var. americana | Oak mountain park road | 34 | 33 | 0.5074626866 | 5/4/22 |
| Heuchera americana var. heteradenia | Heuchera americana var. americana | A55 | 66 | 9 | 0.88 | 5/10/22 |
| Heuchera americana var. americana | Heuchera americana var. americana | A57 | 34 | 1 | 0.9714285714 | 5/11/22 |
| Heuchera americana var. americana | Heuchera americana var. americana | A58 | 10 | 0 | 1 | 5/11/22 |
| Heuchera americana var. americana | Heuchera americana var. americana | A60 | 47 | 1 | 0.9791666667 | 5/13/22 |
| Heuchera americana var. heteradenia | Heuchera americana var. americana | A62 | 15 | 9 | 0.625 | 5/14/22 |
| Heuchera americana var. heteradenia | Heuchera americana var. americana | A63 | 35 | 3 | 0.9210526316 | 5/15/22 |

| | | | | | | |
|---|---|---|---|---|---|---|
| Heuchera americana var. hirsuticaulis | Heuchera americana var. hirsuticaulis | Sardis Lake | 0 | 50 | 0 | 1 May 2021 |
| Heuchera americana var. hirsuticaulis | Heuchera americana var. hirsuticaulis | Lake Wilson AR | 0 | 5 | 0 | 15 May 2021 |
| Heuchera americana var. hirsuticaulis | Heuchera americana var. hirsuticaulis | Onda Mountain Road | 16 | 0 | 1 | 15 May 2021 |
| Heuchera americana var. hirsuticaulis | Heuchera americana var. hirsuticaulis | Taum Sauk | 78 | 115 | 0.4041450777 | 6/2/22 |
| Heuchera richardsonii var. affinis | Heuchera americana var. hirsuticaulis | camper spring/ meramec river | 36 | 23 | 0.6101694915 | 6/2/22 |
| Heuchera richardsonii var. affinis | Heuchera americana var. hirsuticaulis | A44 | 28 | 23 | 0.5490196078 | 6/8/22 |
| Heuchera longiflora var. aceroides | Heuchera longiflora var. aceroides | H89 | 12 | 4 | 0.75 | 5/14/22 |
| Heuchera richardsonii var. affinis | Heuchera richardsonii | H167 Starved rock pool | 5 | 1 | 0.8333333333 | 6/4/22 |
| Heuchera richardsonii var. affinis | Heuchera richardsonii | Stone Barn Savanna | 20 | 19 | 0.5128205128 | 6/4/22 |
| Heuchera richardsonii var. grayana | Heuchera richardsonii | A16 Devil's Lake | 23 | 8 | 0.7419354839 | 6/5/22 |
